# Genomic clustering and sequence context of mutations in human monkeypox virus (hMPXV1) genomes

**DOI:** 10.1101/2022.11.21.517357

**Authors:** Diego Forni, Rachele Cagliani, Uberto Pozzoli, Manuela Sironi

## Abstract

The ongoing worldwide monkeypox outbreak is caused by viral lineages (globally referred to as hMPXV1) that are related to but distinct from clade IIb MPXV viruses transmitted in Nigeria. Analysis of genetic differences indicated that APOBEC-mediated editing might be responsible for the unexpectedly high number of mutations observed in hMPXV1 genomes. Here, using 1624 hMPXV1 publicly available sequences, we analyzed mutations that accrued since 2017 until the emergence of the current predominant variant (B.1), as well as those that that have been accumulating during the 2022 outbreak. We found that substitutions tend to cluster and mutational hot-spots are observed.

Investigation of the sequence context of C to T changes indicated a preference for 5’-TCA/G-3’ motifs, suggesting APOBEC3F- or APOBEC3A-mediated editing. The sequence context has remained unchanged since 2017, indicating that the same mutational mechanism that is driving the accumulation of substitutions during the ongoing human-to-human transmission, was already operating before the virus left Africa. We suggest that APOBEC3A is the most likely candidate, given its expression in the skin and its known role in the editing of human papillomavirus.

## Introduction

Monkeypox is caused by monkeypox virus (MPXV), a member of the *Poxviridae* family. Until recently, the disease was considered a rare zoonosis occasionally transmitted in an endemic area that ranges from West to Central Africa. In the last five years, though, the prevalence of monkeypox has been increasing, both in Africa and worldwide (1). In particular, since the beginning of May 2022, a multi-country outbreak has caused more than 79400 cases in 110 countries (as of November 16^th^). The epidemiology of the ongoing outbreak is distinct from that previously observed in Africa: viral spread is sustained by human-to-human transmission, often mediated by sexual contact (1). Because of distinctive epidemiological and genomic characteristics, the virus causing the outbreak was re-named hMPXV1 (2).

Genomic surveillance of hMPXV1 showed that sequences sampled in 2022, as well as a few sampled in the US in 2021, form a so-called A lineage. Such lineage also includes a few Nigerian strains dating between 2017 and 2019 and is phylogenetically related to MPXV clade IIb. (Fig. 1A) (3, 4). Within the A lineage, a predominant B.1 lineage accounts for the majority of 2022 cases (Fig. 1A). The A and B.1 lineages are characterized by several single nucleotide substitutions, most of which involve GA>AA or TC >TT replacements (3, 4). This led to the suggestion that host apolipoprotein B mRNA editing catalytic polypeptide-like 3 (APOBEC3) enzymes have been driving the evolution of hMPXV1 since 2017, and possibly earlier (3, 4).

**Figure 1.**
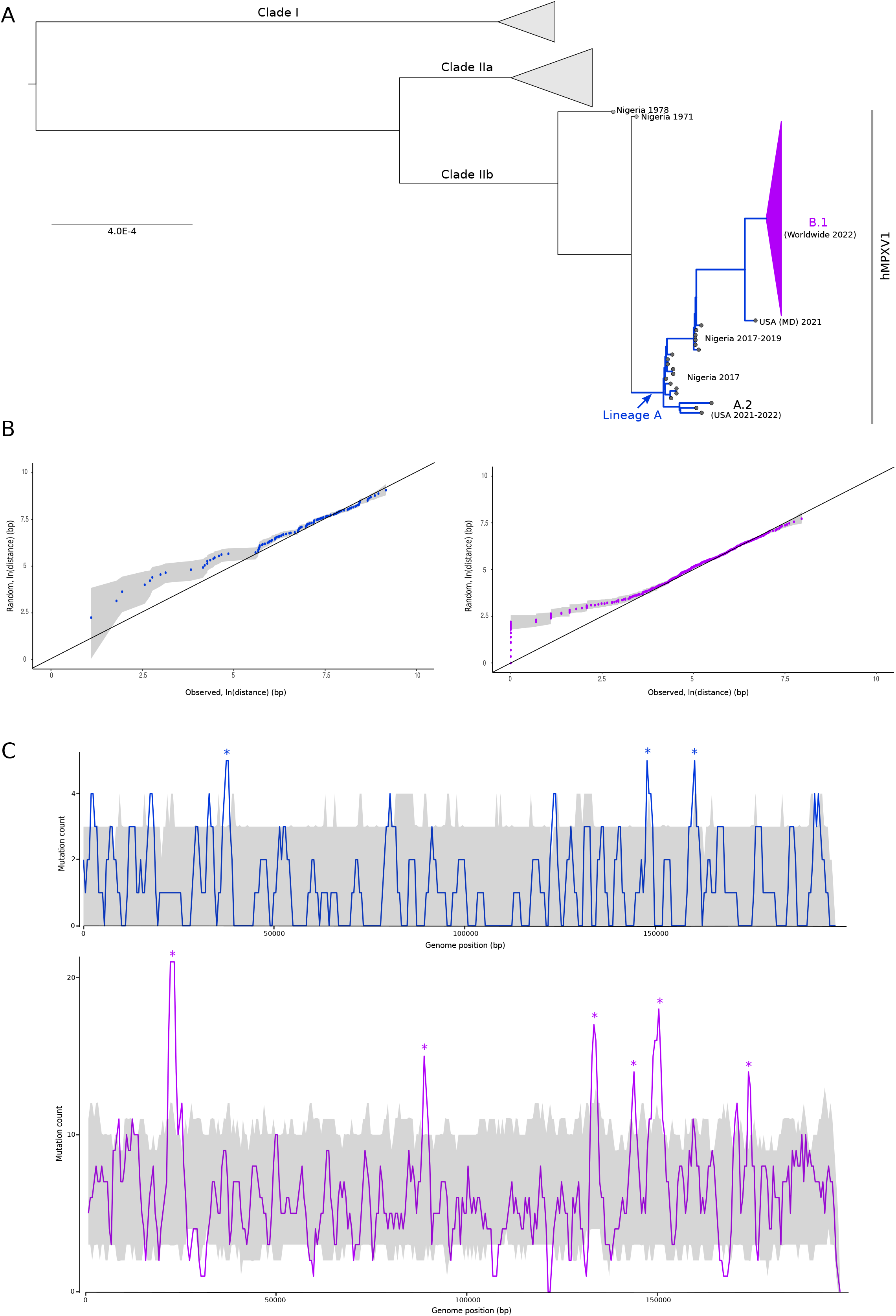
Clustering of mutations in hMPXV1 genomes. **(A)** Phylogenetic tree of 80 representative MPXV/hMPXV1 strains. A conserved genomic region (16) was used and the tree was generated using IQTREE. Blue branches are the ones on which we counted mutations that accumulated in the A lineage since 2017 until the emergence of the current predominant outbreak variant. The collapsed magenta branches contributed mutations that have been accumulating during the 2022 outbreak. **(B)** Quantile-quantile plots of mutation distances in lineages A (left, blue) and B.1 (right, magenta). Observed distances were plotted against distances in 100 random genomes. Dots represent the median, the gray shadow represents the 5^th^ and 95^th^ percentiles. **(C)** Sliding-window analysis of mutation counts. Mutations in lineages A (top, blue) and B.1 (bottom, magenta) were counted in windows of 2000 bp, moving with a step of 500 bp. The gray shadows represent the 5^th^ and 95^th^ percentiles of mutation counts in random genomes. Asterisks denote mutation hot-spots.

## Results and discussion

The MPXV/hMPXV1 genome is a ∼195 kb-long dsDNA molecule, with a GC content of about 33%. The mutations that characterize the A and B.1 lineages tend to be distributed along the genome (3, 4). However, APOBEC-generated mutations might be expected to cluster (5). We reasoned that some level of clustering was possibly missed because of the low GC content of the hMPXV1 genome and due to the relatively small number of mutations analyzed in previous studies. To determine whether this is the case, we retrieved hMPXV1 genomes available in public repositories and we counted the number of mutations that accrued in the A lineage since 2017 until the emergence of the current predominant outbreak variant B.1 (n=121) (i.e., using the Nigeria-SE-1971 strain as reference), as well as mutations (n=620) that have been accumulating during the 2022 outbreak (i.e., using the USA, MD 2021 strain as reference) (Fig. 1A). As expected, the majority of these were G>A or C>T changes (83.5% in lineage A, 84.2% in lineage B.1). As an empirical comparison and to account for local differences in GC content, for both sets of mutations, we generated 100 mutated genomes by randomly changing the same overall number of C, T, G, and A nucleotides as in the real genomes. These are hereafter referred to as random genomes. We next calculated the distance between consecutive mutations in real and random genomes. Quantile-quantile plots of distances in the real genomes against all 100 random genomes indicated that the observed mutations are significantly closer than the random ones (Fig. 1B).

To determine whether and which specific regions of hMPXV1 genomes represent mutation hot-spots, we counted mutations in sliding windows. The same procedure was also performed for random genomes and the 5^th^ and 95^th^ percentiles of mutation counts were calculated. Mutation hot-spots were observed for both A and B1 lineages, with limited overlap (Fig. 1C). This confirms the results obtained by calculating distances and indicates that substitutions are non-randomly distributed and tend to cluster in specific regions. Mutation hot-spots show limited overlap between the two lineages and involve genes with diverse functions (Fig. 2A).

**Figure 2.**
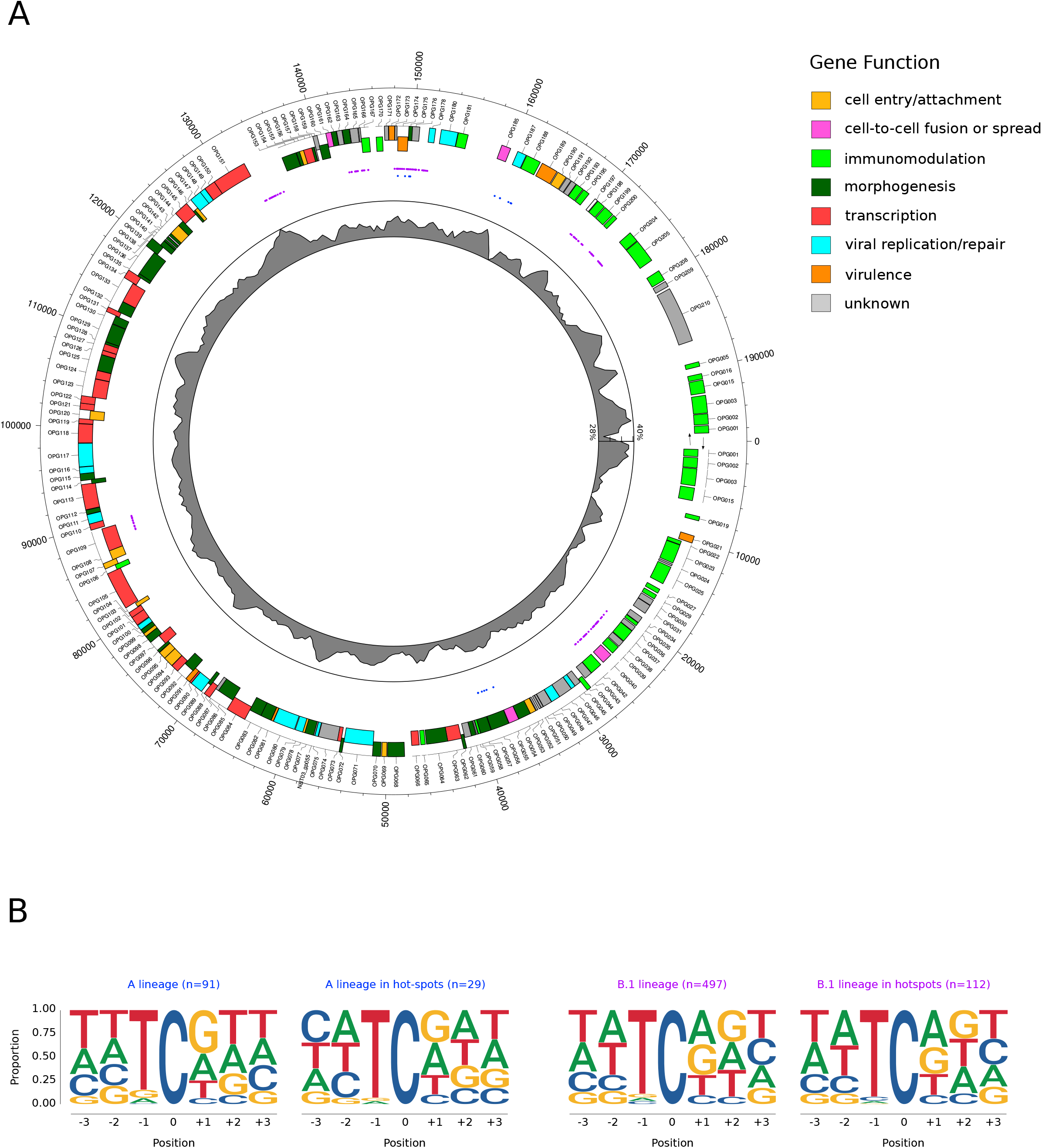
Context of mutations in hMPXV1 genomes. **(A)** Circos plot of the hMPXV1 genome. Positions and annotation refers to the MPXV-M5312_HM12_Rivers strain. Shown are (from the outside to the inside): genes color-coded according to function (1) and to sense of transcription, C>T/G>A mutations within hot-spots (see figure 1), GC content. **(B)** Sequence context in which cytidine mutations occur. Letter size represents the relative frequency of each base flanking mutated cytidines.

We next set out to analyze the sequence context of the observed substitutions. Because these are hypothesized to derive from the action of one or more APOBEC enzymes (i.e., from the deamination of cytosines), G to A mutations were considered C to T changes in the opposite strand. Also, the counts of bases flanking the mutated cytosines were normalized by the frequency of each nucleotide in the hMPXV1 genome. Results confirmed a very strong preference for a T in the -1 position, but also indicated a preference, albeit less marked, for G or A in the +1 position (Fig. 2B). This was observed for both lineage A and lineage B.1, irrespective of the fact that mutations are located or not within the hot-spots. This indicates that, whatever mechanism underlies mutation occurrence, it preferentially, but not exclusively, targets specific genomic regions. Also, this observation strongly supports the view that the mutation process that is driving the accumulation of substitutions during the ongoing human-to-human transmission, was already operating in 2017, before the virus left Africa.

The observed sequence context is consistent with editing being mediated by APOBEC3F (6) or APOBEC3A (7, 8). Both enzymes are present in the cytoplasm (or shuffle between the nucleus and the cytoplasm) (5), where poxvirus replication occurs. The activity of APOBEC3F has been mainly investigated in the context of HIV infection, whereas APOBEC3A has been intensely studied for its role as a mutagen in cancer genomes (5). However, APOBEC3A was also implicated in the editing and restriction of human papillomavirus (HPV), as well as in the deamination of foreign dsDNA (9-11). Also, this enzyme is expressed in skin and mucosal tissues, making it a likely candidate for introducing hMPXV1 mutations (10).

In cancer genomes, there is little evidence for transcriptional asymmetry of APOBEC3A-induced mutations (8). Conversely, replication asymmetry is observed, with most mutations occurring on the lagging strand (8). APOBEC editing of human papillomavirus and of plasmid DNA instead occurs on both strands (9, 11). We thus checked for asymmetries in C>T/A>G changes in hMPXV1 using mutations accumulating in lineage B.1 (because they are more numerous). We observed very similar proportions of TC>TT mutations on both strands (51.6% on the + strand and 48.4% on the - strand).

Assuming that, in analogy to vaccinia virus (VACV), MPXV has one single origin of replication at one end of the genome (12), this implies no replication asymmetry of mutations. Likewise, we found no evidence of transcriptional asymmetry (genes transcribed from the – strand: 112 C>T and 99 G>A; genes transcribed from the + strand: 105 C>T and 116 G>A). Thus, possibly because of distinct replication systems, the effects of APOBEC3A editing seem to differ depending on the target (genomic DNA or virus/plasmid).

Overall, these observations lend further support to the hypothesis that APOBEC3 enzymes, and specifically APOBEC3A, are responsible for introducing mutations in hMPXV genomes. As previously noted, sustained circulation in the human population might have exposed the virus to a different APOBEC3 repertoire than the one of its natural host (most likely African rodents) (3, 4). In fact, primates encode an expanded array of APOBEC3 enzymes compared to other mammals (13). Also, it possible that the change in the route of transmission, which occurs mainly via sexual contact rather than areosolization, resulted in the contact of hMPXV1 with different cell/tissue environments (e.g., the skin and anogenital mucosal surfaces), and, consequently, with distinct innate immunity systems. In this respect, it is worth noting that HPV, which is also characterized by sexual transmission and infects skin keratinocytes, is edited by APOBEC3A. The notion that poxviruses are not targets of APOBEC3-mediated editing derives from the observation that APOBEC3G, 3F or 3H have no effect on VACV replication (14), possibly because the poxvirus replication complex is sequestered in specialized regions of the cytoplasm known as virus factories (13). Nonetheless, it is possible that APOBEC3 over-expression did induce some sub-lethal editing in VACV genomes, which were not sequenced. Indeed, if APOBECs are responsible for mutations in hMPXV1 genomes, they are clearly insufficient to curb viral replication and spread. In this respect, it is also worth noting that we analyzed all observed mutations, without any consideration of their effect. However, mutations that severely affect viral fitness are eliminated by natural selection and are thus under-represented. In this sense, the analysis of mutations that are accumulating in the outbreak predominant population are more reflective of the mutation process than those that accumulated since 2017 in lineage A (because in this case natural selection acted over a longer period).

## Materials and Methods

### Sequences, alignment and phylogenetic tree

hMPXV1 sequences were retrieved from the GISAID Initiative (https://www.gisaid.org) and from the National Center for Biotechnology Information (NCBI) databases (as of October, 20^th^, 2022). In particular, 1603 complete, high coverage, lineage B.1 genomes were retrieved from GISAID and 21 complete/almost complete lineage A genomes were downloaded from NCBI nucleotide database. Strain lists are provided as supplementary dataset 1 and 2. We used as references the Nigeria-SE-1971 (NCBI Accession id: KJ642617) and the MPXV_USA_2021_MD (NCBI Accession ID: ON676708) strains for lineages A and B.1, respectively. Whole genome alignments for the two lineages and the corresponding references were generated using the MAFFT software (v7.427) (15) with default parameters. Substitution were counted by comparing all positions of the lineage alignment with the corresponding reference sequence and, to avoid sequencing errors, we considered mismatches that occurred in at least two sequences. Nucleotide positions were then converted and referred to the reference genomes.

Eighty representative strains were selected to generate a MPXV/hMPXV1 phylogeny. The conserved genomic region (16) was aligned using MAFFT and the tree was generated using IQTREE (17).

### Mutation distribution

To compare mutation distributions, we generated two set of 100 random genomes using the two reference genomes as scaffolds. In particular, for each reference genome, we mutated the same number of nucleotides that we found mutated in the B.1 or in the A lineage, by randomly choosing from all genomic position of the same bases as those where mutations were observed (e.g., we observed 283 mutated cytosines in the B.1 lineage and we mutated the same number of cytosines in each random genome). We then calculated the distance between consecutive mutations for each of the 100 random sequences. To asses whether distances among observed mutations are not random, the observed distribution distances were compared with each of the 100 random distributions using quantile-quantile plot.

The distribution of mutations was also analyzed along the hMPXV1 genome using sliding windows of 2000 nucleotides in size and 500 nucleotide steps. We then counted the number of mutations falling within each window. Again, we performed the same analysis for the random genomes and we compared the number of observed mutations with the distribution from random genomes within the same windows. To be conservative, we considered mutation hot spots only those windows in which the count of observed mutation was higher the maximum value reached by the 95^th^ percentiles in random counts along the whole genome. All analyses were performed in R enviroment (18).

### APOBEC mutation context

C to T and G to A changes were analyzed in the context of APOBEC enzyme activity. Therefore, G to A changes were considered C to T changes occurring in the opposite strand, and the reverse complement sequences were analyzed. In particular, for both lineages, we analyzed C to T changes both found along the whole genome and in mutation hot spots. We retrieved +3 and -3 nucleotides flanking these mutations. Mutations falling in the ITRs were counted only once to avoid redundancy. The relative frequencies at each position were than used to generate sequence logos using the R package ggseqlogo (19).

## Acknowledgments

We gratefully acknowledge the authors, originating and submitting laboratories of the sequences from GISAID’s EpiPox™ database on which this research is based.

