## Supplementary dataset 1 for "Genomic clustering and sequence context of mutations in human monkeypox virus (hMPXV1) genomes"

We gratefully acknowledge the following Authors from the Originating laboratories responsible for obtaining the specimens, as well as the Submitting laboratories where the genome data were generated and shared via GISAID, on which this research is based.

All Submitters of data may be contacted directly via www.gisaid.org

Authors are sorted alphabetically.

| Accession ID | Originating Laboratory | Submitting Laboratory | Authors |
|---|---|---|---|
| EPI_ISL_13052263 | Microbiol Genomics and Bioinformatics, Bundeswehr Institute of Microbiology | Microbiol Genomics and Bioinformatics, Bundeswehr Institute of Microbiology | Antwerpen,M.H., Lang,D., Zange,S., Walter,M.C. and Woelfel,R. |
| EPI_ISL_13052266, EPI_ISL_13052267, EPI_ISL_13052268, EPI_ISL_13052269, EPI_ISL_13052270, EPI_ISL_13052272, EPI_ISL_13052273 | Instituto Nacional de Saude Doutor Ricardo Jorge (INSA) | Instituto Nacional de Saude Doutor Ricardo Jorge (INSA) | Joana Isidro, Vítor Borges, Miguel Pinto, Daniel Sobral, João Dourado Santos, Alexandra Nunes, Verónica Mixão, Rita Ferreira, Daniela Santos, Silvia Duarte, Luís Vieira, Maria José Borrego, Sofia Núncio, Isabel Lopes de Carvalho, Ana Pelerito, Rita Cordeiro, João Paulo Gomes |
| EPI_ISL_13052274 | Laboratory of Virology, University Hospitals of Geneva | Laboratory of Virology, University Hospitals of Geneva | Laubscher,F., Chudzinski,V., Schibler,M., Kaiser,L. and Renzoni,A. |
| EPI_ISL_13052278 | Research and Evaluation, UKHSA | Research and Evaluation, UKHSA | Osman,K.L., Lewandowski,K.S., Pullan,S.T., Carter,D.P., Crook,J.M., Vipond,R. and Chand,M. |
| EPI_ISL_13052279, EPI_ISL_13052280, EPI_ISL_13052281 | Research and Evaluation, UKHSA | Research and Evaluation, UKHSA | Osman,K.L., Lewandowski,K.S., Carter,D.P., Crook,J.M., Pullan,S.T., Vipond,R. and Chand,M. |
| EPI_ISL_13052282 | Microbiology, Immunology and Transplantation, KU Leuven, Rega Institute | Microbiology, Immunology and Transplantation, KU Leuven, Rega Institute | Vanmechelen,B., Wawina-Bokalanga,T., Logist,A.-S., Sinnesael,R., Ysebaert,L., Verlinden,J., Bloemen,M. and Maes,P. |
| EPI_ISL_13052283 | Microbiology, Immunology and Transplantation, KU Leuven, Rega Institute | Microbiology, Immunology and Transplantation, KU Leuven, Rega Institute | Wawina-Bokalanga,T., Vanmechelen,B., Logist,A.-S., Sinnesael,R., Ysebaert,L., Verlinden,J., Bloemen,M. and Maes,P. |
| EPI_ISL_13052285 | Laboratory of Virology, University Hospitals of Geneva | Laboratory of Virology, University Hospitals of Geneva | Laubscher,F., Schibler,M., Kaiser,L. and Renzoni,A. |
| EPI_ISL_13052287 | Virology, GENomique EPIdemiologique des maladies Infectieuses | Virology, GENomique EPIdemiologique des maladies Infectieuses | unknown |
| EPI_ISL_13052288 | Department of Health, Utah Public Health Laboratory | Department of Health, Utah Public Health Laboratory | Young,E.L., Hergert,J. and Oakeson,K.F. |
| EPI_ISL_13052290 | Laboratory for Diagnostics of Zoonoses and WHO Centre, Institute of Microbiology and Immunology, Faculty of Medicine, University of Ljubljana | Laboratory for Diagnostics of Zoonoses and WHO Centre, Institute of Microbiology and Immunology, Faculty of Medicine, University of Ljubljana | Zakotnik,S., Vlaj,D., Suljic,A., Zorec,T.M., Korva,M., Poljak,M. and Avsic Zupanc,T. |
| EPI_ISL_13052291 | Laboratory for Diagnostics of Zoonoses and WHO Centre, Institute of Microbiology and Immunology, Faculty of Medicine, University of Ljubljana | Laboratory for Diagnostics of Zoonoses and WHO Centre, Institute of Microbiology and Immunology, Faculty of Medicine, University of Ljubljana | Zakotnik,S., Vlaj,D., Suljic,A., Zorec,T.M., Skubic,C., Rozman,D., Korva,M., Poljak,M. and Avsic Zupanc,T. |
| EPI_ISL_13052292 | Victorian Infectious Diseases Reference Laboratory, Doherty Institute | Victorian Infectious Diseases Reference Laboratory, Doherty Institute | Hammerschlag,Y., MacLeod,G., Papadakis,G., Adan-Sanchez,A., Druce,J.D., Williamson,D.A., Cheng,A.C. and McMahon,J.H. |
| EPI_ISL_13052295 | SC (UCO) Igiene e Sanità Pubblica, ASUGI, Trieste | Genomics and Epigenomics, AREA Science Park | Licastro,D., DeGasperi,M., Negri,C., Piscianz,E., Koncan,R., Dal Monego,S., Segat,L. and D'Agaro,P. |
| EPI_ISL_13056892, EPI_ISL_13056893, EPI_ISL_13056896, EPI_ISL_13056897, EPI_ISL_13056901, EPI_ISL_13056902, EPI_ISL_13056903, EPI_ISL_13056904, EPI_ISL_13056905, EPI_ISL_13056906, EPI_ISL_13056907, EPI_ISL_13056908, EPI_ISL_13056909 | | | |
| see above | Instituto Nacional de Saude Doutor Ricardo Jorge (INSA) | Instituto Nacional de Saude Doutor Ricardo Jorge (INSA) | Joana Isidro, Vítor Borges, Miguel Pinto, Daniel Sobral, João Dourado Santos, Alexandra Nunes, Verónica Mixão, Rita Ferreira, Daniela Santos, Silvia Duarte, Luís Vieira, Maria José Borrego, Sofia Núncio, Isabel Lopes de Carvalho, Ana Pelerito, Rita Cordeiro, João Paulo Gomes |
| EPI_ISL_13056910 | Biochemistry and Molecular Genetics, Israel Institute for Biological Research | Biochemistry and Molecular Genetics, Israel Institute for Biological Research | Israeli,O., Guedj-Dana,Y., Lazar,S., Shifman,O., Erez,N., Weiss,S., Paran,N., Israely,T., Schuster,O., Zvi,A., Beth-Din,A. and Cohen Gihon,I. |
| EPI_ISL_13100618 | Centers for Disease Control & Prevention (CDC), Division of High Consequence Pathogens and Pathology (DHCPP-PRB) | Centers for Disease Control & Prevention (CDC), Division of High Consequence Pathogens and Pathology (DHCPP-PRB) | Gigante,C.M., Ventura,J., Seabolt,M.H., Wilkins,K., McCollum,A., Hutson,C., Davidson,W., Rao,A., Nash,J. and Li,Y. |
| EPI_ISL_13100619 | Centers for Disease Control & Prevention (CDC), Division of High Consequence Pathogens and Pathology (DHCPP-PRB) | Centers for Disease Control & Prevention (CDC), Division of High Consequence Pathogens and Pathology (DHCPP-PRB) | Gigante,C.M., Lee,P., Seabolt,M.H., Wilkins,K., McCollum,A., Hutson,C., Davidson,W., Rao,A., Mendoza,R. and Li,Y. |
| EPI_ISL_13100620 | Centers for Disease Control & Prevention (CDC), Division of High Consequence Pathogens and Pathology (DHCPP-PRB) | Centers for Disease Control & Prevention (CDC), Division of High Consequence Pathogens and Pathology (DHCPP-PRB) | Gigante,C.M., Atkinson,A., Seabolt,M.H., Wilkins,K., McCollum,A., Hutson,C., Davidson,W., Rao,A., Murray,J. and Li,Y. |
| EPI_ISL_13100719 | Centers for Disease Control & Prevention (CDC), Division of High Consequence Pathogens and Pathology (DHCPP-PRB) | Centers for Disease Control & Prevention (CDC), Division of High Consequence Pathogens and Pathology (DHCPP-PRB) | Gigante,C.M., Ventura,J., Seabolt,M.H., Wilkins,K., McCollum,A., Hutson,C., Davidson,W., Rao,A., Nash,J. and Li,Y. |
| EPI_ISL_13117291 | Centre for Biological Threats, Highly Pathogenic Viruses, Robert Koch Institute | Centre for Biological Threats, Highly Pathogenic Viruses, Robert Koch Institute | Brinkmann,A., Kohl,C., Uddin,S., Pape,K., Schrick,L., Michel,J., Jessen,H., Schaade,L. and Michel,A. |
| EPI_ISL_13117292, EPI_ISL_13117293 | Centre for Biological Threats, Highly Pathogenic Viruses, Robert Koch Institute | Centre for Biological Threats, Highly Pathogenic Viruses, Robert Koch Institute | Brinkmann,A., Kohl,C., Uddin,S., Pape,K., Schrick,L., Michel,J., Stocker,H., Schaade,L. and Nitsche,A. |
| EPI_ISL_13117294, EPI_ISL_13117295, EPI_ISL_13117296, EPI_ISL_13117297, EPI_ISL_13117298 | Centre for Biological Threats, Highly Pathogenic Viruses, Robert Koch Institute | Centre for Biological Threats, Highly Pathogenic Viruses, Robert Koch Institute | Brinkmann,A., Kohl,C., Uddin,S., Pape,K., Schrick,L., Michel,J., Schaade,L. and Nitsche,A. |
| EPI_ISL_13148263, EPI_ISL_13148264, EPI_ISL_13148265 | Centre for Biological Threats, Highly Pathogenic Viruses, Robert Koch Institute | Centre for Biological Threats, Highly Pathogenic Viruses, Robert Koch Institute | Brinkmann,A., Kohl,C., Uddin,S., Pape,K., Schrick,L., Michel,J., Jessen,H., Schaade,L. and Nitsche,A. |
| EPI_ISL_13148266, EPI_ISL_13148267, EPI_ISL_13148268, EPI_ISL_13148269 | Centre for Biological Threats, Highly Pathogenic Viruses, Robert Koch Institute | Centre for Biological Threats, Highly Pathogenic Viruses, Robert Koch Institute | Brinkmann,A., Kohl,C., Uddin,S., Pape,K., Schrick,L., Michel,J., Stocker,H., Schaade,L. and Nitsche,A. |
| EPI_ISL_13148270, EPI_ISL_13148271, EPI_ISL_13148272, EPI_ISL_13148273, EPI_ISL_13148274, EPI_ISL_13148275, EPI_ISL_13148276 | Centre for Biological Threats, Highly Pathogenic Viruses, Robert Koch Institute | Centre for Biological Threats, Highly Pathogenic Viruses, Robert Koch Institute | Brinkmann,A., Kohl,C., Uddin,S., Pape,K., Schrick,L., Michel,J., Schaade,L. and Nitsche,A. |
| EPI_ISL_13194516 | Alberta Precision Laboratories | Alberta Precision Laboratories | Matthew Croxen, Ashwin Deo, Paul Dieu, Xiaoli Dong, Kara Gill, David Granger, Christina Ferrato, Vanipriyadarsini Ikkurti, Jamil Kanji, Petya Koleva, Vincent Li, Colin Lloyd, Tarah Lynch, Raymond Ma, Kanti Pabbaraju, Silas Rotich, Hilary Sergeant, Steven Shideler, Todd Skitsko, Sandy Shokopoles, Graham Tipples, Johanna Thayer, Anita Wong |
| EPI_ISL_13234112 | Laboratório Central de Saúde Pública do Estado do Rio Grande do Sul | Instituto Adolfo Lutz Strategic Laboratory | Claudio Tavares Sacchi, Karoline Rodrigues Campos, Adriano Abbud, Adriana Bugno |
| EPI_ISL_13244349 | Erasmus Medical Center Department of Virology | Erasmus Medical Center Department of Virology | Bas Oude Munnink, Marjan Boter, Babette Weller, Richard Molenkamp, Janette Rahamat-Langendoen, Reina Sikkema, Marion Koopmans |

| | | | |
|---|---|---|---|
| EPI_ISL_13251120 | Laboratory of Virology, INMI Lazzaro Spallanzani IRCCS | Laboratory of Virology, INMI Lazzaro Spallanzani IRCCS | Giombini,E., Gruber,C.E.M., Rueca,M., Gramigna,G., VIta,S., Carletti,F., D'Abramo,A., Lapa,D., Puro,V., Fabeni,L., Butera,O., Colavita,F., Meschi,S., Matusali,G., Specchiarello,E., Vairo,F., Vaia,F., Garbuglia,A.R., Nicastri,E., Antinori,A., GIrardi,E. and Maggi,F. |
| EPI_ISL_13251157, EPI_ISL_13251723 | checkin Zollhaus | Institute of Medical Virology, University of Zurich | Verena Kufner, Gabriela Ziltener, Maryam Zaheri, Stefan Schmutz, Annette Audigé, Odette Bernasconi, Kevin Steiner, Jon Huder, Cyril Shah, Riccarda Capaul, Guido Bloemberg, Jürg Böni, Michael Huber, Alexandra Trkola |
| EPI_ISL_13269478 | Alberta Precision Laboratories | Alberta Precision Laboratories | Matthew Croxen, Ashwin Deo, Paul Dieu, Xiaoli Dong, Kara Gill, David Granger, Christina Ferrato, Vanipriyadarsini Ikkurti, Jamil Kanji, Petya Koleva, Vincent Li, Colin Lloyd, Tarah Lynch, Raymond Ma, Kanti Pabbaraju, Silas Rotich, Hilary Sergeant, Steven Shideler, Todd Skitsko, Sandy Shokopoles, Graham Tipples, Johanna Thayer, Anita Wong |
| EPI_ISL_13302316 | Laboratory of Clinical Microbiology, Virology and Bioemergencies. ASST-Fatebenefratelli-Sacco, L.Sacco University Hospital | Army Medical and Veterinary Research Center | Silvia Fillo, Riccardo De Sanctis, Giovanni Faggioni, Andrea Ciammaruconi, Anna Anselmo, Vanessa Vera Fain, Simone Di Sabatino, Francesco Giordani, Antonella Fortunato, Rossella Brandi, Giulia Campoli, Marzia Cavalli, Anella Monte, Martina Lipari, Maria Di Spirito, Giorgia Grilli, Silvia Chimienti, Giandomenico Cerreto, Filippo Molinari, Giancarlo Petralito, Davide Mileto, Valeria Micheli, Maria Rita Gismondo, Florigio Lista |
| EPI_ISL_13308117, EPI_ISL_13308118, EPI_ISL_13308119, EPI_ISL_13308121, EPI_ISL_13308122, EPI_ISL_13308124, EPI_ISL_13308125, EPI_ISL_13308127, EPI_ISL_13308129, EPI_ISL_13308131, EPI_ISL_13308133, EPI_ISL_13308135, EPI_ISL_13308137, EPI_ISL_13308139, EPI_ISL_13308140, EPI_ISL_13308142, EPI_ISL_13308144, EPI_ISL_13308145, EPI_ISL_13308146, EPI_ISL_13308147, EPI_ISL_13308148, EPI_ISL_13308150, EPI_ISL_13308151, EPI_ISL_13308153, EPI_ISL_13308155, EPI_ISL_13308157 | | | |
| see above | Centre for Biological Threats, Highly Pathogenic Viruses, Robert Koch Institute | Centre for Biological Threats, Highly Pathogenic Viruses, Robert Koch Institute | Brinkmann,A., Kohl,C., Uddin,S., Pape,K., Schrick,L., Michel,J., Schaade,L. and Nitsche,A. |
| EPI_ISL_13308158, EPI_ISL_13308160 | IRBA Research Institute Biomédicale Des Armées | IRBA Research Institute Biomédicale Des Armées | Jarjaval,F., Nolent,F., Criqui,A., Chapus,C., Lamer,O., Ferraris,O. and Gorge,O. |
| EPI_ISL_13308163, EPI_ISL_13308167 | Laboratory for Diagnostics of Zoonoses and WHO Centre, Institute of Microbiology and Immunology, Faculty of Medicine, University of Ljubljana | Laboratory for Diagnostics of Zoonoses and WHO Centre, Institute of Microbiology and Immunology, Faculty of Medicine, University of Ljubljana | Zakotnik,S., Vlaj,D., Suljic,A., Zorec,T.M., Korva,M., Poljak,M. and Avsic Zupanc,T. |
| EPI_ISL_13314740 | Laboratorio de Vigilancia em Saude de Vinhedo | Instituto Adolfo Lutz Strategic Laboratory | Claudio Tavares Sacchi, Karoline Rodrigues Campos, Adriano Abbud, Adriana Bugno |
| EPI_ISL_13331713 | Laboratory of Virology, INMI Lazzaro Spallanzani IRCCS | Laboratory of Virology, INMI Lazzaro Spallanzani IRCCS | Gramigna,G., Giombini,E., Gruber,C.E.M., Rueca,M., Carletti,F., Cicalini,S., Lapa,D., Puro,V., Marani,A., Fabeni,L., Butera,O., Colavita,F., Meschi,S., Matusali,G., Rivano Capparuccia,M., Specchiarello,E., Vairo,F., Vaia,F., Nicastri,E., Antinori,A., Girardi,E. and Maggi,F. |
| EPI_ISL_13331714 | Department of Virology, Faculty of Medicine, University of Helsinki, Hartmaninkatu 3 | Department of Virology, Faculty of Medicine, University of Helsinki, Hartmaninkatu 3 | Kant,R., Smura,T., Vauhkonen,H. and Vapalahti,O. |
| EPI_ISL_13331715 | Department of Virology, Faculty of Medicine, University of Helsinki, Hartmaninkatu 3 | Department of Virology, Faculty of Medicine, University of Helsinki, Hartmaninkatu 3 | Kant,R., Smura,T., Vauhkonen,H., Vapalahti,O. and Sironen,T. |
| EPI_ISL_13331717 | Genomics Division, Instituto Tecnologico y de Energias Renovables (ITER), Poligono Industrial de Granadilla | Genomics Division, Instituto Tecnologico y de Energias Renovables (ITER), Poligono Industrial de Granadilla, | Alcoba-Florez,J., Munoz-Barrera,A., Ciuffreda,L., Rodriguez-Perez,H., Rubio-Rodriguez,L.A., Gil-Campesino,H., Garcia-Martinez de Artola,D., Inigo-Campos,A., Diez-Gil,O., Gonzalez-Montelongo,R., Valenzuela-Fernandez,A., Lorenzo-Salazar,J.M. and Flores,C. |
| EPI_ISL_13339105 | Microbiology Service, Hospital Universitario Clinico San Cecilio, Granada | Microbiology Service, Hospital Universitario Clinico San Cecilio, Granada | Chueca N, de Salazar A, Viñuela L, Fuentes A, Casimiro-Soriguer CS, Perez-Florido J, Dopazo J, Garcia F |
| EPI_ISL_13343634 | Instituto de Infectologia Emilio Ribas | Instituto Adolfo Lutz Strategic Laboratory | Claudio Tavares Sacchi, Karoline Rodrigues Campos, Adriano Abbud, Adriana Bugno |
| EPI_ISL_13343697 | Fleury Medicina Dignóstica | Instituto Adolfo Lutz Strategic Laboratory | Claudio Tavares Sacchi, Karoline Rodrigues Campos, Adriano Abbud, Adriana Bugno |
| EPI_ISL_13343718 | Hospital Santa Ignes | Instituto Adolfo Lutz Strategic Laboratory | Claudio Tavares Sacchi, Karoline Rodrigues Campos, Adriano Abbud, Adriana Bugno |
| EPI_ISL_13351002 | B.C. Centre for Disease Control Public Health Laboratory | B.C. Centre for Disease Control Public Health Laboratory | John Tyson, Tracy Lee, Anthea Lam, Josh Quick, Agatha Jassem, Natalie Prystajecky, Linda Hoang, Inna Sekirov, Catherine Hogn, Frankie Tsang, Mel Krajden |
| EPI_ISL_13362760 | Laboratorio di Epidemiologia Molecolare e Sanità Pubblica-Policlinico Bari | Istituto Zooprofilattico Sperimentale della Puglia e della Basilicata | Parisi A, Simone D, Capozzi L, Del Sambro L, Bianco A, Chironna M, Loconsole D, Sallustio F, Galante D, Pace L, Manzulli V, Fasanella A. |
| EPI_ISL_13363142 | Hospital Universitari Vall d'Hebron | Hospital Universitari Vall d'Hebron | Maria Piñana, Cristina Andrés, Alejandra González-Sánchez, Damir Garcia-Cehic, Ariadna Rando, Juliana Esperalba, Maria Gema Codina, Maria Carmen Martin, Carla Castillo, Karen García, Rodrigo Vásquez, Maria Piquer, Tomàs Pumarola, Josep Quer, Andrés Antón |
| EPI_ISL_13374487 | National Public Health Center, National Biosafety Laboratory | National Public Health Center, National Biosafety Laboratory | Judit Henczkó, Dániel Déri, Fruzsina Petrovay, Lili Jármi, Bernadett Pályi, Eszter Balla, Zoltán Kis |
| EPI_ISL_13408799, EPI_ISL_13408801 | Public Health Agency of Canada, National Microbiology Laboratory | Public Health Agency of Canada, National Microbiology Laboratory | Knox,N., Hole,D., Duggan,A., Yadav,C., Haidl,E., Chapel,M., Graham,M., Domselaar,G.V., Jolly,G., Audet,J., Fernando,L., Antonation,K., Hagan,M., Griffiths,E., Leung,A., Safronetz,D., Eshaghi,A., Gubbay,J.B., Hasso,M., Marchand-Austin,A., Olsha,R. and Patel,S.N. |
| EPI_ISL_13408807, EPI_ISL_13408809, EPI_ISL_13408811, EPI_ISL_13408813, EPI_ISL_13408815, EPI_ISL_13408817, EPI_ISL_13408819, EPI_ISL_13408821, EPI_ISL_13408823, EPI_ISL_13408827, EPI_ISL_13408831, EPI_ISL_13408833, EPI_ISL_13408835 | | | |
| see above | Public Health Agency of Canada, National Microbiology Laboratory | Public Health Agency of Canada, National Microbiology Laboratory | ncknox |
| EPI_ISL_13408837, EPI_ISL_13408841, EPI_ISL_13408843, EPI_ISL_13408847, EPI_ISL_13408849, EPI_ISL_13408851, EPI_ISL_13408853, EPI_ISL_13408855, EPI_ISL_13408857, EPI_ISL_13408859, EPI_ISL_13408861 | | | |
| see above | Public Health Agency of Canada, National Microbiology Laboratory | Public Health Agency of Canada, National Microbiology Laboratory | Knox,N., Duggan,A., Yadav,C., Hole,D., Haidl,E., Chapel,M., Jolly,G., Domselaar,G.V., Antonation,K., Leung,A., Fernando,L., Audet,J., Hagan,M., Graham,M., Griffiths,E., Safronetz,D., Charest,H., Levade,I. and Fafard,J. |
| EPI_ISL_13411153, EPI_ISL_13411154, EPI_ISL_13411155, EPI_ISL_13411156, EPI_ISL_13411157, EPI_ISL_13411158 | Centre for Biological Threats, Highly Pathogenic Viruses, Robert Koch Institute | Centre for Biological Threats, Highly Pathogenic Viruses, Robert Koch Institute | Brinkmann,A., Kohl,C., Uddin,S., Pape,K., Schrick,L., Michel,J., Schaade,L. and Nitsche,A. |
| EPI_ISL_13411159, EPI_ISL_13411160, EPI_ISL_13411161, EPI_ISL_13411162 | Centre for Biological Threats, Highly Pathogenic Viruses, Robert Koch Institute | Centre for Biological Threats, Highly Pathogenic Viruses, Robert Koch Institute | Brinkmann,A., Kohl,C., Uddin,S., Pape,K., Schrick,L., Michel,J., Stocker,H., Schaade,L. and Nitsche,A. |
| EPI_ISL_13411163, EPI_ISL_13411164, EPI_ISL_13411165 | Centre for Biological Threats, Highly Pathogenic Viruses, Robert Koch Institute | Centre for Biological Threats, Highly Pathogenic Viruses, Robert Koch Institute | Brinkmann,A., Kohl,C., Uddin,S., Pape,K., Schrick,L., Michel,J., Schaade,L. and Nitsche,A. |
| EPI_ISL_13411166, EPI_ISL_13411167 | Centre for Biological Threats, Highly Pathogenic Viruses, Robert Koch Institute | Centre for Biological Threats, Highly Pathogenic Viruses, Robert Koch Institute | Brinkmann,A., Kohl,C., Uddin,S., Pape,K., Schrick,L., Michel,J., Jessen,H., Schaade,L. and Nitsche,A. |
| EPI_ISL_13436658 | Coordenadoria de Vigilancia em Saude - Sao Paulo | Instituto Adolfo Lutz Strategic Laboratory | Claudio Tavares Sacchi, Karoline Rodrigues Campos, Ariadne Fereira Amarante, Adriano Abbud, Adriana Bugno |
| EPI_ISL_13436792 | Hospital Santa Ignes | Instituto Adolfo Lutz Strategic Laboratory | Claudio Tavares Sacchi, Karoline Rodrigues Campos, Adriano Abbud, Adriana Bugno |
| EPI_ISL_13437056 | Hosp. Alemao Oswaldo Cruz | Instituto Adolfo Lutz Strategic Laboratory | Claudio Tavares Sacchi, Karoline Rodrigues Campos, Ariadne Ferreira Amarante, Adriano Abbud, Adriana Bugno |
| EPI_ISL_13445553 | Laboratory for Diagnostics of Zoonoses and WHO Centre, Institute of Microbiology and Immunology, Faculty of Medicine, University of Ljubljana | Laboratory for Diagnostics of Zoonoses and WHO Centre, Institute of Microbiology and Immunology, Faculty of Medicine, University of Ljubljana | Zakotnik,S., Vlaj,D., Suljic,A., Zorec,T.M., Korva,M., Poljak,M. and Avsic Zupanc,T. |
| EPI_ISL_13449965, EPI_ISL_13449966 | Hospital Universitario La Paz, Microbiology | Hospital Universitario La Paz, Microbiology | de la Hoz-Sanchez,B., Lopez-Ortiz,M., Gutierrez-Arroyo,A., Roces-Alvarez,P., Lazaro-Peona,F., Dahdouh,E., Bloise,I., Garcia-Rodriguez,J. and Mingorance,J. |
| EPI_ISL_13459346 | CRT-DST-AIDS | Instituto Adolfo Lutz Strategic Laboratory | Claudio Tavares Sacchi, Karoline Rodrigues Campos, Ariadne Ferreira Amarante, Adriano Abbud, Adriana Bugno |
| EPI_ISL_13459347, EPI_ISL_13459482, EPI_ISL_13459483 | Instituto de Infectologia Emilio Ribas | Instituto Adolfo Lutz Strategic Laboratory | Claudio Tavares Sacchi, Karoline Rodrigues Campos, Ariadne Ferreira Amarante, Adriano Abbud, Adriana Bugno |
| EPI_ISL_13466448, EPI_ISL_13466449, EPI_ISL_13466450, EPI_ISL_13466451, EPI_ISL_13466452, EPI_ISL_13466453, EPI_ISL_13466455, EPI_ISL_13466456, EPI_ISL_13466457, EPI_ISL_13466459, EPI_ISL_13466460, EPI_ISL_13466461, EPI_ISL_13466465 | | | |

| | | | |
|---|---|---|---|
| see above | Department of Infectious Diseases, National Institute of Health Doutor Ricardo Jorge, Portugal (INSA) | Department of Infectious Diseases, National Institute of Health Doutor Ricardo Jorge, Portugal (INSA) | Isidro,J., Borges,V., Pinto,M., Sobral,D., Santos,J., Nunes,A., Mixao,V., Ferreira,R., Santos,D., Duarte,S., Vieira,L., Borrego,M.J., Nuncio,S., Lopes de Carvalho,I., Pelerito,A., Cordeiro,R., Gomes,J.P. |
| EPI_ISL_13483155, EPI_ISL_13483157, EPI_ISL_13483159, EPI_ISL_13483161, EPI_ISL_13483162, EPI_ISL_13483163, EPI_ISL_13483164 | Centre for Biological Threats, Highly Pathogenic Viruses, Robert Koch Institute | Centre for Biological Threats, Highly Pathogenic Viruses, Robert Koch Institute | Brinkmann,A., Kohl,C., Uddin,S., Pape,K., Schrick,L., Michel,J., Jessen,H., Schaade,L. and Nitsche,A. |
| EPI_ISL_13483165, EPI_ISL_13483167, EPI_ISL_13483168, EPI_ISL_13483170, EPI_ISL_13483171, EPI_ISL_13483173, EPI_ISL_13483177, EPI_ISL_13483178, EPI_ISL_13483180, EPI_ISL_13483182, EPI_ISL_13483183, EPI_ISL_13483185, EPI_ISL_13483187, EPI_ISL_13483188, EPI_ISL_13483191, EPI_ISL_13483193, EPI_ISL_13483195, EPI_ISL_13483196, EPI_ISL_13483198, EPI_ISL_13483200, EPI_ISL_13483201, EPI_ISL_13483203, EPI_ISL_13483205, EPI_ISL_13483206, EPI_ISL_13483208 | | | |
| see above | Centre for Biological Threats, Highly Pathogenic Viruses, Robert Koch Institute | Centre for Biological Threats, Highly Pathogenic Viruses, Robert Koch Institute | Brinkmann,A., Kohl,C., Uddin,S., Pape,K., Schrick,L., Michel,J., Schaade,L. and Nitsche,A. |
| EPI_ISL_13484458 | Laboratorio de Enterovirus, Instituto Oswaldo Cruz, Fiocruz | Instituto Oswaldo Cruz FIOCRUZ - Laboratory of Respiratory Viruses and Measles (LVRS) | Paola Resende, Elisa Cavalcante Pereira, Bruna Mendonça da Silva, Jéssica Graça Macedo de Carvalho, Larissa Macedo Pinto, Victor Guimaraes, Marilda Siqueira, Renan da Silva Faustino, Marilia Santini, Edson Elias da Silva on behalf of the Fiocruz Genomic Surveillance Network |
| EPI_ISL_13498265 | National Institute for Communicable Diseases of the National Health Laboratory Service | National Institute for Communicable Diseases of the National Health Laboratory Service | Chan WY, Mtshali PS, Grobbelaar A, Moolla N, Mohale T, Du Plessis MG, Ismail A, Weyer J |
| EPI_ISL_13502582 | Laboratory of Microbiology and Virology, Ospedale Amedeo di Savoia, ASL "Città di Torino" | Laboratory of Microbiology and Virology, Ospedale Amedeo di Savoia, ASL "Città di Torino" | Francesco Cerutti, Antonella Bottoni, Marisa Cazzadore, Tiziano Allice, Maria Grazia Milia, Gabriella Gregori, Elisa Burdino, Valeria Ghisetti |
| EPI_ISL_13508393 | Hosp. Itacolomy Butanta | Instituto Adolfo Lutz Strategic Laboratory | Claudio Tavares Sacchi, Karoline Rodrigues Campos, Ariadne Ferreira Amarante, Adriano Abbud, Adriana Bugno |
| EPI_ISL_13508471 | Instituto de Infectologia Emilio Ribas | Instituto Adolfo Lutz Strategic Laboratory | Claudio Tavares Sacchi, Karoline Rodrigues Campos, Ariadne Ferreira Amarante, Adriano Abbud, Adriana Bugno |
| EPI_ISL_13530881 | Laboratorio de Referencia Nacional de Virus Respiratorios. Centro Nacional de Salud Publica. Instituto Nacional de Salud Peru. | Laboratorio de Referencia Nacional de Virus Respiratorios. Centro Nacional de Salud Publica. Instituto Nacional de Salud Peru. | Carlos Padilla Rojas, Veronica Hurtado Vela, Iris Silva Molina, Luren Sevilla Castañeda, Victor Jimenez Vasquez, Orson Mestanza Millones, Luis Barcena Flores, Wendy Lizarraga Olivares, Alicia Nuñez Llanos, Steve Acedo Lazo, Francisco Ascue Orosco, Kelly Izarra Rojas, Princesa Medrano Alhuay, Karla Vasquez Cajachahua, Estela Huaman Angeles, Jorge Giraldo Chavez, Lilian Huarca Balbin, Lisbet Roxana Inga Angulo, Maria Sandra Villar Saavedra, Henri Bailon Calderon, Lely Solari Zerpa, Gloria Arotinco Garayar. Equipo de vigilancia genomica del Instituto Nacional de Salud. |
| EPI_ISL_13537923 | Microbiology, Immunology and Transplantation, KU Leuven, Rega Institute | Microbiology, Immunology and Transplantation, KU Leuven, Rega Institute | Wawina-Bokalanga,T., Vanmechelen,B., Logist,A.-S., Sinnesael,R., Ysebaert,L., Bloemen,M. and Maes,P. |
| EPI_ISL_13537924, EPI_ISL_13537925, EPI_ISL_13537926 | Microbiology, Immunology and Transplantation, KU Leuven, Rega Institute | Microbiology, Immunology and Transplantation, KU Leuven, Rega Institute | Vanmechelen,B., Wawina-Bokalanga,T., Logist,A.-S., Sinnesael,R., Ysebaert,L., Verlinden,J., Van Holm,B., Bloemen,M. and Maes,P. |
| EPI_ISL_13544223, EPI_ISL_13544226, EPI_ISL_13544227, EPI_ISL_13544228, EPI_ISL_13544229, EPI_ISL_13544230, EPI_ISL_13544231, EPI_ISL_13544233, EPI_ISL_13544234, EPI_ISL_13544235 | Public Health Agency of Canada, National Microbiology Laboratory | Public Health Agency of Canada, National Microbiology Laboratory | Duggan,A., Hole,D., Knox,N., Yadav,C., Haidl,E., Chapel,M., Domselaar,G.V., Jolly,G., Audet,J., Fernando,L., Antonation,K., Safronetz,D., Hagan,M., Griffiths,E., Leung,A., Graham,M., Peters,G., Go,A., Laminman,V., Kaplen,B., Eshaghi,A., Gubbay,J.B., Hasso,M., Marchand-Austin,A., Olsha,R. and Patel,S.N. |
| EPI_ISL_13544237, EPI_ISL_13544239, EPI_ISL_13544241, EPI_ISL_13544244, EPI_ISL_13544245, EPI_ISL_13544247, EPI_ISL_13544248, EPI_ISL_13544249, EPI_ISL_13544252, EPI_ISL_13544254, EPI_ISL_13544255, EPI_ISL_13544256, EPI_ISL_13544257, EPI_ISL_13544258, EPI_ISL_13544260, EPI_ISL_13544261, EPI_ISL_13544263, EPI_ISL_13544264, EPI_ISL_13544265, EPI_ISL_13544266, EPI_ISL_13544267 | | | |
| see above | Public Health Agency of Canada, National Microbiology Laboratory | Public Health Agency of Canada, National Microbiology Laboratory | Duggan,A., Hole,D., Knox,N., Yadav,C., Haidl,E., Chapel,M., Domselaar,G.V., Fernando,L., Graham,M., Antonation,K., Audet,J., Hagan,M., Safronetz,D., Leung,A., Peters,G., Go,A., Laminman,V., Kaplen,B., Jolly,G., Charest,H., Levade,I. and Fafard,J. |
| EPI_ISL_13573943 | Center for Virology, Medical University of Vienna | Medical University of Vienna Center for Virology | Jeremy V. Camp, Monika Redlberger-Fritz, Stephan W. Aberle |
| EPI_ISL_13584854, EPI_ISL_13586184 | Institute for Virology, Philipps-University Marburg | Institute for Virology, Philipps-University Marburg | Eickmann, M., Lier, C., Kowalski, K., Kraft, F., Becker, S. |
| EPI_ISL_13607904 | Servicio de Infectologia, Hospital Universitario Dr. José Eleuterio Gonzalez, Universidad Autonoma de Nuevo Leon | Centro de Investigacion e Innovacion en Virologia Medica, Departamento de Bioquimica y Medicina Molecular, Facultad de Medicina, Universidad Autonoma de Nuevo Leon | Kame A. Galan-Huerta, Manuel Paz Infanzon, Ali F. Ruiz Higareda, Laura Nuzzolo-Shihadeh, Adrian Camacho-Ortiz, Paola Bocanegra-Ibarias, Ana M. Rivas-Estilla, Daniel Zacarias-Villarreal, Luis A. Yamallel-Ortega, Maria D. Guerrero-Putz, Jorge Ocampo-Candiani |
| EPI_ISL_13624509 | Instituto de Diagnóstico y Referencia Epidemiológicos/Jurisdicción Sanitaria Cuauhtémoc/Hospital Ángeles Roma | Instituto de Diagnóstico y Referencia Epidemiológicos/Instituto de Biotecnología UNAM | Adnan Araiza-Rodríguez, Adriana Salvador-Patiño, Alejandro Sánchez-Flores, América del Pilar Mandujano-Martínez, Blanca Taboada, Carlos Eduardo Hernández-Sánchez, Carlos F. Arias, Claudia Elena Wong-Arámbula, Daniel José Regalado-Santiago, David Esaú Fragoso-Fonseca, Elizabeth Andrade-Montiel, Fabiola Garcés-Ayala, Fernando González-Dominguez, Gabriel García-Rodríguez, Gloria Vázquez-Castro, Hugo López Gatell Ramírez, Irma López-Martínez, Jerome Verleyen, Jesús Trujillo, Jorge Ochoa, José Ernesto Ramírez-González, Karel Estrada-Guerra, Lucía Hernández-Rivas, Magaly Guadalupe Landa-Flores, Maribel González-Villa, Mireya Mederos-Michel, Nancy Martínez-Velázquez, Noé Escobar-Escamilla, Oliva López, Ricardo Cortés-Alcalá, Ricardo Grande, Verónica Jiménez-Jacinto |
| EPI_ISL_13632071 | Center of Diagnostics and Vaccine Development, Centers for Disease Control, Taiwan | Center of Diagnostics and Vaccine Development, Centers for Disease Control, Taiwan | Jih-Hui Lin, Shu-Chun Chiu, Hsin-I, Huang, Wei-Lun Huang, Wen-Bin, Fann, Pei-Yu, Hsieh, Jyh-Yuan Yang |
| EPI_ISL_13632288 | National Institute for Communicable Diseases of the National Health Laboratory Service | National Institute for Communicable Diseases of the National Health Laboratory Service | Chan WY, Mtshali PS, Grobbelaar A, Moolla N, Mohale T, Lowe M, Du Plessis MG, Ismail A, Weyer J |
| EPI_ISL_13651348, EPI_ISL_13651349, EPI_ISL_13651350 | Laboratorio de Referencia Nacional de Virus Respiratorio. Centro Nacional de Salud Publica. Instituto Nacional de Salud. | Laboratorio de Referencia Nacional de Virus Respiratorio. Centro Nacional de Salud Publica. Instituto Nacional de Salud. | Carlos Padilla Rojas, Veronica Hurtado Vela, Iris Silva Molina, Luren Sevilla Castañeda, Victor Jimenez Vasquez, Orson Mestanza Millones, Luis Barcena Flores, Wendy Lizarraga Olivares, Alicia Nuñez Llanos, Steve Acedo Lazo, Francisco Ascue Orosco, Kelly Izarra Rojas, Princesa Medrano Alhuay, Karla Vasquez Cajachahua, Estela Huaman Angeles, Jorge Giraldo Chavez, Lilian Huarca Balbin, Lisbet Roxana Inga Angulo, Maria Sandra Villar Saavedra, Henri Bailon Calderon, Lely Solari Zerpa, Gloria Arotinco Garayar. Equipo de vigilancia genomica del Instituto Nacional de Salud. |
| EPI_ISL_13658019, EPI_ISL_13658021 | Erasmus Medical Center Department of Virology | Erasmus Medical Center Department of Virology | Bas Oude Munnink, Marjan Boter, Babette Weller, Richard Molenkamp, Janette Rahamat-Langendoen, Reina Sikkema, Marion Koopmans |
| EPI_ISL_13705358 | Hosp. Alemao Oswaldo Cruz | Instituto Adolfo Lutz Strategic Laboratory | Claudio Tavares Sacchi, Karoline Rodrigues Campos, Ariadne Ferreira Amarante, Marlon Benedito Nascimento Santos, Alex Domingos Reis, Adriano Abbud, Adriana Bugno |
| EPI_ISL_13705407 | Hosp. Sirio-Libanes | Instituto Adolfo Lutz Strategic Laboratory | Claudio Tavares Sacchi, Karoline Rodrigues Campos, Ariadne Ferreira Amarante, Marlon Benedito Nascimento Santos, Alex Domingos Reis, Adriano Abbud, Adriana Bugno |
| EPI_ISL_13728303 | Department of Medical Microbiology & Infection prevention, Amsterdam University Medical Centers location AMC | Department of Medical Microbiology & Infection prevention, Amsterdam University Medical Centers location AMC | Matthijs Welkers, Jelle Koopsen, Robin van Houdt, Marcel Jonges, Sebastien Matamoros, Sjoerd Rebers, Fokla Zorgdrager, Sylvia Bruisten, Judith den Uil, Akke Cornelissen, Janke Schinkel, Menno de Jong, Gini van Rijckevorsel and Mariken van der Lubben on behalf of the Amsterdam Regional Genomic epidemiology and Outbreak Surveillance (ARGOS) consortium |
| EPI_ISL_13732932 | Hosp. Sao Joaquim - Beneficiencia Portuguesa | Instituto Adolfo Lutz Strategic Laboratory | Claudio Tavares Sacchi, Karoline Rodrigues Campos, Ariadne Ferreira Amarante, Marlon Benedito Nascimento Santos, Alex Domingos Reis, Adriano Abbud, Adriana Bugno |
| EPI_ISL_13734237, EPI_ISL_13734238, EPI_ISL_13734239, EPI_ISL_13734240, EPI_ISL_13734241, EPI_ISL_13734242, EPI_ISL_13734243, EPI_ISL_13734244, EPI_ISL_13734245, EPI_ISL_13734246, EPI_ISL_13734247, EPI_ISL_13734248, EPI_ISL_13734249, EPI_ISL_13734251, EPI_ISL_13734252, EPI_ISL_13734253, EPI_ISL_13734254, EPI_ISL_13734255, EPI_ISL_13734256, EPI_ISL_13734257, EPI_ISL_13734258, EPI_ISL_13734259, EPI_ISL_13734260, EPI_ISL_13734261, EPI_ISL_13734262, EPI_ISL_13734263, EPI_ISL_13734264, EPI_ISL_13734265, EPI_ISL_13734266, EPI_ISL_13734267, EPI_ISL_13734268 | | | |
| see above | Centre for Biological Threats, Highly Pathogenic Viruses, Robert Koch Institute | Centre for Biological Threats, Highly Pathogenic Viruses, Robert Koch Institute | Brinkmann,A., Kohl,C., Pape,K., Uddin,S., Schrick,L., Michel,J., Schaade,L. and Nitsche,A. |
| EPI_ISL_13734270 | Centers for Disease Control & Prevention (CDC), Division of High Consequence Pathogens and Pathology (DHCPP-PRB) | Centers for Disease Control & Prevention (CDC), Division of High Consequence Pathogens and Pathology (DHCPP-PRB) | Gigante,C.M., Ventura,J., Seabolt,M.H., Zhao,H., Wilkins,K., Respress,J., Howard,D., Batra,D., McCollum,A., Hutson,C., Davidson,W., Rao,A., Nash,J. and Li,Y. |
| EPI_ISL_13744896 | Centers for Disease Control & Prevention (CDC), Division of High Consequence Pathogens and Pathology (DHCPP-PRB) | Centers for Disease Control & Prevention (CDC), Division of High Consequence Pathogens and Pathology (DHCPP-PRB) | Gigante,C.M., Ghinai,I., Seabolt,M.H., Zhao,H., Wilkins,K., Respress,J., Howard,D., Batra,D., McCollum,A., Hutson,C., Davidson,W., Rao,A., Kerins,J. and Li,Y. |

| | | | |
|---|---|---|---|
| EPI_ISL_13744897, EPI_ISL_13744898 | Centers for Disease Control & Prevention (CDC), Division of High Consequence Pathogens and Pathology (DHCPP-PRB) | Centers for Disease Control & Prevention (CDC), Division of High Consequence Pathogens and Pathology (DHCPP-PRB) | Gigante,C.M., Hughes,S., Seabolt,M.H., Zhao,H., Wilkins,K., Respress,J., Howard,D., Batra,D., McCollum,A., Hutson,C., Davidson,W., Rao,A., Baumgartner,J. and Li,Y. |
| EPI_ISL_13744899 | Centers for Disease Control & Prevention (CDC), Division of High Consequence Pathogens and Pathology (DHCPP-PRB) | Centers for Disease Control & Prevention (CDC), Division of High Consequence Pathogens and Pathology (DHCPP-PRB) | Gigante,C.M., Ghinai,I., Seabolt,M.H., Zhao,H., Wilkins,K., Respress,J., Howard,D., Batra,D., McCollum,A., Hutson,C., Davidson,W., Rao,A., Kerins,J. and Li,Y. |
| EPI_ISL_13744900, EPI_ISL_13744901 | Centers for Disease Control & Prevention (CDC), Division of High Consequence Pathogens and Pathology (DHCPP-PRB) | Centers for Disease Control & Prevention (CDC), Division of High Consequence Pathogens and Pathology (DHCPP-PRB) | Gigante,C.M., Hughes,S., Seabolt,M.H., Zhao,H., Wilkins,K., Respress,J., Howard,D., Batra,D., McCollum,A., Hutson,C., Davidson,W., Rao,A., Baumgartner,J. and Li,Y. |
| EPI_ISL_13744902 | Department of Virology, Faculty of Medicine, University of Helsinki | Department of Virology, Faculty of Medicine, University of Helsinki | Kant,R., Smura,T., Vauhkonen,H. and Vapalahti,O. |
| EPI_ISL_13744903, EPI_ISL_13744904, EPI_ISL_13744905 | Centre for Biological Threats, Highly Pathogenic Viruses, Robert Koch Institute | Centre for Biological Threats, Highly Pathogenic Viruses, Robert Koch Institute | Brinkmann,A., Kohl,C., Pape,K., Uddin,S., Schrick,L., Michel,J., Jessen,H., Schaade,L. and Nitsche,A. |
| EPI_ISL_13744906, EPI_ISL_13744907, EPI_ISL_13744908, EPI_ISL_13744909, EPI_ISL_13744910, EPI_ISL_13744911, EPI_ISL_13744912, EPI_ISL_13744913, EPI_ISL_13744914, EPI_ISL_13744915, EPI_ISL_13744916, EPI_ISL_13744917, EPI_ISL_13744918, EPI_ISL_13744919, EPI_ISL_13744920, EPI_ISL_13744921, EPI_ISL_13744922, EPI_ISL_13744923, EPI_ISL_13744924, EPI_ISL_13744925, EPI_ISL_13744926, EPI_ISL_13744927, EPI_ISL_13744928, EPI_ISL_13744929, EPI_ISL_13744930, EPI_ISL_13744931 | | | |
| see above | Centre for Biological Threats, Highly Pathogenic Viruses, Robert Koch Institute | Centre for Biological Threats, Highly Pathogenic Viruses, Robert Koch Institute | Brinkmann,A., Kohl,C., Pape,K., Uddin,S., Schrick,L., Michel,J., Schaade,L. and Nitsche,A. |
| EPI_ISL_13822667, EPI_ISL_13822668, EPI_ISL_13822669, EPI_ISL_13822718 | Erasmus Medical Center Department of Virology | Erasmus Medical Center Department of Virology | Bas Oude Munnink, Marjan Boter, Babette Weller, Richard Molenkamp, Janette Rahamat-Langendoen, Reina Sikkema, Marion Koopmans |
| EPI_ISL_13827274, EPI_ISL_13827275, EPI_ISL_13827277, EPI_ISL_13827278, EPI_ISL_13827279, EPI_ISL_13827280, EPI_ISL_13827282 | Public Health Agency of Canada, National Microbiology Laboratory | Public Health Agency of Canada, National Microbiology Laboratory | Duggan,A., Hole,D., Yadav,C., Knox,N., Haidl,E., Chapel,M., Domselaar,G.V., Fernando,L., Graham,M., Antonation,K., Audet,J., Hagan,M., Safronetz,D., Leung,A., Peters,G., Go,A., Laminman,V., Kaplen,B., Jolly,G., Marchand-Austin,A., Eshaghi,A., Patel,S.N., Hasso,M., Gubbay,J.B. and Olsha,R. |
| EPI_ISL_13833194, EPI_ISL_13833195, EPI_ISL_13833196, EPI_ISL_13833197 | Laboratorio de Referencia Nacional de Virus Respiratorio. Centro Nacional de Salud Publica. Instituto Nacional de Salud. | Laboratorio de Referencia Nacional de Virus Respiratorio. Centro Nacional de Salud Publica. Instituto Nacional de Salud. | Carlos Padilla Rojas, Veronica Hurtado Vela, Iris Silva Molina, Luren Sevilla Castañeda, Victor Jimenez Vasquez, Orson Mestanza Millones, Luis Barcena Flores, Wendy Lizarraga Olivares, Alicia Nuñez Llanos, Steve Acedo Lazo, Francisco Ascue Orosco, Kelly Izarra Rojas, Princesa Medrano Alhuay, Karla Vasquez Cajachahua, Estela Huaman Angeles, Jorge Giraldo Chavez, Lilian Huarca Balbin, Lisbet Roxana Inga Angulo, Maria Sandra Villar Saavedra, Henri Bailon Calderon, Lely Solari Zerpa, Gloria Arotinco Garayar. Equipo de vigilancia genomica del Instituto Nacional de Salud. |
| EPI_ISL_13842269, EPI_ISL_13842548 | Center for Virology, Medical University of Vienna | Medical University of Vienna Center for Virology | Jeremy V. Camp, Monika Redlberger-Fritz, Stephan W. Aberle |
| EPI_ISL_13889435, EPI_ISL_13889436, EPI_ISL_13889438, EPI_ISL_13889439, EPI_ISL_13889440, EPI_ISL_13889441, EPI_ISL_13889442, EPI_ISL_13889443, EPI_ISL_13889444, EPI_ISL_13889445, EPI_ISL_13889446, EPI_ISL_13889447, EPI_ISL_13889448, EPI_ISL_13889449, EPI_ISL_13889450, EPI_ISL_13889515, EPI_ISL_13889590, EPI_ISL_13889660, EPI_ISL_13889729, EPI_ISL_13889796, EPI_ISL_13889908, EPI_ISL_13889977, EPI_ISL_13890048, EPI_ISL_13890135, EPI_ISL_13890204, EPI_ISL_13890273, EPI_ISL_13890464, EPI_ISL_13890465, EPI_ISL_13890466, EPI_ISL_13890467, EPI_ISL_13890468, EPI_ISL_13890469, EPI_ISL_13890471, EPI_ISL_13890472, EPI_ISL_13890473, EPI_ISL_13890474, EPI_ISL_13890475, EPI_ISL_13890476, EPI_ISL_13890478, EPI_ISL_13890479, EPI_ISL_13890481 | | | |
| see above | Charité Universitätsmedizin Berlin, Institut für Virologie/Labor Berlin | Charité Universitätsmedizin Berlin, Institut für Virologie | Terry C. Jones, Julia Schneider, Barbara Mühlemann, Talitha Veith, Jörn Beheim-Schwarzbach, Julia Tesch, Marie Luisa Schmidt, Felix Walper, Tobias Bleicker, Caroline Isner, Frieder Pfäfflin, Ricardo Niklas Werner, Victor M. Corman, Christian Drosten |
| EPI_ISL_13908332, EPI_ISL_13908333, EPI_ISL_13908334, EPI_ISL_13908335, EPI_ISL_13908336, EPI_ISL_13908337, EPI_ISL_13908338, EPI_ISL_13908339, EPI_ISL_13908340, EPI_ISL_13908341, EPI_ISL_13908342, EPI_ISL_13908343 | | | |
| see above | Public Health Agency of Canada, National Microbiology Laboratory | Public Health Agency of Canada, National Microbiology Laboratory | Duggan,A., Hole,D., Yadav,C., Knox,N., Chapel,M., Tyler,A., Haidl,E., Domselaar,G.V., Antonation,K., Audet,J., Fernando,L., Hagan,M., Safronetz,D., Graham,M., Peters,G., Go,A., Laminman,V., Kaplen,B., Leung,A., Jolly,G., Fafard,J., Charest,H. and Levade,I. |
| EPI_ISL_13908346, EPI_ISL_13908347, EPI_ISL_13908348, EPI_ISL_13908349 | Centre for Biological Threats, Highly Pathogenic Viruses, Robert Koch Institute | Centre for Biological Threats, Highly Pathogenic Viruses, Robert Koch Institute | Brinkmann,A., Kohl,C., Pape,K., Uddin,S., Schrick,L., Michel,J., Jessen,H., Schaade,L. and Nitsche,A. |
| EPI_ISL_13908350, EPI_ISL_13908351, EPI_ISL_13908352, EPI_ISL_13908353, EPI_ISL_13908354, EPI_ISL_13908355, EPI_ISL_13908356, EPI_ISL_13908357, EPI_ISL_13908358, EPI_ISL_13908359, EPI_ISL_13908360, EPI_ISL_13908361, EPI_ISL_13908362, EPI_ISL_13908363, EPI_ISL_13908364, EPI_ISL_13908365 | | | |
| see above | Centre for Biological Threats, Highly Pathogenic Viruses, Robert Koch Institute | Centre for Biological Threats, Highly Pathogenic Viruses, Robert Koch Institute | Brinkmann,A., Kohl,C., Pape,K., Uddin,S., Schrick,L., Michel,J., Schaade,L. and Nitsche,A. |
| EPI_ISL_13958697 | Research and Evaluation, UKHSA | Research and Evaluation, UKHSA | Groves,N., Osman,K.L., Lewandowski,K.S., Carter,D.P., Pullan,S.T., Myers,R., Vipond,R. and Chand,M. |
| EPI_ISL_13983356 | INSPI-Centro de Referencia Nacional de Virus Exantemáticos, Gastroentéricos y Transmitido por Vectores. | INSPI-Dirección Técnica de Investigación, Desarrollo e Innovación INSPI-Centro de Referencia Nacional de Genómica, Secuenciación y Bioinformática | Andrés Carrazco-Montalvo, Diana Gutiérrez, Naomi Mora, Silvia Salgado-Cisneros, Johana Parrales-Valdiviezo, Martha Sánchez-Domenech, Diego Morales, Gulnara Borja-Cabrera, Leandro Patiño*. |
| EPI_ISL_13993735, EPI_ISL_13993737, EPI_ISL_13993738, EPI_ISL_13993739 | California Department of Public Health | California Department of Public Health | Viral and Rickettsial Disease Laboratory |
| EPI_ISL_14003930 | University of Rochester Medical Center | University of Rochester Medical Center | Andrew Cameron, Mondraya Howard, Sara Connelly, Dwight Hardy, Kelly DeLary |
| EPI_ISL_14021725 | Hosp. Municipal Enf. Antonio Policarpo de Oliveira | Instituto Adolfo Lutz Strategic Laboratory | Claudio Tavares Sacchi, Karoline Rodrigues Campos, Ariadne Ferreira Amarante, Marlon Benedito Nascimento Santos, Alex Domingos Reis, Adriano Abbud, Adriana Bugno |
| EPI_ISL_14033204, EPI_ISL_14033205, EPI_ISL_14033206, EPI_ISL_14033207, EPI_ISL_14033208, EPI_ISL_14033209, EPI_ISL_14033210, EPI_ISL_14033211, EPI_ISL_14033212, EPI_ISL_14033213 | Laboratory Medicine, UW Virology | Laboratory Medicine, UW Virology | Sereewit,J., Xie,H., Pavitra,R. and Greninger,A. |
| EPI_ISL_14050451, EPI_ISL_14050453, EPI_ISL_14050454, EPI_ISL_14050458 | Public Health Agency of Canada, National Microbiology Laboratory | Public Health Agency of Canada, National Microbiology Laboratory | Duggan,A., Hole,D., Yadav,C., Knox,N., Tyler,A., Haidl,E., Chapel,M., Domselaar,G.V., Graham,M., Audet,J., Fernando,L., Hagan,M., Safronetz,D., Leung,A., Peters,G., Go,A., Laminman,V., Kaplen,B., Antonation,K., Jolly,G., Griffiths,E., Charest,H., Levade,I. and Fafard,J. |
| EPI_ISL_14070493, EPI_ISL_14070852, EPI_ISL_14070855 | Instituto de Infectologia Emilio Ribas | Instituto Adolfo Lutz Strategic Laboratory | Claudio Tavares Sacchi, Karoline Rodrigues Campos, Ariadne Ferreira Amarante, Marlon Benedito Nascimento Santos, Alex Domingos Reis, Adriano Abbud, Adriana Bugno |
| EPI_ISL_14153982 | Vajira Hospital | National Institute of Health, Department of Medical Sciences, Ministry of Public Health, Thailand | Pilailuk Okada; Siripaporn Phuygun; Nuttida Thongpramul; Thanutsapa Thanadachakul; Kazuhisa Okada; Archawin Rojanawiwat; Chakkarat Pitayawonganon; Supakit Sirilak |
| EPI_ISL_14166709 | Medical University of Vienna Center for Virology | Medical University of Vienna Center for Virology | Jeremy V Camp, Monika Redlberger-Fritz, Stephan W. Aberle |
| EPI_ISL_14167248, EPI_ISL_14167573, EPI_ISL_14167574, EPI_ISL_14167575 | Medical University of Vienna Center for Virology | Medical University of Vienna Center for Virology | Jeremy V. Camp, Monika Redlberger-Fritz, Stephan W. Aberle |
| EPI_ISL_14181948, EPI_ISL_14181949, EPI_ISL_14181951, EPI_ISL_14181952, EPI_ISL_14181953, EPI_ISL_14181954, EPI_ISL_14181955, EPI_ISL_14181956, EPI_ISL_14181957, EPI_ISL_14181958, EPI_ISL_14181959, EPI_ISL_14181960 | | | |
| see above | Institute of Health Carlos III, Bioinformatics Unit | Institute of Health Carlos III, Bioinformatics Unit | Cuesta,I. |
| EPI_ISL_14207724, EPI_ISL_14207725, EPI_ISL_14207726, EPI_ISL_14207730, EPI_ISL_14207731, EPI_ISL_14207733, EPI_ISL_14207734, EPI_ISL_14207735, EPI_ISL_14207738, EPI_ISL_14207739, EPI_ISL_14207740, EPI_ISL_14207741 | | | |
| see above | Laboratorio de Referencia Nacional de Virus Respiratorio. Centro Nacional de Salud Publica. Instituto Nacional de Salud. | Laboratorio de Referencia Nacional de Virus Respiratorio. Centro Nacional de Salud Publica. Instituto Nacional de Salud. | Carlos Padilla Rojas, Veronica Hurtado Vela, Iris Silva Molina, Luren Sevilla Castañeda, Victor Jimenez Vasquez, Orson Mestanza Millones, Luis Barcena Flores, Wendy Lizarraga Olivares, Alicia Nuñez Llanos, Steve Acedo Lazo, Francisco Ascue Orosco, Kelly Izarra Rojas, Princesa Medrano Alhuay, Karla Vasquez Cajachahua, Estela Huaman Angeles, Jorge Giraldo Chavez, Lilian Huarca Balbin, Lisbet Roxana Inga Angulo, Maria Sandra Villar Saavedra, Henri Bailon Calderon, Lely Solari Zerpa, Gloria Arotinco Garayar. Equipo de vigilancia genomica del Instituto Nacional de Salud. |
| EPI_ISL_14211644, EPI_ISL_14211645 | Public Health Authority of the Slovak Republic | Laboratory of Genomics and Bioinformatics, Comenius University Science Park | Tomáš Szemes, Edita Staroová, Elena Tichá, Lucia Ševíková, Terézia Vrabová, Tatiana Sedláková, Miroslav Böhmer, Jaroslav Budiš, Pavol Mišenko |
| EPI_ISL_14216746, EPI_ISL_14216750, EPI_ISL_14216752, EPI_ISL_14216753, EPI_ISL_14216754, EPI_ISL_14216755, EPI_ISL_14216756, EPI_ISL_14216757, EPI_ISL_14216758, EPI_ISL_14216759, EPI_ISL_14216761, EPI_ISL_14216767, EPI_ISL_14216769 | | | |

| | | | |
|---|---|---|---|
| see above | Laboratory Medicine, UW Virology | Laboratory Medicine, UW Virology | Sereewit,J., Xie,H., Roychoudhury,P. and Greninger,A. |
| EPI_ISL_14224334 | Genetica Molecular and Subdepartamento de Virologia ISP Chile | Instituto de Salud Publica de Chile | Paulo C. Covarrubias, Andrés E. Castillo, Constanza Campano, Mariela Guajardo, Bárbara Parra, Rodrigo Fasce Pineda, Jorge Fernández |
| EPI_ISL_14244555 | Centers for Disease Control & Prevention (CDC), Division of High Consequence Pathogens and Pathology (DHCPP-PRB) | Centers for Disease Control & Prevention (CDC), Division of High Consequence Pathogens and Pathology (DHCPP-PRB) | Gigante,C., Ventura,J., Seabolt,M.H., Zhao,H., Wilkins,K., McCollum,A., Hutson,C., Davidson,W., Rao,A., Nash,J. and Li,Y. |
| EPI_ISL_14244556 | Centers for Disease Control & Prevention (CDC), Division of High Consequence Pathogens and Pathology (DHCPP-PRB) | Centers for Disease Control & Prevention (CDC), Division of High Consequence Pathogens and Pathology (DHCPP-PRB) | Gigante,C., Ventura,J., Seabolt,M.H., Zhao,H., Wilkins,K., McCollum,A., Hutson,C., Davidson,W., Rao,A., Nash,J., Sheth,M. and Li,Y. |
| EPI_ISL_14244557 | Centers for Disease Control & Prevention (CDC), Division of High Consequence Pathogens and Pathology (DHCPP-PRB) | Centers for Disease Control & Prevention (CDC), Division of High Consequence Pathogens and Pathology (DHCPP-PRB) | Gigante,C., Xia,D., Seabolt,M., Zhao,H., Wilkins,K., McCollum,A., Hutson,C., Davidson,W., Rao,A., Plipat,N. and Li,Y. |
| EPI_ISL_14244558 | Centers for Disease Control & Prevention (CDC), Division of High Consequence Pathogens and Pathology (DHCPP-PRB) | Centers for Disease Control & Prevention (CDC), Division of High Consequence Pathogens and Pathology (DHCPP-PRB) | Gigante,C., Goldoft,M., Seabolt,M., Zhao,H., Wilkins,K., McCollum,A., Hutson,C., Davidson,W., Rao,A., Holshue,M. and Li,Y. |
| EPI_ISL_14244559 | Centers for Disease Control & Prevention (CDC), Division of High Consequence Pathogens and Pathology (DHCPP-PRB) | Centers for Disease Control & Prevention (CDC), Division of High Consequence Pathogens and Pathology (DHCPP-PRB) | Gigante,C., Pavlick,J., Seabolt,M., Zhao,H., Wilkins,K., McCollum,A., Hutson,C., Davidson,W., Rao,A., Parrott,T. and Li,Y. |
| EPI_ISL_14251112 | University of Rochester Medical Center | University of Rochester Medical Center | Andrew Cameron, Mondraya Howard, Joel Maki, Sara Connelly, Kelly Delary, Dwight Hardy |
| EPI_ISL_14254435, EPI_ISL_14254436, EPI_ISL_14254437, EPI_ISL_14254438 | Erasmus Medical Center Department of Virology | Erasmus Medical Center Department of Virology | Bas Oude Munnink, Marjan Boter, Babette Weller, Babs Verstrepen, Richard Molenkamp, Janette Rahamat-Langendoen, Reina Sikkema, Marion Koopmans |
| EPI_ISL_14315314, EPI_ISL_14315315, EPI_ISL_14315316, EPI_ISL_14315318, EPI_ISL_14315319, EPI_ISL_14315320, EPI_ISL_14315321, EPI_ISL_14315322, EPI_ISL_14315323, EPI_ISL_14315324 | Laboratory Medicine, UW Virology | Laboratory Medicine, UW Virology | Sereewit,J., Xie,H., Roychoudhury,P. and Greninger,A.L. |
| EPI_ISL_14326638, EPI_ISL_14326639, EPI_ISL_14326640, EPI_ISL_14326641, EPI_ISL_14326642, EPI_ISL_14326643 | Environmental, Agricultural, and Occupational Health, University of Nebraska Medical Center, 984388 Nebraska Medical Center | Environmental, Agricultural, and Occupational Health, University of Nebraska Medical Center, 984388 Nebraska Medical Center | Tegomoh,B., Cross,S.T., Chapman,R.C., Bernhard,K., McCutchen,E.L., Fauver,J.R., Pratt,C.B., Warden,D.E., Iwen,P.C., Donahue,M. and Wiley,M.R. |
| EPI_ISL_14326644 | Environmental, Agricultural, and Occupational Health, University of Nebraska Medical Center, 984388 Nebraska Medical Center | Environmental, Agricultural, and Occupational Health, University of Nebraska Medical Center, 984388 Nebraska Medical Center | Tegomoh,B., Cross,S.T., Chapman,R.C., Bernhard,K., McCutchen,E.L., Fauver,J.R., Pratt,C.B., Warden,D.E., Iwen,P.C., Donahue,M. and Wiley,M.R |
| EPI_ISL_14355206, EPI_ISL_14355207, EPI_ISL_14355208, EPI_ISL_14355209, EPI_ISL_14355210, EPI_ISL_14355211 | Laboratory Medicine, UW Virology | Laboratory Medicine, UW Virology | Sereewit,J., Xie,H., Roychoudhury,P. and Greninger,A.L. |
| EPI_ISL_14362272 | Centers for Disease Control & Prevention (CDC), Division of High Consequence Pathogens and Pathology (DHCPP-PRB) | Centers for Disease Control & Prevention (CDC), Division of High Consequence Pathogens and Pathology (DHCPP-PRB) | Gigante,C.M., Kubin,G., Seabolt,M.H., Zhao,H., Wilkins,K., McCollum,A., Hutson,C., Davidson,W., Rao,A., White,S.L. and Li,Y. |
| EPI_ISL_14362274, EPI_ISL_14362276 | Centers for Disease Control & Prevention (CDC), Division of High Consequence Pathogens and Pathology (DHCPP-PRB) | Centers for Disease Control & Prevention (CDC), Division of High Consequence Pathogens and Pathology (DHCPP-PRB) | Gigante,C.M., Hughes,S., Seabolt,M.H., Zhao,H., Wilkins,K., McCollum,A., Hutson,C., Davidson,W., Rao,A., Baumgartner,J. and Li,Y. |
| EPI_ISL_14362278 | Centers for Disease Control & Prevention (CDC), Division of High Consequence Pathogens and Pathology (DHCPP-PRB) | Centers for Disease Control & Prevention (CDC), Division of High Consequence Pathogens and Pathology (DHCPP-PRB) | Gigante,C.M., Ghinai,I., Seabolt,M.H., Zhao,H., Wilkins,K., McCollum,A., Hutson,C., Davidson,W., Rao,A., Kerins,J. and Li,Y. |
| EPI_ISL_14362280 | Centers for Disease Control & Prevention (CDC), Division of High Consequence Pathogens and Pathology (DHCPP-PRB) | Centers for Disease Control & Prevention (CDC), Division of High Consequence Pathogens and Pathology (DHCPP-PRB) | Gigante,C.M., Lee,P., Seabolt,M.H., Zhao,H., Wilkins,K., McCollum,A., Hutson,C., Davidson,W., Rao,A., Mendoza,R. and Li,Y. |
| EPI_ISL_14362282, EPI_ISL_14362285 | Centers for Disease Control & Prevention (CDC), Division of High Consequence Pathogens and Pathology (DHCPP-PRB) | Centers for Disease Control & Prevention (CDC), Division of High Consequence Pathogens and Pathology (DHCPP-PRB) | Gigante,C.M., Steidley,B., Seabolt,M.H., Zhao,H., Wilkins,K., McCollum,A., Hutson,C., Davidson,W., Rao,A., Davizon,E.S. and Li,Y. |
| EPI_ISL_14362289 | Centers for Disease Control & Prevention (CDC), Division of High Consequence Pathogens and Pathology (DHCPP-PRB) | Centers for Disease Control & Prevention (CDC), Division of High Consequence Pathogens and Pathology (DHCPP-PRB) | Gigante,C.M., Francis,D., Seabolt,M.H., Zhao,H., Wilkins,K., McCollum,A., Hutson,C., Davidson,W., Rao,A., Escobar,J. and Li,Y. |
| EPI_ISL_14394060 | Ryota Kumagai Tokyo Metropolitan Institute of Public Health, Department of Microbiology | Ryota Kumagai Tokyo Metropolitan Institute of Public Health, Department of Microbiology | Kasuya,F., Negishi,A., Kumagai,R., Hasegawa,M., Fujiwara,T.,Miyake,H., Nagashima,M. and Sadamasu,K. |
| EPI_ISL_14414948 | UMS Parque Industrial Curitiba | Instituto Adolfo Lutz Strategic Laboratory | Claudio Tavares Sacchi, Karoline Rodrigues Campos, Ariadne Ferreira Amarante, Marlon Benedito Nascimento Santos, Alex Domingos Reis, Adriano Abbud, Adriana Bugno |
| EPI_ISL_14415810 | CTA Sao Miguel | Instituto Adolfo Lutz Strategic Laboratory | Claudio Tavares Sacchi, Karoline Rodrigues Campos, Ariadne Ferreira Amarante, Marlon Benedito Nascimento Santos, Alex Domingos Reis, Adriano Abbud, Adriana Bugno |
| EPI_ISL_14439712, EPI_ISL_14439713, EPI_ISL_14439714, EPI_ISL_14439715, EPI_ISL_14439716, EPI_ISL_14439717, EPI_ISL_14439718, EPI_ISL_14439719, EPI_ISL_14439720, EPI_ISL_14439721, EPI_ISL_14439722, EPI_ISL_14439723, EPI_ISL_14439724, EPI_ISL_14439725, EPI_ISL_14439726, EPI_ISL_14439728, EPI_ISL_14439729, EPI_ISL_14439730, EPI_ISL_14439731, EPI_ISL_14439732, EPI_ISL_14439733, EPI_ISL_14439734, EPI_ISL_14439735, EPI_ISL_14439736, EPI_ISL_14439737, EPI_ISL_14439738, EPI_ISL_14439739, EPI_ISL_14439740, EPI_ISL_14439741, EPI_ISL_14439742, EPI_ISL_14439743, EPI_ISL_14439745, EPI_ISL_14439746, EPI_ISL_14439747, EPI_ISL_14439748, EPI_ISL_14439749, EPI_ISL_14439750, EPI_ISL_14439751, EPI_ISL_14439752, EPI_ISL_14439753, EPI_ISL_14439754, EPI_ISL_14439755, EPI_ISL_14439756, EPI_ISL_14439757, EPI_ISL_14439758, EPI_ISL_14439759, EPI_ISL_14439760, EPI_ISL_14439761, EPI_ISL_14439762, EPI_ISL_14439763, EPI_ISL_14439764, EPI_ISL_14439765, EPI_ISL_14439766, EPI_ISL_14439767, EPI_ISL_14439768, EPI_ISL_14439769, EPI_ISL_14439770, EPI_ISL_14439771, EPI_ISL_14439772, EPI_ISL_14439773, EPI_ISL_14439774, EPI_ISL_14439775, EPI_ISL_14439776, EPI_ISL_14439777, EPI_ISL_14439778, EPI_ISL_14439779, EPI_ISL_14439780, EPI_ISL_14439781, EPI_ISL_14439782, EPI_ISL_14439784, EPI_ISL_14439785 | | | |
| see above | Research and Evaluation, UKHSA | Research and Evaluation, UKHSA | Groves,N., Osman,K.L., Lewandowski,K.S., Carter,D.P., Pullan,S.T., Myers,R., Vipond,R. and Chand,M. |
| EPI_ISL_14445098, EPI_ISL_14445101, EPI_ISL_14445102, EPI_ISL_14445103, EPI_ISL_14445109, EPI_ISL_14445116, EPI_ISL_14445118, EPI_ISL_14445119, EPI_ISL_14445120, EPI_ISL_14445121, EPI_ISL_14445122, EPI_ISL_14445123, EPI_ISL_14445124, EPI_ISL_14445125, EPI_ISL_14445126, EPI_ISL_14445127, EPI_ISL_14445128, EPI_ISL_14445129, EPI_ISL_14445130, EPI_ISL_14445131, EPI_ISL_14445132, EPI_ISL_14445133, EPI_ISL_14445134, EPI_ISL_14445135, EPI_ISL_14445136, EPI_ISL_14445137, EPI_ISL_14445138, EPI_ISL_14445139, EPI_ISL_14445140, EPI_ISL_14445141, EPI_ISL_14445146, EPI_ISL_14445147, EPI_ISL_14445150, EPI_ISL_14445152, EPI_ISL_14445153 | | | |
| see above | Laboratorio de Referencia Nacional de Virus Respiratorio. Centro Nacional de Salud Publica. Instituto Nacional de Salud. | Laboratorio de Referencia Nacional de Virus Respiratorio. Centro Nacional de Salud Publica. Instituto Nacional de Salud. | Carlos Padilla Rojas, Veronica Hurtado Vela, Iris Silva Molina, Luren Sevilla Castañeda, Victor Jimenez Vasquez, Orson Mestanza Millones, Luis Barcena Flores, Wendy Lizarraga Olivares, Alicia Nuñez Llanos, Steve Acedo Lazo, Francisco Ascue Orosco, Kelly Izarra Rojas, Princesa Medrano Alhuay, Karla Vasquez Cajachahua, Estela Huaman Angeles, Jorge Giraldo Chavez, Lilian Huarca Balbin, Lisbet Roxana Inga Angulo, Maria Sandra Villar Saavedra, Henri Bailon Calderon, Lely Solari Zerpa, Gloria Arotinco Garayar. Equipo de vigilancia genomica del Instituto Nacional de Salud. |
| EPI_ISL_14445154, EPI_ISL_14445155, EPI_ISL_14445156 | Centre for Biological Threats, Highly Pathogenic Viruses, Robert Koch Institute | Centre for Biological Threats, Highly Pathogenic Viruses, Robert Koch Institute | Brinkmann,A., Kohl,C., Pape,K., Uddin,S., Schrick,L., Michel,J., Stocker,H., Schaade,L. and Nitsche,A. |
| EPI_ISL_14445157, EPI_ISL_14445158, EPI_ISL_14445159, EPI_ISL_14445160, EPI_ISL_14445161, EPI_ISL_14445162, EPI_ISL_14445163, EPI_ISL_14445164 | Centre for Biological Threats, Highly Pathogenic Viruses, Robert Koch Institute | Centre for Biological Threats, Highly Pathogenic Viruses, Robert Koch Institute | Brinkmann,A., Kohl,C., Pape,K., Uddin,S., Schrick,L., Michel,J., Jessen,H., Schaade,L. and Nitsche,A. |
| EPI_ISL_14465517 | Centro de Desenvolvimento Científico e Tecnológico (CDCT), Centro Estadual de Vigilância em Saúde (CEVS) da Secretaria Estadual da Saúde (SES-RS) | Centro de Desenvolvimento Científico e Tecnológico (CDCT), Centro Estadual de Vigilância em Saúde (CEVS) da Secretaria Estadual da Saúde (SES-RS) | Richard Steiner Salvato, Regina Bones Barcellos, Fernanda Marques Godinho |
| EPI_ISL_14467428, EPI_ISL_14467429 | Laboratório Central de Saúde Pública do Amazonas - LACEN-AM | Laboratório de Ecologia de Doenças Transmissíveis na Amazônia, Instituto Leônidas e Maria Deane - Fiocruz Amazônia | Victor Souza, Fernanda Nascimento, Matilde Mejía, Dejanane Silva, Luciana Gonçalves, Tatyana Costa Amorim Ramos, Ana Ruth Lima Arcanjo, Valdinete Nascimento, Felipe Naveca on behalf of the Fiocruz COVID-19 Genomic Surveillance Network |

| | | | |
|---|---|---|---|
| EPI_ISL_14487651 | Centers for Disease Control & Prevention (CDC), Division of High Consequence Pathogens and Pathology (DHCPP-PRB) | Centers for Disease Control & Prevention (CDC), Division of High Consequence Pathogens and Pathology (DHCPP-PRB) | Gigante,C.M., Hughes,S., Seabolt,M.H., Zhao,H., Wilkins,K., McCollum,A., Hutson,C., Davidson,W., Rao,A., Baumgartner,J. and Li,Y. |
| EPI_ISL_14487652 | Centers for Disease Control & Prevention (CDC), Division of High Consequence Pathogens and Pathology (DHCPP-PRB) | Centers for Disease Control & Prevention (CDC), Division of High Consequence Pathogens and Pathology (DHCPP-PRB) | Gigante,C.M., Griffin-Thomas,L., Seabolt,M.H., Zhao,H., Wilkins,K., McCollum,A., Hutson,C., Davidson,W., Rao,A., Crain,J. and Li,Y. |
| EPI_ISL_14487653 | Centers for Disease Control & Prevention (CDC), Division of High Consequence Pathogens and Pathology (DHCPP-PRB) | Centers for Disease Control & Prevention (CDC), Division of High Consequence Pathogens and Pathology (DHCPP-PRB) | Gigante,C.M., Ghinai,I., Seabolt,M.H., Zhao,H., Wilkins,K., McCollum,A., Hutson,C., Davidson,W., Rao,A., Kerins,J. and Li,Y. |
| EPI_ISL_14487654 | Centers for Disease Control & Prevention (CDC), Division of High Consequence Pathogens and Pathology (DHCPP-PRB) | Centers for Disease Control & Prevention (CDC), Division of High Consequence Pathogens and Pathology (DHCPP-PRB) | Gigante,C.M., Steidley,B., Seabolt,M.H., Zhao,H., Wilkins,K., McCollum,A., Hutson,C., Davidson,W., Rao,A., Davizon,E. and Li,Y. |
| EPI_ISL_14487657 | Centers for Disease Control & Prevention (CDC), Division of High Consequence Pathogens and Pathology (DHCPP-PRB) | Centers for Disease Control & Prevention (CDC), Division of High Consequence Pathogens and Pathology (DHCPP-PRB) | Gigante,C.M., Ghinai,I., Seabolt,M.H., Zhao,H., Wilkins,K., McCollum,A., Hutson,C., Davidson,W., Rao,A., Kerins,J. and Li,Y. |
| EPI_ISL_14487659 | Centers for Disease Control & Prevention (CDC), Division of High Consequence Pathogens and Pathology (DHCPP-PRB) | Centers for Disease Control & Prevention (CDC), Division of High Consequence Pathogens and Pathology (DHCPP-PRB) | Gigante,C.M., Hauser,J.R., Seabolt,M.H., Zhao,H., Wilkins,K., McCollum,A., Hutson,C., Davidson,W., Rao,A., Mangla,A. and Li,Y. |
| EPI_ISL_14487660 | Centers for Disease Control & Prevention (CDC), Division of High Consequence Pathogens and Pathology (DHCPP-PRB) | Centers for Disease Control & Prevention (CDC), Division of High Consequence Pathogens and Pathology (DHCPP-PRB) | Gigante,C.M., Ghinai,I., Seabolt,M.H., Zhao,H., Wilkins,K., McCollum,A., Hutson,C., Davidson,W., Rao,A., Kerins,J. and Li,Y. |
| EPI_ISL_14494949 | Division of High-risk Pathogens, Korea Disease Control and Prevention Agency | Division of High-risk Pathogens, Korea Disease Control and Prevention Agency | Rhie,G.-E. |
| EPI_ISL_14515101, EPI_ISL_14515102, EPI_ISL_14515103, EPI_ISL_14515104, EPI_ISL_14515105, EPI_ISL_14515106, EPI_ISL_14515107, EPI_ISL_14515108, EPI_ISL_14515109, EPI_ISL_14515110, EPI_ISL_14515111, EPI_ISL_14515112, EPI_ISL_14515113 | | | |
| see above | Centre for Biological Threats, Highly Pathogenic Viruses, Robert Koch Institute | Centre for Biological Threats, Highly Pathogenic Viruses, Robert Koch Institute | Brinkmann,A., Kohl,C., Pape,K., Uddin,S., Schrick,L., Michel,J., Jessen,H., Schaade,L. and Nitsche,A. |
| EPI_ISL_14515114, EPI_ISL_14515115, EPI_ISL_14515116, EPI_ISL_14515117, EPI_ISL_14515118, EPI_ISL_14515119, EPI_ISL_14515120, EPI_ISL_14515121, EPI_ISL_14515122, EPI_ISL_14515123, EPI_ISL_14515124, EPI_ISL_14515125, EPI_ISL_14515126, EPI_ISL_14515127, EPI_ISL_14515128, EPI_ISL_14515129, EPI_ISL_14515130, EPI_ISL_14515131, EPI_ISL_14515132, EPI_ISL_14515133, EPI_ISL_14515134, EPI_ISL_14515136, EPI_ISL_14515137, EPI_ISL_14515138, EPI_ISL_14515139, EPI_ISL_14515140, EPI_ISL_14515141, EPI_ISL_14515142, EPI_ISL_14515143, EPI_ISL_14515144, EPI_ISL_14515145, EPI_ISL_14515146, EPI_ISL_14515147, EPI_ISL_14515148, EPI_ISL_14515149, EPI_ISL_14515150, EPI_ISL_14515151, EPI_ISL_14515152 | | | |
| see above | Centre for Biological Threats, Highly Pathogenic Viruses, Robert Koch Institute | Centre for Biological Threats, Highly Pathogenic Viruses, Robert Koch Institute | Brinkmann,A., Kohl,C., Pape,K., Uddin,S., Schrick,L., Michel,J., Schaade,L. and Nitsche,A. |
| EPI_ISL_14515153, EPI_ISL_14515154, EPI_ISL_14515155 | Centre for Biological Threats, Highly Pathogenic Viruses, Robert Koch Institute | Centre for Biological Threats, Highly Pathogenic Viruses, Robert Koch Institute | Brinkmann,A., Kohl,C., Pape,K., Uddin,S., Schrick,L., Michel,J., Jessen,H., Schaade,L. and Nitsche,A. |
| EPI_ISL_14515157, EPI_ISL_14515158, EPI_ISL_14515159, EPI_ISL_14515160, EPI_ISL_14515161, EPI_ISL_14515162, EPI_ISL_14515163, EPI_ISL_14515164, EPI_ISL_14515165, EPI_ISL_14515166, EPI_ISL_14515167, EPI_ISL_14515168, EPI_ISL_14515169, EPI_ISL_14515170, EPI_ISL_14515171, EPI_ISL_14515172 | | | |
| see above | Centre for Biological Threats, Highly Pathogenic Viruses, Robert Koch Institute | Centre for Biological Threats, Highly Pathogenic Viruses, Robert Koch Institute | Brinkmann,A., Kohl,C., Pape,K., Uddin,S., Schrick,L., Michel,J., Schaade,L. and Nitsche,A. |
| EPI_ISL_14515201 | Department of Infectious Diseases, National Institute of Health Doutor Ricardo Jorge, Portugal (INSA) | Department of Infectious Diseases, National Institute of Health Doutor Ricardo Jorge, Portugal (INSA) | Isidro,J., Borges,V., Pinto,M., Sobral,D., Santos,J., Nunes,A., Mixao,V., Ferreira,R., Santos,D., Duarte,S., Vieira,L., Borrego,M.J., Nuncio,S., Lopes de Carvalho,I., Pelerito,A., Cordeiro,R. and Gomes,J.P. |
| EPI_ISL_14526939, EPI_ISL_14526942, EPI_ISL_14526944, EPI_ISL_14526945, EPI_ISL_14526948, EPI_ISL_14526949, EPI_ISL_14526950, EPI_ISL_14526952, EPI_ISL_14526953, EPI_ISL_14526955, EPI_ISL_14526956 | | | |
| see above | Connecticut Department of Public Health | Grubaugh Lab - Yale School of Public Health | Nicholas F. G. Chen, Chrispin Chaguza, Kien Pham, Nathan D. Grubaugh, Christina Nishimura, Claire Pearson, Kutluhan Incekara, Jian Ping Huang, Emily Gagnon, Ethan Reever, Jafar Razeq, Anthony Muyombwe, Chantal B. F. Vogels |
| EPI_ISL_14541645, EPI_ISL_14541654 | Public Health Authority of the Slovak Republic | Laboratory of Genomics and Bioinformatics, Comenius University Science Park | Tomáš Szemes, Edita Staroová, Elena Tichá, Lucia Ševíková, Terézia Vrabová, Tatiana Sedláková, Miroslav Böhmer, Jaroslav Budiš, Pavol Mišenko |
| EPI_ISL_14562478, EPI_ISL_14562479, EPI_ISL_14562480, EPI_ISL_14562481, EPI_ISL_14562482, EPI_ISL_14562483, EPI_ISL_14562484, EPI_ISL_14562485, EPI_ISL_14562486, EPI_ISL_14562487, EPI_ISL_14562488, EPI_ISL_14562489, EPI_ISL_14562491, EPI_ISL_14562492, EPI_ISL_14562493, EPI_ISL_14562494, EPI_ISL_14562495, EPI_ISL_14562496, EPI_ISL_14562497, EPI_ISL_14562498, EPI_ISL_14562499, EPI_ISL_14562500, EPI_ISL_14562501, EPI_ISL_14562502 | | | |
| see above | Laboratory Medicine, UW Virology | Laboratory Medicine, UW Virology | Sereewit,J., Xie,H., Roychoudhury,P. and Greninger,A.L. |
| EPI_ISL_14562503, EPI_ISL_14562504, EPI_ISL_14562505, EPI_ISL_14562506, EPI_ISL_14562507, EPI_ISL_14562508, EPI_ISL_14562509, EPI_ISL_14562510, EPI_ISL_14562511, EPI_ISL_14562512 | Centre for Biological Threats, Highly Pathogenic Viruses, Robert Koch Institute | Centre for Biological Threats, Highly Pathogenic Viruses, Robert Koch Institute | Brinkmann,A., Kohl,C., Uddin,S., Pape,K., Schrick,L., Michel,J., Schaade,L. and Nitsche,A. |
| EPI_ISL_14571429 | Hosp. Municipal Dr. Jose de Carvalho Florence | Instituto Adolfo Lutz Strategic Laboratory | Claudio Tavares Sacchi, Karoline Rodrigues Campos, Ariadne Ferreira Amarante, Marlon Benedito Nascimento Santos, Alex Domingos Reis, Adriano Abbud, Adriana Bugno |
| EPI_ISL_14571433 | Casa de Saude Stella Maris | Instituto Adolfo Lutz Strategic Laboratory | Claudio Tavares Sacchi, Karoline Rodrigues Campos, Ariadne Ferreira Amarante, Marlon Benedito Nascimento Santos, Alex Domingos Reis, Adriano Abbud, Adriana Bugno |
| EPI_ISL_14571435 | Secrtetaria Municipal de Saude de Sertaozinho | Instituto Adolfo Lutz Strategic Laboratory | Claudio Tavares Sacchi, Karoline Rodrigues Campos, Ariadne Ferreira Amarante, Marlon Benedito Nascimento Santos, Alex Domingos Reis, Adriano Abbud, Adriana Bugno |
| EPI_ISL_14571439 | Secrtetaria Municipal de Saude de Sata Barbara D Oeste | Instituto Adolfo Lutz Strategic Laboratory | Claudio Tavares Sacchi, Karoline Rodrigues Campos, Ariadne Ferreira Amarante, Marlon Benedito Nascimento Santos, Alex Domingos Reis, Adriano Abbud, Adriana Bugno |
| EPI_ISL_14571441 | Hosp. Municipal Dr. Waldemar Tebaldi | Instituto Adolfo Lutz Strategic Laboratory | Claudio Tavares Sacchi, Karoline Rodrigues Campos, Ariadne Ferreira Amarante, Marlon Benedito Nascimento Santos, Alex Domingos Reis, Adriano Abbud, Adriana Bugno |
| EPI_ISL_14571442 | Instituto de Infectologia Emilio Ribas II Baixada Santista | Instituto Adolfo Lutz Strategic Laboratory | Claudio Tavares Sacchi, Karoline Rodrigues Campos, Ariadne Ferreira Amarante, Marlon Benedito Nascimento Santos, Alex Domingos Reis, Adriano Abbud, Adriana Bugno |
| EPI_ISL_14571444 | UBDS DR. Italo Baruffi Castelo Branco | Instituto Adolfo Lutz Strategic Laboratory | Claudio Tavares Sacchi, Karoline Rodrigues Campos, Ariadne Ferreira Amarante, Marlon Benedito Nascimento Santos, Alex Domingos Reis, Adriano Abbud, Adriana Bugno |
| EPI_ISL_14584274, EPI_ISL_14584275, EPI_ISL_14584277, EPI_ISL_14584279, EPI_ISL_14584281, EPI_ISL_14584282, EPI_ISL_14584283, EPI_ISL_14584284, EPI_ISL_14584286, EPI_ISL_14584287, EPI_ISL_14584289, EPI_ISL_14584290, EPI_ISL_14584291, EPI_ISL_14584292, EPI_ISL_14584293, EPI_ISL_14584294, EPI_ISL_14584295, EPI_ISL_14584296, EPI_ISL_14584297, EPI_ISL_14584298, EPI_ISL_14584299, EPI_ISL_14584300, EPI_ISL_14584302, EPI_ISL_14584303, EPI_ISL_14584304, EPI_ISL_14584305, EPI_ISL_14584306, EPI_ISL_14584308, EPI_ISL_14584309, EPI_ISL_14584310, EPI_ISL_14584311 | | | |
| see above | Laboratorio de Referencia Nacional de Virus Respiratorio. Centro Nacional de Salud Publica. Instituto Nacional de Salud. | Laboratorio de Referencia Nacional de Virus Respiratorio. Centro Nacional de Salud Publica. Instituto Nacional de Salud. | Carlos Padilla Rojas, Veronica Hurtado Vela, Iris Silva Molina, Luren Sevilla Castañeda, Victor Jimenez Vasquez, Orson Mestanza Millones, Luis Barcena Flores, Wendy Lizarraga Olivares, Alicia Nuñez Llanos, Steve Acedo Lazo, Francisco Ascue Orosco, Kelly Izarra Rojas, Princesa Medrano Alhuay, Karla Vasquez Cajachahua, Estela Huaman Angeles, Jorge Giraldo Chavez, Lilian Huarca Balbin, Lisbet Roxana Inga Angulo, Maria Sandra Villar Saavedra, Henri Bailon Calderon, Lely Solari Zerpa, Gloria Arotinco Garayar. Equipo de vigilancia genomica del Instituto Nacional de Salud. |
| EPI_ISL_14586688 | Public Health Authority of the Slovak Republic | Laboratory of Genomics and Bioinformatics, Comenius University Science Park | Tomáš Szemes, Edita Staroová, Elena Tichá, Lucia Ševíková, Terézia Vrabová, Tatiana Sedláková, Miroslav Böhmer, Jaroslav Budiš, Pavol Mišenko |
| EPI_ISL_14587544, EPI_ISL_14587545, EPI_ISL_14587546, EPI_ISL_14587548, EPI_ISL_14587549, EPI_ISL_14587550 | Public Health Agency of Canada, National Microbiology Laboratory | Public Health Agency of Canada, National Microbiology Laboratory | Duggan,A., Hole,D., Yadav,C., Knox,N., Tyler,A., Haidl,E., Chapel,M., Domselaar,G.V., Graham,M., Audet,J., Fernando,L., Hagan,M., Safronetz,D., Leung,A., Peters,G., Go,A., Laminman,V., Kaplen,B., Antonation,K., Griffiths,E., Jolly,G., Charest,H., Levade,I. and Fafard,J. |
| EPI_ISL_14587552, EPI_ISL_14587554, EPI_ISL_14587555, EPI_ISL_14587556, | Centre for Biological Threats, Highly Pathogenic Viruses, Robert Koch Institute | Centre for Biological Threats, Highly Pathogenic Viruses, Robert Koch Institute | Brinkmann,A., Kohl,C., Pape,K., Uddin,S., Schrick,L., Michel,J., Friesen,J., Schaade,L. and Nitsche,A. |

| | | | |
|---|---|---|---|
| EPI_ISL_14587557, EPI_ISL_14587558 | | | |
| EPI_ISL_14594043, EPI_ISL_14594051, EPI_ISL_14594054, EPI_ISL_14594056 | Public Health Agency of Canada, National Microbiology Laboratory | Public Health Agency of Canada, National Microbiology Laboratory | Duggan,A., Hole,D., Yadav,C., Knox,N., Tyler,A., Haidl,E., Chapel,M., Domselaar,G.V., Graham,M., Audet,J., Fernando,L., Antonation,K., Safronetz,D., Hagan,M., Peters,G., Go,A., Laminman,V., Kaplen,B., Leung,A., Griffiths,E., Jolly,G., Eshaghi,A., Gubbay,J.B., Hasso,M., Marchand-Austin,A., Olsha,R. and Patel,S.N. |
| EPI_ISL_14615579 | RSUPN dr. Cipto Mangunkusumo | National Institute of Health Research and Development | Hana Apsari Pawestri, Arie Ardiansyah Nugraha, Fajar Nur Sulistiyohadi, Subangkit, Krisna NA Pangesti, Tze Minn Mak, I Gede Made Wirabrata |
| EPI_ISL_14621526 | Virology, APHP Pitie Salpetriere SU | Virology, APHP Pitie Salpetriere SU | Seang,S., Burrel,S., Todesco,E., Leducq,V., Monsel,G., Le Pluart,D., Cordevant,C., Pourcher,V. and Palich,R. |
| EPI_ISL_14622055 | Instituto de Infectologia Emilio Ribas | Instituto Adolfo Lutz Strategic Laboratory | Claudio Tavares Sacchi, Karoline Rodrigues Campos, Ariadne Ferreira Amarante, Marlon Benedito Nascimento Santos, Alex Domingos Reis, Adriano Abbud, Adriana Bugno |
| EPI_ISL_14622520 | UBS Jovaia | Instituto Adolfo Lutz Strategic Laboratory | Claudio Tavares Sacchi, Karoline Rodrigues Campos, Ariadne Ferreira Amarante, Marlon Benedito Nascimento Santos, Alex Domingos Reis, Adriano Abbud, Adriana Bugno |
| EPI_ISL_14622705 | UBS Jardim Santista | Instituto Adolfo Lutz Strategic Laboratory | Claudio Tavares Sacchi, Karoline Rodrigues Campos, Ariadne Ferreira Amarante, Marlon Benedito Nascimento Santos, Alex Domingos Reis, Adriano Abbud, Adriana Bugno |
| EPI_ISL_14622706 | Centro de Referencia Modulo I SAE II Bauru | Instituto Adolfo Lutz Strategic Laboratory | Claudio Tavares Sacchi, Karoline Rodrigues Campos, Ariadne Ferreira Amarante, Marlon Benedito Nascimento Santos, Alex Domingos Reis, Adriano Abbud, Adriana Bugno |
| EPI_ISL_14622707 | USF Boicucanga I Sao Sebastiao | Instituto Adolfo Lutz Strategic Laboratory | Claudio Tavares Sacchi, Karoline Rodrigues Campos, Ariadne Ferreira Amarante, Marlon Benedito Nascimento Santos, Alex Domingos Reis, Adriano Abbud, Adriana Bugno |
| EPI_ISL_14622913 | Secretaria Municipal de Saude de Caxias do Sul | Instituto Adolfo Lutz Strategic Laboratory | Claudio Tavares Sacchi, Karoline Rodrigues Campos, Ariadne Ferreira Amarante, Marlon Benedito Nascimento Santos, Alex Domingos Reis, Adriano Abbud, Adriana Bugno |
| EPI_ISL_14622953 | Sistema de Vigilancia em Saude Viamao | Instituto Adolfo Lutz Strategic Laboratory | Claudio Tavares Sacchi, Karoline Rodrigues Campos, Ariadne Ferreira Amarante, Marlon Benedito Nascimento Santos, Alex Domingos Reis, Adriano Abbud, Adriana Bugno |
| EPI_ISL_14622960 | Vigilancia Epidemiologica Municipal | Instituto Adolfo Lutz Strategic Laboratory | Claudio Tavares Sacchi, Karoline Rodrigues Campos, Ariadne Ferreira Amarante, Marlon Benedito Nascimento Santos, Alex Domingos Reis, Adriano Abbud, Adriana Bugno |
| EPI_ISL_14623175 | Centro de Referencia em Especialidades Central Rib Preto | Instituto Adolfo Lutz Strategic Laboratory | Claudio Tavares Sacchi, Karoline Rodrigues Campos, Ariadne Ferreira Amarante, Marlon Benedito Nascimento Santos, Alex Domingos Reis, Adriano Abbud, Adriana Bugno |
| EPI_ISL_14623523 | Laboratorio Municipal de Piracicaba | Instituto Adolfo Lutz Strategic Laboratory | Claudio Tavares Sacchi, Karoline Rodrigues Campos, Ariadne Ferreira Amarante, Marlon Benedito Nascimento Santos, Alex Domingos Reis, Adriano Abbud, Adriana Bugno |
| EPI_ISL_14623704 | Unidade Basica de Saude Esplanada | Instituto Adolfo Lutz Strategic Laboratory | Claudio Tavares Sacchi, Karoline Rodrigues Campos, Ariadne Ferreira Amarante, Marlon Benedito Nascimento Santos, Alex Domingos Reis, Adriano Abbud, Adriana Bugno |
| EPI_ISL_14624411 | Hospital Albert Sabin Atibaia | Instituto Adolfo Lutz Strategic Laboratory | Claudio Tavares Sacchi, Karoline Rodrigues Campos, Ariadne Ferreira Amarante, Marlon Benedito Nascimento Santos, Alex Domingos Reis, Adriano Abbud, Adriana Bugno |
| EPI_ISL_14624610 | USAFA Forte | Instituto Adolfo Lutz Strategic Laboratory | Claudio Tavares Sacchi, Karoline Rodrigues Campos, Ariadne Ferreira Amarante, Marlon Benedito Nascimento Santos, Alex Domingos Reis, Adriano Abbud, Adriana Bugno |
| EPI_ISL_14624698 | Centro de Referencia em AIDS SECRAIDS | Instituto Adolfo Lutz Strategic Laboratory | Claudio Tavares Sacchi, Karoline Rodrigues Campos, Ariadne Ferreira Amarante, Marlon Benedito Nascimento Santos, Alex Domingos Reis, Adriano Abbud, Adriana Bugno |
| EPI_ISL_14624832 | Servico de Vigilancia Epidemiologica e de Zoonoses do Guaruja | Instituto Adolfo Lutz Strategic Laboratory | Claudio Tavares Sacchi, Karoline Rodrigues Campos, Ariadne Ferreira Amarante, Marlon Benedito Nascimento Santos, Alex Domingos Reis, Adriano Abbud, Adriana Bugno |
| EPI_ISL_14625156 | Secretaria Municipal de Saude de Suzano | Instituto Adolfo Lutz Strategic Laboratory | Claudio Tavares Sacchi, Karoline Rodrigues Campos, Ariadne Ferreira Amarante, Marlon Benedito Nascimento Santos, Alex Domingos Reis, Adriano Abbud, Adriana Bugno |
| EPI_ISL_14625157 | PSF Vila Nossa Senhora de Fatima Fartura | Instituto Adolfo Lutz Strategic Laboratory | Claudio Tavares Sacchi, Karoline Rodrigues Campos, Ariadne Ferreira Amarante, Marlon Benedito Nascimento Santos, Alex Domingos Reis, Adriano Abbud, Adriana Bugno |
| EPI_ISL_14625190 | Ambulatorio de Atendimentode DST de Guariba | Instituto Adolfo Lutz Strategic Laboratory | Claudio Tavares Sacchi, Karoline Rodrigues Campos, Ariadne Ferreira Amarante, Marlon Benedito Nascimento Santos, Alex Domingos Reis, Adriano Abbud, Adriana Bugno |
| EPI_ISL_14625230 | UBS Centro Clair Aparecida Pavan | Instituto Adolfo Lutz Strategic Laboratory | Claudio Tavares Sacchi, Karoline Rodrigues Campos, Ariadne Ferreira Amarante, Marlon Benedito Nascimento Santos, Alex Domingos Reis, Adriano Abbud, Adriana Bugno |
| EPI_ISL_14625256 | UMS Campina do Siqueira | Instituto Adolfo Lutz Strategic Laboratory | Claudio Tavares Sacchi, Karoline Rodrigues Campos, Ariadne Ferreira Amarante, Marlon Benedito Nascimento Santos, Alex Domingos Reis, Adriano Abbud, Adriana Bugno |
| EPI_ISL_14625282 | Hospital Edmundo Vasconcelos | Instituto Adolfo Lutz Strategic Laboratory | Claudio Tavares Sacchi, Karoline Rodrigues Campos, Ariadne Ferreira Amarante, Marlon Benedito Nascimento Santos, Alex Domingos Reis, Adriano Abbud, Adriana Bugno |
| EPI_ISL_14666780 | Public Health Authority of the Slovak Republic | Laboratory of Genomics and Bioinformatics, Comenius University Science Park | Tomáš Szemes, Edita Staroová, Elena Tichá, Lucia Ševíková, Terézia Vrabová, Tatiana Sedláková, Miroslav Böhmer, Jaroslav Budiš, Pavol Mišenko |
| EPI_ISL_14699907, EPI_ISL_14699908, EPI_ISL_14699909, EPI_ISL_14699910 | Centers for Disease Control & Prevention (CDC), Division of High Consequence Pathogens and Pathology (DHCPP-PRB) | Centers for Disease Control & Prevention (CDC), Division of High Consequence Pathogens and Pathology (DHCPP-PRB) | Gigante,C.M., Lee,P., Zhao,H., Batra,D., Hetrick,E.E., Howard,D.T., Kovar,L., Seabolt,M.H., Weigand,M.R., Burroughs,M.S., Lee,J., Wilkins,K., McCollum,A., Hutson,C., Davidson,W., Rao,A., Mendoza,R. and Li,Y. |
| EPI_ISL_14699911, EPI_ISL_14699912, EPI_ISL_14699913, EPI_ISL_14699914, EPI_ISL_14699915 | Centers for Disease Control & Prevention (CDC), Division of High Consequence Pathogens and Pathology (DHCPP-PRB) | Centers for Disease Control & Prevention (CDC), Division of High Consequence Pathogens and Pathology (DHCPP-PRB) | Gigante,C.M., Ghinai,I., Zhao,H., Batra,D., Hetrick,E.E., Howard,D.T., Kovar,L., Seabolt,M.H., Weigand,M.R., Burroughs,M.S., Lee,J., Wilkins,K., McCollum,A., Hutson,C., Davidson,W., Rao,A., Kerins,J. and Li,Y. |
| EPI_ISL_14699916 | Centers for Disease Control & Prevention (CDC), Division of High Consequence Pathogens and Pathology (DHCPP-PRB) | Centers for Disease Control & Prevention (CDC), Division of High Consequence Pathogens and Pathology (DHCPP-PRB) | Gigante,C.M., Kubin,G., Zhao,H., Batra,D., Hetrick,E.E., Howard,D.T., Kovar,L., Seabolt,M.H., Weigand,M.R., Burroughs,M.S., Lee,J., Wilkins,K., McCollum,A., Hutson,C., Davidson,W., Rao,A., White,S.L. and Li,Y. |
| EPI_ISL_14699917 | Centers for Disease Control & Prevention (CDC), Division of High Consequence Pathogens and Pathology (DHCPP-PRB) | Centers for Disease Control & Prevention (CDC), Division of High Consequence Pathogens and Pathology (DHCPP-PRB) | Gigante,C.M., Winter,K., Zhao,H., Batra,D., Hetrick,E.E., Howard,D.T., Kovar,L., Seabolt,M.H., Weigand,M.R., Burroughs,M.S., Lee,J., Wilkins,K., McCollum,A., Hutson,C., Davidson,W., Rao,A., Arora,V. and Li,Y. |
| EPI_ISL_14699918 | Centers for Disease Control & Prevention (CDC), Division of High Consequence Pathogens and Pathology (DHCPP-PRB) | Centers for Disease Control & Prevention (CDC), Division of High Consequence Pathogens and Pathology (DHCPP-PRB) | Gigante,C.M., Hughes,S., Zhao,H., Batra,D., Hetrick,E.E., Howard,D.T., Kovar,L., Seabolt,M.H., Weigand,M.R., Burroughs,M.S., Lee,J., Wilkins,K., McCollum,A., Hutson,C., Davidson,W., Rao,A., Baumgartner,J. and Li,Y. |
| EPI_ISL_14699919 | Centers for Disease Control & Prevention (CDC), Division of High Consequence Pathogens and Pathology (DHCPP-PRB) | Centers for Disease Control & Prevention (CDC), Division of High Consequence Pathogens and Pathology (DHCPP-PRB) | Gigante,C.M., Ghinai,I., Zhao,H., Batra,D., Hetrick,E.E., Howard,D.T., Kovar,L., Seabolt,M.H., Weigand,M.R., Burroughs,M.S., Lee,J., Wilkins,K., McCollum,A., Hutson,C., Davidson,W., Rao,A., Kerins,J. and Li,Y. |
| EPI_ISL_14699920, EPI_ISL_14699921 | Centers for Disease Control & Prevention (CDC), Division of High Consequence Pathogens and Pathology (DHCPP-PRB) | Centers for Disease Control & Prevention (CDC), Division of High Consequence Pathogens and Pathology (DHCPP-PRB) | Gigante,C.M., Iwen,P.C., Zhao,H., Batra,D., Hetrick,E.E., Howard,D.T., Kovar,L., Seabolt,M.H., Weigand,M.R., Burroughs,M.S., Lee,J., Wilkins,K., McCollum,A., Hutson,C., Davidson,W., Rao,A., Donahue,M. and Li,Y. |
| EPI_ISL_14699922, EPI_ISL_14699923, EPI_ISL_14699925 | Centers for Disease Control & Prevention (CDC), Division of High Consequence Pathogens and Pathology (DHCPP-PRB) | Centers for Disease Control & Prevention (CDC), Division of High Consequence Pathogens and Pathology (DHCPP-PRB) | Gigante,C.M., Hughes,S., Zhao,H., Batra,D., Hetrick,E.E., Howard,D.T., Kovar,L., Seabolt,M.H., Weigand,M.R., Burroughs,M.S., Lee,J., Wilkins,K., McCollum,A., Hutson,C., Davidson,W., Rao,A., Baumgartner,J. and Li,Y. |
| EPI_ISL_14699926 | Centers for Disease Control & Prevention (CDC), Division of High Consequence Pathogens and Pathology (DHCPP-PRB) | Centers for Disease Control & Prevention (CDC), Division of High Consequence Pathogens and Pathology (DHCPP-PRB) | Gigante,C.M., Griffin-Thomas,L., Zhao,H., Batra,D., Hetrick,E.E., Howard,D.T., Kovar,L., Seabolt,M.H., Weigand,M.R., Burroughs,M.S., Lee,J., Wilkins,K., McCollum,A., Hutson,C., Davidson,W., Rao,A., Crain,J. and Li,Y. |

EPI_ISL_14699927, EPI_ISL_14699928, EPI_ISL_14699929, EPI_ISL_14699930, EPI_ISL_14699931, EPI_ISL_14699932, EPI_ISL_14699933, EPI_ISL_14699934, EPI_ISL_14699935, EPI_ISL_14699936, EPI_ISL_14699937, EPI_ISL_14699938, EPI_ISL_14699939, EPI_ISL_14699940, EPI_ISL_14699941, EPI_ISL_14699942,

| | | | |
|---|---|---|---|
| EPI_ISL_14699943, EPI_ISL_14699944, EPI_ISL_14699945, EPI_ISL_14699946, EPI_ISL_14699947, EPI_ISL_14699948, EPI_ISL_14699949, EPI_ISL_14699950, EPI_ISL_14699951, EPI_ISL_14699952, EPI_ISL_14699953, EPI_ISL_14699954, EPI_ISL_14699955, EPI_ISL_14699957, EPI_ISL_14699958 | | | |
| see above | Laboratory Medicine, UW Virology | Laboratory Medicine, UW Virology | Sereewit,J., Xie,H., Roychoudhury,P. and Greninger,A.L. |
| EPI_ISL_14721255, EPI_ISL_14721259, EPI_ISL_14721262, EPI_ISL_14721265 | National Public Health Laboratory, National Centre for Infectious Diseases | National Public Health Laboratory, National Centre for Infectious Diseases | Yichen Ding, Benny Yeo, Daniel Lim, Zhenyang Zhou, Royce Ang, Samuel Loo, Lin Cui, Raymond Tzer Pin Lin |
| EPI_ISL_14736400, EPI_ISL_14736403 | California Department of Public Health | California Department of Public Health | Viral and Rickettsial Disease Laboratory |
| EPI_ISL_14752090, EPI_ISL_14752091, EPI_ISL_14752093, EPI_ISL_14752094, EPI_ISL_14752096 | Environmental, Agricultural, and Occupational Health, University of Nebraska Medical Center | Environmental, Agricultural, and Occupational Health, University of Nebraska Medical Center | Tegomoh,B., Cross,S.T., Chapman,R.C., Bernhard,K., McCutchen,E.L., Fauver,J.R., Pratt,C.B., Warden,D.E., Iwen,P.C., Donahue,M. and Wiley,M.R. |
| EPI_ISL_14752216 | Department of Infectious Diseases, National Institute of Health Doutor Ricardo Jorge (INSA) | Department of Infectious Diseases, National Institute of Health Doutor Ricardo Jorge (INSA) | Isidro,J., Borges,V., Pinto,M., Sobral,D., Santos,J., Nunes,A., Mixao,V., Ferreira,R., Santos,D., Duarte,S., Vieira,L., Borrego,M.J., Nuncio,S., Lopes de Carvalho,I., Pelerito,A., Cordeiro,R. and Gomes,J.P. |
| EPI_ISL_14752257, EPI_ISL_14752259, EPI_ISL_14752260, EPI_ISL_14752261, EPI_ISL_14752262, EPI_ISL_14752263 | Laboratory Medicine, UW Virology | Laboratory Medicine, UW Virology | Sereewit,J., Xie,H., Roychoudhury,P. and Greninger,A.L. |
| EPI_ISL_14752264, EPI_ISL_14752265, EPI_ISL_14752267, EPI_ISL_14752269, EPI_ISL_14752270, EPI_ISL_14752272, EPI_ISL_14752274, EPI_ISL_14752276, EPI_ISL_14752278, EPI_ISL_14752280, EPI_ISL_14752282 | | | |
| see above | Centre for Biological Threats, Highly Pathogenic Viruses, Robert Koch Institute | Centre for Biological Threats, Highly Pathogenic Viruses, Robert Koch Institute | Brinkmann,A., Kohl,C., Uddin,S., Pape,K., Schrick,L., Michel,J., Schaade,L. and Nitsche,A. |
| EPI_ISL_14752284 | Research and Evaluation, UKHSA | Research and Evaluation, UKHSA | Grove,N., Osman,K.L., Lewandowski,K.S., Carter,D.P., Pullan,S.T., Myers,R., Vipond,R. and Chand,M. |
| EPI_ISL_14752290 | Research and Evaluation, UKHSA | Research and Evaluation, UKHSA | Groves,N., Osman,K.L., Lewandowski,K.S., Carter,D.P., Pullan,S.T., Myers,R., Vipond,R. and Chand,M. |
| EPI_ISL_14752293 | Medical Microbiology & Infection Prevention, Amsterdam Medical Centres location AMC | Medical Microbiology & Infection Prevention, Amsterdam Medical Centres location AMC | Welkers,M., Jonges,M., de Regt,M., Ooijevaar,R. and Wagemakers,A. |
| EPI_ISL_14772912 | USF Jardim Oratorio | Instituto Adolfo Lutz Strategic Laboratory | Claudio Tavares Sacchi, Karoline Rodrigues Campos, Ariadne Ferreira Amarante, Marlon Benedito Nascimento Santos, Alex Domingos Reis, Adriano Abbud, Adriana Bugno |
| EPI_ISL_14772913 | Vigilancia Epidemiologica Jardinopolis - SP | Instituto Adolfo Lutz Strategic Laboratory | Claudio Tavares Sacchi, Karoline Rodrigues Campos, Ariadne Ferreira Amarante, Marlon Benedito Nascimento Santos, Alex Domingos Reis, Adriano Abbud, Adriana Bugno |
| EPI_ISL_14772914 | Pronto Atendimento Infantil e entral de Quimioterapia SJrpreto | Instituto Adolfo Lutz Strategic Laboratory | Claudio Tavares Sacchi, Karoline Rodrigues Campos, Ariadne Ferreira Amarante, Marlon Benedito Nascimento Santos, Alex Domingos Reis, Adriano Abbud, Adriana Bugno |
| EPI_ISL_14773001 | CEDIC CTA | Instituto Adolfo Lutz Strategic Laboratory | Claudio Tavares Sacchi, Karoline Rodrigues Campos, Ariadne Ferreira Amarante, Marlon Benedito Nascimento Santos, Alex Domingos Reis, Adriano Abbud, Adriana Bugno |
| EPI_ISL_14783237 | Sicilian Regional Laboratory - AOUP "P. Giaccone" - University of Palermo | Sicilian Regional Laboratory - AOUP "P. Giaccone" - University of Palermo | Fabio Tramuto, Carmelo Massimo Maida, Giulia Randazzo, Valeria Guzzetta, Walter Mazzucco, Giorgio Graziano, Vincenzo Restivo, Claudio Costantino, Francesco Vitale |
| EPI_ISL_14793992 | Erasmus Medical Center Department of Virology | Erasmus Medical Center Department of Virology | Bas Oude Munnink, Leonard Schuele, Marjan Boter, Babette Weller, Babs Verstrepen, Richard Molenkamp, Janette Rahamat-Langendoen, Reina Sikkema, Marion Koopmans |
| EPI_ISL_14804638, EPI_ISL_14804639, EPI_ISL_14804640, EPI_ISL_14804641, EPI_ISL_14804642, EPI_ISL_14804643, EPI_ISL_14804644, EPI_ISL_14804645, EPI_ISL_14804646, EPI_ISL_14804647 | Nebraska Public Health Laboratory | University of Nebraska Medical Center, Oklahoma Pathogen Genomics Consortium | Chapman,R.C., Bernhard,K., McCutchen,E.L., Fauver,J.R., O'Dell,J.X., Mannell,M., Wiley,M.R., Cross,S.T. |
| EPI_ISL_14809096 | AMA Capao Redondo | Instituto Adolfo Lutz Strategic Laboratory | Claudio Tavares Sacchi, Karoline Rodrigues Campos, Ariadne Ferreira Amarante, Marlon Benedito Nascimento Santos, Alex Domingos Reis, Adriano Abbud, Adriana Bugno |
| EPI_ISL_14809098 | Laboratorio Municipal de Piracicaba | Instituto Adolfo Lutz Strategic Laboratory | Claudio Tavares Sacchi, Karoline Rodrigues Campos, Ariadne Ferreira Amarante, Marlon Benedito Nascimento Santos, Alex Domingos Reis, Adriano Abbud, Adriana Bugno |
| EPI_ISL_14809099 | Centro de Saude Gabriel de Lara | Instituto Adolfo Lutz Strategic Laboratory | Claudio Tavares Sacchi, Karoline Rodrigues Campos, Ariadne Ferreira Amarante, Marlon Benedito Nascimento Santos, Alex Domingos Reis, Adriano Abbud, Adriana Bugno |
| EPI_ISL_14809100 | Secretaria Municipal da Saude de Joanopolis | Instituto Adolfo Lutz Strategic Laboratory | Claudio Tavares Sacchi, Karoline Rodrigues Campos, Ariadne Ferreira Amarante, Marlon Benedito Nascimento Santos, Alex Domingos Reis, Adriano Abbud, Adriana Bugno |
| EPI_ISL_14810370, EPI_ISL_14810404, EPI_ISL_14810405, EPI_ISL_14810406, EPI_ISL_14810407 | Erasmus Medical Center Department of Virology | Erasmus Medical Center Department of Virology | Leonard Schuele, Bas Oude Munnink, Marjan Boter, Babette Weller, Babs Verstrepen, Richard Molenkamp, Janette Rahamat-Langendoen, Reina Sikkema, Marion Koopmans |
| EPI_ISL_14818585 | Los Angeles County Public Health Laboratories | Los Angeles County Public Health Laboratories | P. Hemarajata et al. |
| EPI_ISL_14818783, EPI_ISL_14818784, EPI_ISL_14818785, EPI_ISL_14818787, EPI_ISL_14818788, EPI_ISL_14818789, EPI_ISL_14818790, EPI_ISL_14818791, EPI_ISL_14818792, EPI_ISL_14818793, EPI_ISL_14818794, EPI_ISL_14818795, EPI_ISL_14818796, EPI_ISL_14818797, EPI_ISL_14818798, EPI_ISL_14818800, EPI_ISL_14818801, EPI_ISL_14818802, EPI_ISL_14818803, EPI_ISL_14818804, EPI_ISL_14818805, EPI_ISL_14818806, EPI_ISL_14818807, EPI_ISL_14818808, EPI_ISL_14818810, EPI_ISL_14818811, EPI_ISL_14818812, EPI_ISL_14818813, EPI_ISL_14818814, EPI_ISL_14818815, EPI_ISL_14818817, EPI_ISL_14818818, EPI_ISL_14818819, EPI_ISL_14818820, EPI_ISL_14818821, EPI_ISL_14818822 | | | |
| see above | Laboratorio de Referencia Nacional de Virus Inmunoprevenibles. Centro Nacional de Salud Publica. Instituto Nacional de Salud. | Laboratorio de Referencia Nacional de Virus Inmunoprevenibles. Centro Nacional de Salud Publica. Instituto Nacional de Salud. | Carlos Padilla Rojas, Veronica Hurtado Vela, Iris Silva Molina, Luren Sevilla Castañeda, Victor Jimenez Vasquez, Luis Barcena Flores, Alicia Nuñez Llanos, Kelly Izarra Rojas, Karla Vasquez Cajachahua, Estela Huaman Angeles, Jorge Giraldo Chavez, Lilian Huarca Balbin, Maria Sandra Villar Saavedra, Henri Bailon Calderon, Lely Solari Zerpa, Gloria Arotinco Garayar. Equipo de vigilancia genomica del Instituto Nacional de Salud. |
| EPI_ISL_14842151 | Division of High Consequence Pathogens and Pathology (DHCPP)-PRB, CDC | Division of High Consequence Pathogens and Pathology (DHCPP)-PRB, CDC | Gigante,C.M., Hauser,J.R., Zhao,H., Batra,D., Hetrick,E.E., Howard,D.T., Kovar,L., Seabolt,M.H., Weigand,M.R., Burroughs,M., Lee,J., Wilkins,K., McCollum,A., Hutson,C., Davidson,W., Rao,A., Mangla,A. and Li,Y. |
| EPI_ISL_14842152 | Division of High Consequence Pathogens and Pathology (DHCPP)-PRB, CDC | Division of High Consequence Pathogens and Pathology (DHCPP)-PRB, CDC | Gigante,C.M., Pavlick,J., Zhao,H., Batra,D., Hetrick,E.E., Howard,D.T., Kovar,L., Seabolt,M.H., Weigand,M.R., Burroughs,M., Lee,J., Wilkins,K., McCollum,A., Hutson,C., Davidson,W., Rao,A., Parrott,T. and Li,Y. |
| EPI_ISL_14842153, EPI_ISL_14842154 | Division of High Consequence Pathogens and Pathology (DHCPP)-PRB, CDC | Division of High Consequence Pathogens and Pathology (DHCPP)-PRB, CDC | Gigante,C.M., Hauser,J.R., Zhao,H., Batra,D., Hetrick,E.E., Howard,D.T., Kovar,L., Seabolt,M.H., Weigand,M.R., Burroughs,M., Lee,J., Wilkins,K., McCollum,A., Hutson,C., Davidson,W., Rao,A., Mangla,A. and Li,Y. |
| EPI_ISL_14842155, EPI_ISL_14842156, EPI_ISL_14842157 | Division of High Consequence Pathogens and Pathology (DHCPP)-PRB, CDC | Division of High Consequence Pathogens and Pathology (DHCPP)-PRB, CDC | Gigante,C.M., Pavlick,J., Zhao,H., Batra,D., Hetrick,E.E., Howard,D.T., Kovar,L., Seabolt,M.H., Weigand,M.R., Burroughs,M., Lee,J., Wilkins,K., McCollum,A., Hutson,C., Davidson,W., Rao,A., Parrott,T. and Li,Y. |
| EPI_ISL_14842158 | Division of High Consequence Pathogens and Pathology (DHCPP)-PRB, CDC | Division of High Consequence Pathogens and Pathology (DHCPP)-PRB, CDC | Gigante,C.M., Hughes,S., Zhao,H., Batra,D., Hetrick,E.E., Howard,D.T., Kovar,L., Seabolt,M.H., Weigand,M.R., Burroughs,M., Lee,J., Wilkins,K., McCollum,A., Hutson,C., Davidson,W., Rao,A., Baumgartner,J. and Li,Y. |
| EPI_ISL_14842159 | Division of High Consequence Pathogens and Pathology (DHCPP)-PRB, CDC | Division of High Consequence Pathogens and Pathology (DHCPP)-PRB, CDC | Gigante,C.M., Johnson,S., Zhao,H., Batra,D., Hetrick,E.E., Howard,D.T., Kovar,L., Seabolt,M.H., Weigand,M.R., Burroughs,M., Lee,J., Wilkins,K., McCollum,A., Hutson,C., Davidson,W., Rao,A., Riner,D. and Li,Y. |
| EPI_ISL_14842160 | Division of High Consequence Pathogens and Pathology (DHCPP)-PRB, CDC | Division of High Consequence Pathogens and Pathology (DHCPP)-PRB, CDC | Gigante,C.M., Manuzak,A., Zhao,H., Batra,D., Hetrick,E.E., Howard,D.T., Kovar,L., Seabolt,M.H., Weigand,M.R., Burroughs,M., Lee,J., Wilkins,K., McCollum,A., Hutson,C., Davidson,W., Rao,A., Gose,R. and Li,Y. |
| EPI_ISL_14842161, EPI_ISL_14842162 | Division of High Consequence Pathogens and Pathology (DHCPP)-PRB, CDC | Division of High Consequence Pathogens and Pathology (DHCPP)-PRB, CDC | Gigante,C.M., Kubin,G., Zhao,H., Batra,D., Hetrick,E.E., Howard,D.T., Kovar,L., Seabolt,M.H., Weigand,M.R., Burroughs,M., Lee,J., Wilkins,K., McCollum,A., Hutson,C., Davidson,W., Rao,A., White,S.L. and Li,Y. |

| | | | |
|---|---|---|---|
| EPI_ISL_14842163 | Division of High Consequence Pathogens and Pathology (DHCPP)-PRB, CDC | Division of High Consequence Pathogens and Pathology (DHCPP)-PRB, CDC | Gigante,C.M., Ghinai,I., Zhao,H., Batra,D., Hetrick,E.E., Howard,D.T., Kovar,L., Seabolt,M.H., Weigand,M.R., Burroughs,M., Lee,J., Wilkins,K., McCollum,A., Hutson,C., Davidson,W., Rao,A., Kerins,J. and Li,Y. |
| EPI_ISL_14842164, EPI_ISL_14842165 | Division of High Consequence Pathogens and Pathology (DHCPP)-PRB, CDC | Division of High Consequence Pathogens and Pathology (DHCPP)-PRB, CDC | Gigante,C.M., Griffin-Thomas,L., Zhao,H., Batra,D., Hetrick,E.E., Howard,D.T., Kovar,L., Seabolt,M.H., Weigand,M.R., Burroughs,M., Lee,J., Wilkins,K., McCollum,A., Hutson,C., Davidson,W., Rao,A., Crain,J. and Li,Y. |
| EPI_ISL_14842166 | Division of High Consequence Pathogens and Pathology (DHCPP)-PRB, CDC | Division of High Consequence Pathogens and Pathology (DHCPP)-PRB, CDC | Gigante,C.M., Xia,D., Zhao,H., Batra,D., Hetrick,E.E., Howard,D.T., Kovar,L., Seabolt,M.H., Weigand,M.R., Burroughs,M., Lee,J., Wilkins,K., McCollum,A., Hutson,C., Davidson,W., Rao,A., Plipat,N. and Li,Y. |
| EPI_ISL_14842167, EPI_ISL_14842168 | Division of High Consequence Pathogens and Pathology (DHCPP)-PRB, CDC | Division of High Consequence Pathogens and Pathology (DHCPP)-PRB, CDC | Gigante,C.M., Griffin-Thomas,L., Zhao,H., Batra,D., Hetrick,E.E., Howard,D.T., Kovar,L., Seabolt,M.H., Weigand,M.R., Burroughs,M., Lee,J., Wilkins,K., McCollum,A., Hutson,C., Davidson,W., Rao,A., Crain,J. and Li,Y. |
| EPI_ISL_14863048, EPI_ISL_14863050, EPI_ISL_14863051, EPI_ISL_14863052, EPI_ISL_14863053, EPI_ISL_14863055, EPI_ISL_14863057, EPI_ISL_14863059 | MEPHI, IHU - Mediterranee Infection | MEPHI, IHU - Mediterranee Infection | Colson,P. |
| EPI_ISL_14865785 | UBS J COPA | Instituto Adolfo Lutz Strategic Laboratory | Claudio Tavares Sacchi, Karoline Rodrigues Campos, Ariadne Ferreira Amarante, Marlon Benedito Nascimento Santos, Alex Domingos Reis, Adriano Abbud, Adriana Bugno |
| EPI_ISL_14866481 | PR S da Familia Unidade de Saude Adalberto Rocha | Instituto Adolfo Lutz Strategic Laboratory | Claudio Tavares Sacchi, Karoline Rodrigues Campos, Ariadne Ferreira Amarante, Marlon Benedito Nascimento Santos, Alex Domingos Reis, Adriano Abbud, Adriana Bugno |
| EPI_ISL_14866751 | Pronto Socorro da Vila Dirce | Instituto Adolfo Lutz Strategic Laboratory | Claudio Tavares Sacchi, Karoline Rodrigues Campos, Ariadne Ferreira Amarante, Marlon Benedito Nascimento Santos, Alex Domingos Reis, Adriano Abbud, Adriana Bugno |
| EPI_ISL_14866752 | Secretaria Municipal de Saude Sao Carlos | Instituto Adolfo Lutz Strategic Laboratory | Claudio Tavares Sacchi, Karoline Rodrigues Campos, Ariadne Ferreira Amarante, Marlon Benedito Nascimento Santos, Alex Domingos Reis, Adriano Abbud, Adriana Bugno |
| EPI_ISL_14910864, EPI_ISL_14910865, EPI_ISL_14910866, EPI_ISL_14910867, EPI_ISL_14910868, EPI_ISL_14910869, EPI_ISL_14910870, EPI_ISL_14910871, EPI_ISL_14910872, EPI_ISL_14910874, EPI_ISL_14910875, EPI_ISL_14910877, EPI_ISL_14910878, EPI_ISL_14910879, EPI_ISL_14910880, EPI_ISL_14910881, EPI_ISL_14910883, EPI_ISL_14910884, EPI_ISL_14910885 | | | |
| see above | Laboratory Medicine, UW Virology | Laboratory Medicine, UW Virology | Sereewit,J., Xie,H., Roychoudhury,P. and Greninger,A.L. |
| EPI_ISL_14910886 | Research and Evaluation, UKHSA | Research and Evaluation, UKHSA | Burton,J., Easterbrook,L., Drinkwater,E., Groves,N., Osman,K.L., Lewandowski,K.S., Carter,D., Pullan,S.T., Myers,R., Vipond,R. and Chand,M. |
| EPI_ISL_14917581 | Los Angeles County Public Health Laboratories | Los Angeles County Public Health Laboratories | P. Hemarajata et al. |
| EPI_ISL_14923900, EPI_ISL_14923901, EPI_ISL_14923902, EPI_ISL_14923903, EPI_ISL_14923904 | Research and Evaluation, UKHSA | Research and Evaluation, UKHSA | Groves,N., Osman,K.L., Lewandowski,K.S., Carter,D.P., Pullan,S.T., Myers,R., Vipond,R. and Chand,M. |
| EPI_ISL_14934116 | Medical Center of Vienna Center for Virology | Medical University of Vienna Center for Virology | Jeremy V. Camp, Monika Redlberger-Fritz, Stephan W. Aberle |
| EPI_ISL_14934140 | Center for Virology Medical Unviersity of Vienna | Medical University of Vienna Center for Virology | Jeremy V. Camp, Monika Redlberger-Fritz, Stephan W. Aberle |
| EPI_ISL_14934382 | Medical University of Vienna Center for Virology | Medical University of Vienna Center for Virology | Jeremy V. Camp, Monika Redlberger-Fritz, Stephan W. Aberle |
| EPI_ISL_14934478 | Medical University of Vienna Center for Virology | Medical University of Vienna Center for Virology | Jeremy V. Camp, Monika Redlberg-Fritz, Stephan W. Aberle |
| EPI_ISL_14934480, EPI_ISL_14934481, EPI_ISL_14934483, EPI_ISL_14934484, EPI_ISL_14934485, EPI_ISL_14934486, EPI_ISL_14934488, EPI_ISL_14934489, EPI_ISL_14934490, EPI_ISL_14934492, EPI_ISL_14934493, EPI_ISL_14934494, EPI_ISL_14934495 | | | |
| see above | Research and Evaluation, UKHSA | Research and Evaluation, UKHSA | Groves,N., Osman,K.L., Lewandowski,K.S., Carter,D.P., Pullan,S.T., Myers,R., Vipond,R. and Chand,M. |
| EPI_ISL_14934538, EPI_ISL_14934540, EPI_ISL_14934543, EPI_ISL_14934550, EPI_ISL_14934574, EPI_ISL_14934587 | Department of Infectious Diseases, National Institute of Health Doutor Ricardo Jorge, Portugal (INSA) | Department of Infectious Diseases, National Institute of Health Doutor Ricardo Jorge, Portugal (INSA) | Isidro,J., Borges,V., Pinto,M., Sobral,D., Santos,J., Nunes,A., Mixao,V., Ferreira,R., Santos,D., Duarte,S., Vieira,L., Borrego,M.J., Nuncio,S., Lopes de Carvalho,I., Pelerito,A., Cordeiro,R. and Gomes,J.P. |
| EPI_ISL_14934620, EPI_ISL_14934622, EPI_ISL_14934623, EPI_ISL_14934624, EPI_ISL_14934626, EPI_ISL_14934627, EPI_ISL_14934628, EPI_ISL_14934629, EPI_ISL_14934630, EPI_ISL_14934632, EPI_ISL_14934633, EPI_ISL_14934637, EPI_ISL_14934638, EPI_ISL_14934639, EPI_ISL_14934640, EPI_ISL_14934641, EPI_ISL_14934642, EPI_ISL_14934643, EPI_ISL_14934644, EPI_ISL_14934645, EPI_ISL_14934646, EPI_ISL_14934647, EPI_ISL_14934648, EPI_ISL_14934649, EPI_ISL_14934650, EPI_ISL_14934651, EPI_ISL_14934652, EPI_ISL_14934654, EPI_ISL_14934655, EPI_ISL_14934656, EPI_ISL_14934657, EPI_ISL_14934658, EPI_ISL_14934659, EPI_ISL_14934661, EPI_ISL_14934662, EPI_ISL_14934663, EPI_ISL_14934664, EPI_ISL_14934665, EPI_ISL_14934666, EPI_ISL_14934667, EPI_ISL_14934668, EPI_ISL_14934670, EPI_ISL_14934671, EPI_ISL_14934672, EPI_ISL_14934673, EPI_ISL_14934674, EPI_ISL_14934675, EPI_ISL_14934677, EPI_ISL_14934678, EPI_ISL_14934679, EPI_ISL_14934680, EPI_ISL_14934681, EPI_ISL_14934682, EPI_ISL_14934683, EPI_ISL_14934684, EPI_ISL_14934685, EPI_ISL_14934686, EPI_ISL_14934687, EPI_ISL_14934688, EPI_ISL_14934689, EPI_ISL_14934690, EPI_ISL_14934691, EPI_ISL_14934692, EPI_ISL_14934693, EPI_ISL_14934694, EPI_ISL_14934695, EPI_ISL_14934696, EPI_ISL_14934697, EPI_ISL_14934698, EPI_ISL_14934699, EPI_ISL_14934700, EPI_ISL_14934701, EPI_ISL_14934702, EPI_ISL_14934703, EPI_ISL_14934704, EPI_ISL_14934705, EPI_ISL_14934706, EPI_ISL_14934707 | | | |
| see above | Research and Evaluation, UKHSA | Research and Evaluation, UKHSA | Groves,N., Osman,K.L., Lewandowski,K.S., Carter,D.P., Pullan,S.T., Myers,R., Vipond,R. and Chand,M. |
| EPI_ISL_14961089, EPI_ISL_14961090 | Public Health Authority of the Slovak Republic | Laboratory of Genomics and Bioinformatics, Comenius University Science Park | Tomáš Szemes, Edita Staroová, Elena Tichá, Lucia Ševíková, Terézia Vrabová, Tatiana Sedláková, Miroslav Böhmer, Jaroslav Budiš, Pavol Mišenko |
| EPI_ISL_14977306, EPI_ISL_14977307, EPI_ISL_14977308, EPI_ISL_14977309, EPI_ISL_14977310 | Environmental, Agricultural, and Occupational Health, University of Nebraska Medical Center | Environmental, Agricultural, and Occupational Health, University of Nebraska Medical Center | Tegomoh,B., Cross,S.T., Chapman,R.C., Bernhard,K., McCutchen,E.L., Fauver,J.R., Pratt,C.B., Warden,D.E., Iwen,P.C., Donahue,M. and Wiley,M.R. |
| EPI_ISL_14995206 | Pronto Socorro Municiplal de Cravinhos | Instituto Adolfo Lutz Strategic Laboratory | Claudio Tavares Sacchi, Karoline Rodrigues Campos, Ariadne Ferreira Amarante, Marlon Benedito Nascimento Santos, Alex Domingos Reis, Adriano Abbud, Adriana Bugno |
| EPI_ISL_14995578 | Hosp. Municipal de Ilhabela Gov. Mario Covas Jr. | Instituto Adolfo Lutz Strategic Laboratory | Claudio Tavares Sacchi, Karoline Rodrigues Campos, Ariadne Ferreira Amarante, Marlon Benedito Nascimento Santos, Alex Domingos Reis, Adriano Abbud, Adriana Bugno |
| EPI_ISL_14995579 | Secretaria Municipal de Saude de Feira de Santana | Instituto Adolfo Lutz Strategic Laboratory | Claudio Tavares Sacchi, Karoline Rodrigues Campos, Ariadne Ferreira Amarante, Marlon Benedito Nascimento Santos, Alex Domingos Reis, Adriano Abbud, Adriana Bugno |
| EPI_ISL_14995580 | UBS Alexander Fleming Simioni | Instituto Adolfo Lutz Strategic Laboratory | Claudio Tavares Sacchi, Karoline Rodrigues Campos, Ariadne Ferreira Amarante, Marlon Benedito Nascimento Santos, Alex Domingos Reis, Adriano Abbud, Adriana Bugno |
| EPI_ISL_14995582 | Hosp. Municipla. Dr. Jose de Carvalho Florence | Instituto Adolfo Lutz Strategic Laboratory | Claudio Tavares Sacchi, Karoline Rodrigues Campos, Ariadne Ferreira Amarante, Marlon Benedito Nascimento Santos, Alex Domingos Reis, Adriano Abbud, Adriana Bugno |
| EPI_ISL_14995585 | Pronto Socorro Municipal do Promorar | Instituto Adolfo Lutz Strategic Laboratory | Claudio Tavares Sacchi, Karoline Rodrigues Campos, Ariadne Ferreira Amarante, Marlon Benedito Nascimento Santos, Alex Domingos Reis, Adriano Abbud, Adriana Bugno |
| EPI_ISL_14995586 | UPA Centro | Instituto Adolfo Lutz Strategic Laboratory | Claudio Tavares Sacchi, Karoline Rodrigues Campos, Ariadne Ferreira Amarante, Marlon Benedito Nascimento Santos, Alex Domingos Reis, Adriano Abbud, Adriana Bugno |
| EPI_ISL_14995587 | Centro de Saude 24 horas | Instituto Adolfo Lutz Strategic Laboratory | Claudio Tavares Sacchi, Karoline Rodrigues Campos, Ariadne Ferreira Amarante, Marlon Benedito Nascimento Santos, Alex Domingos Reis, Adriano Abbud, Adriana Bugno |
| EPI_ISL_14995591 | Instituto de Infectologia Emilio Ribas | Instituto Adolfo Lutz Strategic Laboratory | Claudio Tavares Sacchi, Karoline Rodrigues Campos, Ariadne Ferreira Amarante, Marlon Benedito Nascimento Santos, Alex Domingos Reis, Adriano Abbud, Adriana Bugno |
| EPI_ISL_14995593 | SAE DST / Aids Ipiranga Jose Francisco Araujo | Instituto Adolfo Lutz Strategic Laboratory | Claudio Tavares Sacchi, Karoline Rodrigues Campos, Ariadne Ferreira Amarante, Marlon Benedito Nascimento Santos, Alex Domingos Reis, Adriano Abbud, Adriana Bugno |
| EPI_ISL_14995611 | UBS Horto Florestal | Instituto Adolfo Lutz Strategic Laboratory | Claudio Tavares Sacchi, Karoline Rodrigues Campos, Ariadne Ferreira Amarante, Marlon Benedito Nascimento Santos, Alex Domingos Reis, Adriano Abbud, Adriana Bugno |

| | | | |
|---|---|---|---|
| EPI_ISL_14995612 | Secretaria Minicipal de Saude de IRECE | Instituto Adolfo Lutz Strategic Laboratory | Claudio Tavares Sacchi, Karoline Rodrigues Campos, Ariadne Ferreira Amarante, Marlon Benedito Nascimento Santos, Alex Domingos Reis, Adriano Abbud, Adriana Bugno |
| EPI_ISL_14995619 | Hosp. Tereza de Lisieux | Instituto Adolfo Lutz Strategic Laboratory | Claudio Tavares Sacchi, Karoline Rodrigues Campos, Ariadne Ferreira Amarante, Marlon Benedito Nascimento Santos, Alex Domingos Reis, Adriano Abbud, Adriana Bugno |
| EPI_ISL_14995622 | UBS Parque Meia Lua | Instituto Adolfo Lutz Strategic Laboratory | Claudio Tavares Sacchi, Karoline Rodrigues Campos, Ariadne Ferreira Amarante, Marlon Benedito Nascimento Santos, Alex Domingos Reis, Adriano Abbud, Adriana Bugno |
| EPI_ISL_14995653 | Unidade Basica de Saude Vila Cristina | Instituto Adolfo Lutz Strategic Laboratory | Claudio Tavares Sacchi, Karoline Rodrigues Campos, Ariadne Ferreira Amarante, Marlon Benedito Nascimento Santos, Alex Domingos Reis, Adriano Abbud, Adriana Bugno |
| EPI_ISL_14995723 | Unidade Mista de Atendimento Infantil Carapicuiba | Instituto Adolfo Lutz Strategic Laboratory | Claudio Tavares Sacchi, Karoline Rodrigues Campos, Ariadne Ferreira Amarante, Marlon Benedito Nascimento Santos, Alex Domingos Reis, Adriano Abbud, Adriana Bugno |
| EPI_ISL_14995724 | Hosp. Carlos Chagas | Instituto Adolfo Lutz Strategic Laboratory | Claudio Tavares Sacchi, Karoline Rodrigues Campos, Ariadne Ferreira Amarante, Marlon Benedito Nascimento Santos, Alex Domingos Reis, Adriano Abbud, Adriana Bugno |
| EPI_ISL_14997063, EPI_ISL_14997067, EPI_ISL_14997068, EPI_ISL_14997069, EPI_ISL_14997070, EPI_ISL_14997071, EPI_ISL_14997072, EPI_ISL_14997073, EPI_ISL_14997074 | Laboratory Medicine, UW Virology | Laboratory Medicine, UW Virology | Sereewit,J., Xie,H., Roychoudhury,P. and Greninger,A.L. |
| EPI_ISL_15005641 | Chongqing Municipal Center for Disease Control and Prevention | Chongqing Municipal Center for Disease Control and Prevention | Sheng Ye, Yun Tang, Shuang Chen, Mingyue Wang, Zhangping Tan, Zhen Yu |
| EPI_ISL_15016105 | Centers for Disease Control & Prevention (CDC), Division of High Consequence Pathogens and Pathology (DHCPP-PRB) | Centers for Disease Control & Prevention (CDC), Division of High Consequence Pathogens and Pathology (DHCPP-PRB) | Gigante,C.M., Kubin,G., Zhao,H., Batra,D., Hetrick,E.E., Howard,D.T., Kovar,L., Seabolt,M.H., Weigand,M.R., Burroughs,M., Lee,J., Wilkins,K., McCollum,A., Hutson,C., Davidson,W., Rao,A., White,S.L. and Li,Y. |
| EPI_ISL_15016106 | Centers for Disease Control & Prevention (CDC), Division of High Consequence Pathogens and Pathology (DHCPP-PRB) | Centers for Disease Control & Prevention (CDC), Division of High Consequence Pathogens and Pathology (DHCPP-PRB) | Gigante,C.M., Goldoft,M., Zhao,H., Batra,D., Hetrick,E.E., Howard,D.T., Kovar,L., Seabolt,M.H., Weigand,M.R., Burroughs,M., Lee,J., Wilkins,K., McCollum,A., Hutson,C., Davidson,W., Rao,A., Holshue,M. and Li,Y. |
| EPI_ISL_15016107 | Centers for Disease Control & Prevention (CDC), Division of High Consequence Pathogens and Pathology (DHCPP-PRB) | Centers for Disease Control & Prevention (CDC), Division of High Consequence Pathogens and Pathology (DHCPP-PRB) | Gigante,C.M., Thomas,L., Zhao,H., Batra,D., Hetrick,E.E., Howard,D.T., Kovar,L., Seabolt,M.H., Weigand,M.R., Burroughs,M., Lee,J., Wilkins,K., McCollum,A., Hutson,C., Davidson,W., Rao,A., Dunn,J. and Li,Y. |
| EPI_ISL_15016109 | Centers for Disease Control & Prevention (CDC), Division of High Consequence Pathogens and Pathology (DHCPP-PRB) | Centers for Disease Control & Prevention (CDC), Division of High Consequence Pathogens and Pathology (DHCPP-PRB) | Gigante,C.M., Buttery,E., Zhao,H., Batra,D., Hetrick,E.E., Howard,D.T., Kovar,L., Seabolt,M.H., Weigand,M.R., Burroughs,M., Lee,J., Wilkins,K., McCollum,A., Hutson,C., Davidson,W., Rao,A., Raman,D. and Li,Y. |
| EPI_ISL_15016110 | Centers for Disease Control & Prevention (CDC), Division of High Consequence Pathogens and Pathology (DHCPP-PRB) | Centers for Disease Control & Prevention (CDC), Division of High Consequence Pathogens and Pathology (DHCPP-PRB) | Gigante,C.M., Kubin,G., Zhao,H., Batra,D., Hetrick,E.E., Howard,D.T., Kovar,L., Seabolt,M.H., Weigand,M.R., Burroughs,M., Lee,J., Wilkins,K., McCollum,A., Hutson,C., Davidson,W., Rao,A., White,S.L. and Li,Y. |
| EPI_ISL_15016111 | Centers for Disease Control & Prevention (CDC), Division of High Consequence Pathogens and Pathology (DHCPP-PRB) | Centers for Disease Control & Prevention (CDC), Division of High Consequence Pathogens and Pathology (DHCPP-PRB) | Gigante,C.M., Thomas,L., Zhao,H., Batra,D., Hetrick,E.E., Howard,D.T., Kovar,L., Seabolt,M.H., Weigand,M.R., Burroughs,M., Lee,J., Wilkins,K., McCollum,A., Hutson,C., Davidson,W., Rao,A., Dunn,J. and Li,Y. |
| EPI_ISL_15016112 | Centers for Disease Control & Prevention (CDC), Division of High Consequence Pathogens and Pathology (DHCPP-PRB) | Centers for Disease Control & Prevention (CDC), Division of High Consequence Pathogens and Pathology (DHCPP-PRB) | Gigante,C.M., Pettit,D., Zhao,H., Batra,D., Hetrick,E.E., Howard,D.T., Kovar,L., Seabolt,M.H., Weigand,M.R., Burroughs,M., Lee,J., Wilkins,K., McCollum,A., Hutson,C., Davidson,W., Rao,A., Deutsch-Feldman,M. and Li,Y. |
| EPI_ISL_15016113 | Centers for Disease Control & Prevention (CDC), Division of High Consequence Pathogens and Pathology (DHCPP-PRB) | Centers for Disease Control & Prevention (CDC), Division of High Consequence Pathogens and Pathology (DHCPP-PRB) | Gigante,C.M., Ghinai,I., Zhao,H., Batra,D., Hetrick,E.E., Howard,D.T., Kovar,L., Seabolt,M.H., Weigand,M.R., Burroughs,M., Lee,J., Wilkins,K., McCollum,A., Hutson,C., Davidson,W., Rao,A., Kerins,J. and Li,Y. |
| EPI_ISL_15016115 | Centers for Disease Control & Prevention (CDC), Division of High Consequence Pathogens and Pathology (DHCPP-PRB) | Centers for Disease Control & Prevention (CDC), Division of High Consequence Pathogens and Pathology (DHCPP-PRB) | Gigante,C.M., Thomas,L., Zhao,H., Batra,D., Hetrick,E.E., Howard,D.T., Kovar,L., Seabolt,M.H., Weigand,M.R., Burroughs,M., Lee,J., Wilkins,K., McCollum,A., Hutson,C., Davidson,W., Rao,A., Dunn,J. and Li,Y. |
| EPI_ISL_15016117 | Centers for Disease Control & Prevention (CDC), Division of High Consequence Pathogens and Pathology (DHCPP-PRB) | Centers for Disease Control & Prevention (CDC), Division of High Consequence Pathogens and Pathology (DHCPP-PRB) | Gigante,C.M., Epie,N., Zhao,H., Batra,D., Hetrick,E.E., Howard,D.T., Kovar,L., Seabolt,M.H., Weigand,M.R., Burroughs,M., Lee,J., Wilkins,K., McCollum,A., Hutson,C., Davidson,W., Rao,A., Perez,T. and Li,Y. |
| EPI_ISL_15016118, EPI_ISL_15016119, EPI_ISL_15016120 | Centers for Disease Control & Prevention (CDC), Division of High Consequence Pathogens and Pathology (DHCPP-PRB) | Centers for Disease Control & Prevention (CDC), Division of High Consequence Pathogens and Pathology (DHCPP-PRB) | Gigante,C.M., Acheampong,E., Zhao,H., Batra,D., Hetrick,E.E., Howard,D.T., Kovar,L., Seabolt,M.H., Weigand,M.R., Burroughs,M., Lee,J., Wilkins,K., McCollum,A., Hutson,C., Davidson,W., Rao,A., McDermott,D. and Li,Y. |
| EPI_ISL_15016121 | Centers for Disease Control & Prevention (CDC), Division of High Consequence Pathogens and Pathology (DHCPP-PRB) | Centers for Disease Control & Prevention (CDC), Division of High Consequence Pathogens and Pathology (DHCPP-PRB) | Gigante,C.M., Pettit,D., Zhao,H., Batra,D., Hetrick,E.E., Howard,D.T., Kovar,L., Seabolt,M.H., Weigand,M.R., Burroughs,M., Lee,J., Wilkins,K., McCollum,A., Hutson,C., Davidson,W., Rao,A., Deutsch-Feldman,M. and Li,Y. |
| EPI_ISL_15016122 | Centers for Disease Control & Prevention (CDC), Division of High Consequence Pathogens and Pathology (DHCPP-PRB) | Centers for Disease Control & Prevention (CDC), Division of High Consequence Pathogens and Pathology (DHCPP-PRB) | Gigante,C.M., Wang,X., Zhao,H., Batra,D., Hetrick,E.E., Howard,D.T., Kovar,L., Seabolt,M.H., Weigand,M.R., Burroughs,M., Lee,J., Wilkins,K., McCollum,A., Hutson,C., Davidson,W., Rao,A., Ostadkar,R. and Li,Y. |
| EPI_ISL_15016124 | Centers for Disease Control & Prevention (CDC), Division of High Consequence Pathogens and Pathology (DHCPP-PRB) | Centers for Disease Control & Prevention (CDC), Division of High Consequence Pathogens and Pathology (DHCPP-PRB) | Gigante,C.M., Haydel,D., Zhao,H., Batra,D., Hetrick,E.E., Howard,D.T., Kovar,L., Seabolt,M.H., Weigand,M.R., Burroughs,M., Lee,J., Wilkins,K., McCollum,A., Hutson,C., Davidson,W., Rao,A., Salinas,A. and Li,Y. |
| EPI_ISL_15016125, EPI_ISL_15016126 | Centers for Disease Control & Prevention (CDC), Division of High Consequence Pathogens and Pathology (DHCPP-PRB) | Centers for Disease Control & Prevention (CDC), Division of High Consequence Pathogens and Pathology (DHCPP-PRB) | Gigante,C.M., Pavlick,J., Zhao,H., Batra,D., Hetrick,E.E., Howard,D.T., Kovar,L., Seabolt,M.H., Weigand,M.R., Burroughs,M., Lee,J., Wilkins,K., McCollum,A., Hutson,C., Davidson,W., Rao,A., Parrott,T. and Li,Y. |
| EPI_ISL_15016127, EPI_ISL_15016128 | Centers for Disease Control & Prevention (CDC), Division of High Consequence Pathogens and Pathology (DHCPP-PRB) | Centers for Disease Control & Prevention (CDC), Division of High Consequence Pathogens and Pathology (DHCPP-PRB) | Gigante,C.M., Pearson,C., Zhao,H., Batra,D., Hetrick,E.E., Howard,D.T., Kovar,L., Seabolt,M.H., Weigand,M.R., Burroughs,M., Lee,J., Wilkins,K., McCollum,A., Hutson,C., Davidson,W., Rao,A., Maloney,M. and Li,Y. |
| EPI_ISL_15016129 | Centers for Disease Control & Prevention (CDC), Division of High Consequence Pathogens and Pathology (DHCPP-PRB) | Centers for Disease Control & Prevention (CDC), Division of High Consequence Pathogens and Pathology (DHCPP-PRB) | Gigante,C.M., Hauser,J.R., Zhao,H., Batra,D., Hetrick,E.E., Howard,D.T., Kovar,L., Seabolt,M.H., Weigand,M.R., Burroughs,M., Lee,J., Wilkins,K., McCollum,A., Hutson,C., Davidson,W., Rao,A., Mangla,A. and Li,Y. |
| EPI_ISL_15016130 | Centers for Disease Control & Prevention (CDC), Division of High Consequence Pathogens and Pathology (DHCPP-PRB) | Centers for Disease Control & Prevention (CDC), Division of High Consequence Pathogens and Pathology (DHCPP-PRB) | Gigante,C.M., Pearson,C., Zhao,H., Batra,D., Hetrick,E.E., Howard,D.T., Kovar,L., Seabolt,M.H., Weigand,M.R., Burroughs,M., Lee,J., Wilkins,K., McCollum,A., Hutson,C., Davidson,W., Rao,A., Maloney,M. and Li,Y. |
| EPI_ISL_15016131, EPI_ISL_15016132, EPI_ISL_15016133 | Centers for Disease Control & Prevention (CDC), Division of High Consequence Pathogens and Pathology (DHCPP-PRB) | Centers for Disease Control & Prevention (CDC), Division of High Consequence Pathogens and Pathology (DHCPP-PRB) | Gigante,C.M., Ventura,J., Zhao,H., Batra,D., Hetrick,E.E., Howard,D.T., Kovar,L., Seabolt,M.H., Weigand,M.R., Burroughs,M., Lee,J., Wilkins,K., McCollum,A., Hutson,C., Davidson,W., Rao,A., Nash,J. and Li,Y. |
| EPI_ISL_15023203 | Centers for Disease Control & Prevention (CDC), Division of High Consequence Pathogens and Pathology (DHCPP-PRB) | Centers for Disease Control & Prevention (CDC), Division of High Consequence Pathogens and Pathology (DHCPP-PRB) | Gigante,C.M., Pavlick,J., Zhao,H., Batra,D., Hetrick,E.E., Howard,D.T., Kovar,L., Seabolt,M.H., Weigand,M.R., Burroughs,M., Lee,J., Wilkins,K., McCollum,A., Hutson,C., Davidson,W., Rao,A., Parrott,T. and Li,Y. |
| EPI_ISL_15055820 | Sicilian Regional Laboratory - AOUP "P. Giaccone" - University of Palermo | Sicilian Regional Laboratory - AOUP "P. Giaccone" - University of Palermo | Fabio Tramuto, Carmelo Massimo Maida, Giulia Randazzo, Valeria Guzzetta, Walter Mazzucco, Giorgio Graziano, Vincenzo Restivo, Claudio Costantino, Francesco Vitale |
| EPI_ISL_15076130, EPI_ISL_15076131 | Environmental, Agricultural, and Occupational Health, University of Nebraska Medical Center | Environmental, Agricultural, and Occupational Health, University of Nebraska Medical Center | Chapman,R.C., Bernhard,K., McCutchen,E.L., Fauver,J.R., O'Dell,J.X., Mannell,M., Wiley,M.R. and Cross,S.T. |
| EPI_ISL_15076132, EPI_ISL_15076133, EPI_ISL_15076135, EPI_ISL_15076136, EPI_ISL_15076137, EPI_ISL_15076138, EPI_ISL_15076139, EPI_ISL_15076140, EPI_ISL_15076141, EPI_ISL_15076142, EPI_ISL_15076143, EPI_ISL_15076144, EPI_ISL_15076145, EPI_ISL_15076146, EPI_ISL_15076147, EPI_ISL_15076148, EPI_ISL_15076149, EPI_ISL_15076150, EPI_ISL_15076151, EPI_ISL_15076152, EPI_ISL_15076153, EPI_ISL_15076154, EPI_ISL_15076155, EPI_ISL_15076156, EPI_ISL_15076157, EPI_ISL_15076158, EPI_ISL_15076159, EPI_ISL_15076160, EPI_ISL_15076161, EPI_ISL_15076162, EPI_ISL_15076163, EPI_ISL_15076164, EPI_ISL_15076165 | | | |
| see above | Centre for Biological Threats, Highly Pathogenic Viruses, Robert Koch Institute | Centre for Biological Threats, Highly Pathogenic Viruses, Robert Koch Institute | Brinkmann,A., Kohl,C., Pape,K., Uddin,S., Schrick,L., Michel,J., Schaade,L. and Nitsche,A |
| EPI_ISL_15076180, EPI_ISL_15076183, EPI_ISL_15076188 | Department of Genetics, University of North Carolina at Chapel Hill | Department of Genetics, University of North Carolina at Chapel Hill | Deanhardt,B., Miller,M. and Wang,J.R. |
| EPI_ISL_15116266, EPI_ISL_15116267, EPI_ISL_15116268, EPI_ISL_15116269, EPI_ISL_15116270, EPI_ISL_15116271, EPI_ISL_15116272, EPI_ISL_15116273, EPI_ISL_15116274, EPI_ISL_15116275, EPI_ISL_15116276, EPI_ISL_15116277, EPI_ISL_15116278, EPI_ISL_15116279, EPI_ISL_15116280, EPI_ISL_15116281, | | | |

| | | | |
|---|---|---|---|
| EPI_ISL_15116282, EPI_ISL_15116284, EPI_ISL_15116285, EPI_ISL_15116287, EPI_ISL_15116288, EPI_ISL_15116289, EPI_ISL_15116290, EPI_ISL_15116291, EPI_ISL_15116292, EPI_ISL_15116293, EPI_ISL_15116294, EPI_ISL_15116295, EPI_ISL_15116296, EPI_ISL_15116297, EPI_ISL_15116298, EPI_ISL_15116299 | | | |
| see above | Centre for Biological Threats, Highly Pathogenic Viruses, Robert Koch Institute | Centre for Biological Threats, Highly Pathogenic Viruses, Robert Koch Institute | Brinkmann,A., Kohl,C., Pape,K., Schrick,L., Michel,J., Schaade,L. and Nitsche,A. |
| EPI_ISL_15120452, EPI_ISL_15120480, EPI_ISL_15120496 | Los Angeles County Public Health Laboratories | Los Angeles County Public Health Laboratories | P. Hemarajata et al. |
| EPI_ISL_15158315, EPI_ISL_15158316, EPI_ISL_15158336, EPI_ISL_15158341, EPI_ISL_15158358, EPI_ISL_15158360, EPI_ISL_15158361, EPI_ISL_15158363, EPI_ISL_15158364, EPI_ISL_15158367, EPI_ISL_15158369, EPI_ISL_15158371, EPI_ISL_15158374, EPI_ISL_15158384, EPI_ISL_15158385, EPI_ISL_15158387, EPI_ISL_15158390, EPI_ISL_15158391, EPI_ISL_15158394, EPI_ISL_15158395, EPI_ISL_15158398 | | | |
| see above | Molecular Biology, Microbiology, and Biochemistry, Southern Illinois University | Molecular Biology, Microbiology, and Biochemistry, Southern Illinois University | Gagnon,K.T. |
| EPI_ISL_15158402, EPI_ISL_15158406, EPI_ISL_15158407, EPI_ISL_15158408, EPI_ISL_15158409, EPI_ISL_15158414, EPI_ISL_15158416, EPI_ISL_15158417, EPI_ISL_15158418, EPI_ISL_15158419, EPI_ISL_15158421, EPI_ISL_15158423, EPI_ISL_15158428, EPI_ISL_15158430, EPI_ISL_15158433, EPI_ISL_15158435, EPI_ISL_15158437, EPI_ISL_15158446, EPI_ISL_15158447, EPI_ISL_15158448, EPI_ISL_15158450, EPI_ISL_15158455, EPI_ISL_15158458, EPI_ISL_15158459, EPI_ISL_15158460, EPI_ISL_15158461, EPI_ISL_15158462, EPI_ISL_15158463, EPI_ISL_15158464, EPI_ISL_15158466, EPI_ISL_15158467, EPI_ISL_15158468 | | | |
| see above | Research and Evaluation, UKHSA | Research and Evaluation, UKHSA | Groves,N., Osman,K.L., Lewandowski,K.S., Carter,D.P., Pullan,S.T., Myers,R., Vipond,R. and Chand,M. |
| EPI_ISL_15165603, EPI_ISL_15165604, EPI_ISL_15165608, EPI_ISL_15165614, EPI_ISL_15165618 | Centro de Desenvolvimento Científico e Tecnológico (CDCT), Centro Estadual de Vigilância em Saúde (CEVS) da Secretaria Estadual da Saúde (SES-RS) | Centro de Desenvolvimento Científico e Tecnológico (CDCT), Centro Estadual de Vigilância em Saúde (CEVS) da Secretaria Estadual da Saúde (SES-RS) | Richard Steiner Salvato, Fernanda Marques Godinho, Regina Bones Barcellos, Patricia Sesterheim, Amanda Pellenz Ruivo, Viviane Horn de Melo, Júlio Augusto Schroder |
| EPI_ISL_15199671, EPI_ISL_15199672, EPI_ISL_15199682, EPI_ISL_15199706, EPI_ISL_15199727, EPI_ISL_15199738, EPI_ISL_15199739, EPI_ISL_15199743, EPI_ISL_15199776, EPI_ISL_15199795, EPI_ISL_15199796 | | | |
| see above | Department of Infectious Diseases, National Institute of Health Doutor Ricardo Jorge, Portugal (INSA) | Department of Infectious Diseases, National Institute of Health Doutor Ricardo Jorge, Portugal (INSA) | Isidro,J., Borges,V., Pinto,M., Sobral,D., Santos,J., Nunes,A., Mixao,V., Ferreira,R., Santos,D., Duarte,S., Vieira,L., Borrego,M.J., Nuncio,S., Lopes de Carvalho,I., Pelerito,A., Cordeiro,R. and Gomes,J.P. |
| EPI_ISL_15247221 | Medical University of Vienna Center for Virology | Medical University of Vienna Center for Virology | Jeremy V. Camp, Monika Redlberger-Fritz, Stephan W. Aberle |
| EPI_ISL_15257669 | Public Health Agency of Canada, National Microbiology Laboratory | Public Health Agency of Canada, National Microbiology Laboratory | Duggan,A., Hole,D., Yadav,C., Knox,N., Haidl,E., Chapel,M., Tyler,A.D., Domselaar,G.V., Graham,M., Audet,J., Fernando,L., Hagan,M., Safronetz,D., Leung,A., Peters,G., Go,A., Kaplen,B., Antonation,K., Laminman,V., Jolly,G., Croxen,M., Deo,A., Dieu,P., Dong,X., Gill,K., Granger,D., Ferrato,C., Ikkurti,V., Kanji,J., Koleva,P., Li,V., Lloyd,C., Lynch,T., Ma,R., Pabbaraju,K., Rotich,S., Sergeant,H., Skitsko,T., Tipples,G., Thayer,J., Shideler,S. and Wong,A. |
| EPI_ISL_15257681, EPI_ISL_15257682, EPI_ISL_15257687 | Viral and Rickettsial Disease Laboratory, California Department of Public Health | Viral and Rickettsial Disease Laboratory, California Department of Public Health | Probert,W., Espinosa,A., Kath,C., Haw,M., O'Neil,R., Bell,J. and Hacker,J. |
| EPI_ISL_15263355 | Sicilian Regional Laboratory - AOUP "P. Giaccone" - University of Palermo | Sicilian Regional Laboratory - AOUP "P. Giaccone" - University of Palermo | Fabio Tramuto, Carmelo Massimo Maida, Giulia Randazzo, Valeria Guzzetta, Walter Mazzucco, Giorgio Graziano, Vincenzo Restivo, Claudio Costantino, Francesco Vitale |
| EPI_ISL_15269698, EPI_ISL_15269699, EPI_ISL_15269702, EPI_ISL_15269704 | Erasmus Medical Center Department of Virology | Erasmus Medical Center Department of Virology | Leonard Schuele, Bas Oude Munnink, Marjan Boter, Babette Weller, Babs Verstrepen, Richard Molenkamp, Janette Rahamat-Langendoen, Reina Sikkema, Marion Koopmans |
| EPI_ISL_15283583, EPI_ISL_15283585, EPI_ISL_15283587, EPI_ISL_15283589, EPI_ISL_15283590, EPI_ISL_15283592, EPI_ISL_15283594, EPI_ISL_15283596, EPI_ISL_15283598, EPI_ISL_15283600, EPI_ISL_15283601, EPI_ISL_15283603, EPI_ISL_15283605, EPI_ISL_15283607, EPI_ISL_15283608, EPI_ISL_15283610, EPI_ISL_15283611, EPI_ISL_15283612, EPI_ISL_15283614, EPI_ISL_15283616, EPI_ISL_15283618, EPI_ISL_15283620 | | | |
| see above | Centre for Biological Threats, Highly Pathogenic Viruses, Robert Koch Institute | Centre for Biological Threats, Highly Pathogenic Viruses, Robert Koch Institute | Brinkmann,A., Kohl,C., Pape,K., Schrick,L., Michel,J., Schaade,L. and Nitsche,A. |
| EPI_ISL_15292976, EPI_ISL_15292977, EPI_ISL_15292978, EPI_ISL_15292979, EPI_ISL_15292980, EPI_ISL_15292981, EPI_ISL_15292982, EPI_ISL_15292983, EPI_ISL_15292984, EPI_ISL_15292985, EPI_ISL_15292986, EPI_ISL_15292987, EPI_ISL_15292988, EPI_ISL_15292989, EPI_ISL_15292991, EPI_ISL_15292992, EPI_ISL_15292993, EPI_ISL_15292994, EPI_ISL_15292995, EPI_ISL_15292996, EPI_ISL_15292997, EPI_ISL_15292998, EPI_ISL_15292999, EPI_ISL_15293000, EPI_ISL_15293001, EPI_ISL_15293002, EPI_ISL_15293004, EPI_ISL_15293005, EPI_ISL_15293006, EPI_ISL_15293007, EPI_ISL_15293008, EPI_ISL_15293009 | | | |
| see above | Institute of Health Carlos III, Bioinformatics Unit | Institute of Health Carlos III, Bioinformatics Unit | Cuesta,I. |
| EPI_ISL_15293815 | National Institute for Viral Disease Control and Prevention (IVDC), Chinese Center for Disease Control and Prevention , Beijing, China | National Institute for Viral Disease Control and Prevention (IVDC), Chinese Center for Disease Control and Prevention , Beijing, China | Wenjie Tan, Changcheng Wu, Ruhan A, Wenling Wang, Roujian Lu, Li Zhao, Baoying Huang, Fei Ye, Wenbo Xu |
| EPI_ISL_15317143 | Centers for Disease Control & Prevention (CDC), Division of High Consequence Pathogens and Pathology (DHCPP-PRB) | Centers for Disease Control & Prevention (CDC), Division of High Consequence Pathogens and Pathology (DHCPP-PRB) | Gigante,C.M., Vang,K., Zhao,H., Batra,D., Hetrick,E.E., Howard,D.T., Kovar,L., Seabolt,M.H., Weigand,M.R., Burroughs,M., Lee,J., Wilkins,K., McCollum,A., Hutson,C., Davidson,W., Rao,A., Seely,K. and Li,Y. |
| EPI_ISL_15317144 | Centers for Disease Control & Prevention (CDC), Division of High Consequence Pathogens and Pathology (DHCPP-PRB) | Centers for Disease Control & Prevention (CDC), Division of High Consequence Pathogens and Pathology (DHCPP-PRB) | Gigante,C.M., Segaloff,H., Zhao,H., Batra,D., Hetrick,E.E., Howard,D.T., Kovar,L., Seabolt,M.H., Weigand,M.R., Burroughs,M., Lee,J., Wilkins,K., McCollum,A., Hutson,C., Davidson,W., Rao,A., Florek,K. and Li,Y. |
| EPI_ISL_15317145 | Centers for Disease Control & Prevention (CDC), Division of High Consequence Pathogens and Pathology (DHCPP-PRB) | Centers for Disease Control & Prevention (CDC), Division of High Consequence Pathogens and Pathology (DHCPP-PRB) | Gigante,C.M., Pavlick,J., Zhao,H., Batra,D., Hetrick,E.E., Howard,D.T., Kovar,L., Seabolt,M.H., Weigand,M.R., Burroughs,M., Lee,J., Wilkins,K., McCollum,A., Hutson,C., Davidson,W., Rao,A., Parrott,T. and Li,Y. |
| EPI_ISL_15317149 | Centers for Disease Control & Prevention (CDC), Division of High Consequence Pathogens and Pathology (DHCPP-PRB) | Centers for Disease Control & Prevention (CDC), Division of High Consequence Pathogens and Pathology (DHCPP-PRB) | Gigante,C.M., Xia,D., Zhao,H., Batra,D., Hetrick,E.E., Howard,D.T., Kovar,L., Seabolt,M.H., Weigand,M.R., Burroughs,M., Lee,J., Wilkins,K., McCollum,A., Hutson,C., Davidson,W., Rao,A., Plipat,N. and Li,Y. |
| EPI_ISL_15317150 | Centers for Disease Control & Prevention (CDC), Division of High Consequence Pathogens and Pathology (DHCPP-PRB) | Centers for Disease Control & Prevention (CDC), Division of High Consequence Pathogens and Pathology (DHCPP-PRB) | Gigante,C.M., Ruiz,V., Zhao,H., Batra,D., Hetrick,E.E., Howard,D.T., Kovar,L., Seabolt,M.H., Weigand,M.R., Burroughs,M., Lee,J., Wilkins,K., McCollum,A., Hutson,C., Davidson,W., Rao,A., Wang,J. and Li,Y. |
| EPI_ISL_15317151 | Centers for Disease Control & Prevention (CDC), Division of High Consequence Pathogens and Pathology (DHCPP-PRB) | Centers for Disease Control & Prevention (CDC), Division of High Consequence Pathogens and Pathology (DHCPP-PRB) | Gigante,C.M., Francis,D., Zhao,H., Batra,D., Hetrick,E.E., Howard,D.T., Kovar,L., Seabolt,M.H., Weigand,M.R., Burroughs,M., Lee,J., Wilkins,K., McCollum,A., Hutson,C., Davidson,W., Rao,A., Escobar,J. and Li,Y. |
| EPI_ISL_15317152 | Centers for Disease Control & Prevention (CDC), Division of High Consequence Pathogens and Pathology (DHCPP-PRB) | Centers for Disease Control & Prevention (CDC), Division of High Consequence Pathogens and Pathology (DHCPP-PRB) | Gigante,C.M., Johnson,S., Zhao,H., Batra,D., Hetrick,E.E., Howard,D.T., Kovar,L., Seabolt,M.H., Weigand,M.R., Burroughs,M., Lee,J., Wilkins,K., McCollum,A., Hutson,C., Davidson,W., Rao,A., Riner,D. and Li,Y. |
| EPI_ISL_15317153 | Centers for Disease Control & Prevention (CDC), Division of High Consequence Pathogens and Pathology (DHCPP-PRB) | Centers for Disease Control & Prevention (CDC), Division of High Consequence Pathogens and Pathology (DHCPP-PRB) | Gigante,C.M., Culbertson,M., Zhao,H., Batra,D., Hetrick,E.E., Howard,D.T., Kovar,L., Seabolt,M.H., Weigand,M.R., Burroughs,M., Lee,J., Wilkins,K., McCollum,A., Hutson,C., Davidson,W., Rao,A., Pope,B. and Li,Y. |
| EPI_ISL_15317154 | Centers for Disease Control & Prevention (CDC), Division of High Consequence Pathogens and Pathology (DHCPP-PRB) | Centers for Disease Control & Prevention (CDC), Division of High Consequence Pathogens and Pathology (DHCPP-PRB) | Gigante,C.M., Ventura,J., Zhao,H., Batra,D., Hetrick,E.E., Howard,D.T., Kovar,L., Seabolt,M.H., Weigand,M.R., Burroughs,M., Lee,J., Wilkins,K., McCollum,A., Hutson,C., Davidson,W., Rao,A., Nash,J. and Li,Y. |
| EPI_ISL_15317155 | Centers for Disease Control & Prevention (CDC), Division of High Consequence Pathogens and Pathology (DHCPP-PRB) | Centers for Disease Control & Prevention (CDC), Division of High Consequence Pathogens and Pathology (DHCPP-PRB) | Gigante,C.M., Francis,D., Zhao,H., Batra,D., Hetrick,E.E., Howard,D.T., Kovar,L., Seabolt,M.H., Weigand,M.R., Burroughs,M., Lee,J., Wilkins,K., McCollum,A., Hutson,C., Davidson,W., Rao,A., Escobar,J. and Li,Y. |
| EPI_ISL_15317157 | Centers for Disease Control & Prevention (CDC), Division of High Consequence Pathogens and Pathology (DHCPP-PRB) | Centers for Disease Control & Prevention (CDC), Division of High Consequence Pathogens and Pathology (DHCPP-PRB) | Gigante,C.M., Pavlick,J., Zhao,H., Batra,D., Hetrick,E.E., Howard,D.T., Kovar,L., Seabolt,M.H., Weigand,M.R., Burroughs,M., Lee,J., Wilkins,K., McCollum,A., Hutson,C., Davidson,W., Rao,A., Parrott,T. and Li,Y. |
| EPI_ISL_15317158, EPI_ISL_15317159, EPI_ISL_15317160 | Centers for Disease Control & Prevention (CDC), Division of High Consequence Pathogens and Pathology (DHCPP-PRB) | Centers for Disease Control & Prevention (CDC), Division of High Consequence Pathogens and Pathology (DHCPP-PRB) | Gigante,C.M., Ventura,J., Zhao,H., Batra,D., Hetrick,E.E., Howard,D.T., Kovar,L., Seabolt,M.H., Weigand,M.R., Burroughs,M., Lee,J., Wilkins,K., McCollum,A., Hutson,C., Davidson,W., Rao,A., Nash,J. and Li,Y. |
| EPI_ISL_15332321 | Centers for Disease Control & Prevention (CDC), Division of High Consequence Pathogens and Pathology (DHCPP-PRB) | Centers for Disease Control & Prevention (CDC), Division of High Consequence Pathogens and Pathology (DHCPP-PRB) | Gigante,C.M., Carlson,C., Zhao,H., Batra,D., Hetrick,E.E., Howard,D.T., Kovar,L., Seabolt,M.H., Knipe,K., Burroughs,S., Lee,J., Wilkins,K., McCollum,A., Hutson,C., Davidson,W., Rao,A., Southern,T. and Li,Y. |
| EPI_ISL_15332324 | Centers for Disease Control & Prevention (CDC), Division of High Consequence Pathogens and Pathology (DHCPP-PRB) | Centers for Disease Control & Prevention (CDC), Division of High Consequence Pathogens and Pathology (DHCPP-PRB) | Gigante,C.M., Xia,D., Zhao,H., Batra,D., Hetrick,E.E., Howard,D.T., Kovar,L., Seabolt,M.H., Knipe,K., Burroughs,S., Lee,J., Wilkins,K., McCollum,A., Hutson,C., Davidson,W., Rao,A., Plipat,N. and Li,Y. |
| EPI_ISL_15332325 | Centers for Disease Control & Prevention (CDC), Division of High Consequence Pathogens and Pathology (DHCPP-PRB) | Centers for Disease Control & Prevention (CDC), Division of High Consequence Pathogens and Pathology (DHCPP-PRB) | Gigante,C.M., Pettit,D., Zhao,H., Batra,D., Hetrick,E.E., Howard,D.T., Kovar,L., Seabolt,M.H., Knipe,K., Burroughs,S., Lee,J., Wilkins,K., McCollum,A., Hutson,C., Davidson,W., Rao,A., Deutsch-Feldman,M. and Li,Y. |
| EPI_ISL_15332327 | Centers for Disease Control & Prevention (CDC), Division of | Centers for Disease Control & Prevention (CDC), Division of | Gigante,C.M., Acheampong,E., Zhao,H., Batra,D., Hetrick,E.E., Howard,D.T., Kovar,L., Seabolt,M.H., Knipe,K., Burroughs,S., Lee,J., Wilkins,K., |

| | | | |
|---|---|---|---|
| | High Consequence Pathogens and Pathology (DHCPP-PRB) | High Consequence Pathogens and Pathology (DHCPP-PRB) | McCollum,A., Hutson,C., Davidson,W., Rao,A., McDermott,D. and Li,Y. |
| EPI_ISL_15332328 | Centers for Disease Control & Prevention (CDC), Division of High Consequence Pathogens and Pathology (DHCPP-PRB) | Centers for Disease Control & Prevention (CDC), Division of High Consequence Pathogens and Pathology (DHCPP-PRB) | Gigante,C.M., Pettit,D., Zhao,H., Batra,D., Hetrick,E.E., Howard,D.T., Kovar,L., Seabolt,M.H., Knipe,K., Burroughs,S., Lee,J., Wilkins,K., McCollum,A., Hutson,C., Davidson,W., Rao,A., Deutsch-Feldman,M. and Li,Y. |
| EPI_ISL_15332329, EPI_ISL_15332330 | Centers for Disease Control & Prevention (CDC), Division of High Consequence Pathogens and Pathology (DHCPP-PRB) | Centers for Disease Control & Prevention (CDC), Division of High Consequence Pathogens and Pathology (DHCPP-PRB) | Gigante,C.M., Myers,R., Zhao,H., Batra,D., Hetrick,E.E., Howard,D.T., Kovar,L., Seabolt,M.H., Knipe,K., Burroughs,S., Lee,J., Wilkins,K., McCollum,A., Hutson,C., Davidson,W., Rao,A., Blythe,D. and Li,Y. |
| EPI_ISL_15332331 | Centers for Disease Control & Prevention (CDC), Division of High Consequence Pathogens and Pathology (DHCPP-PRB) | Centers for Disease Control & Prevention (CDC), Division of High Consequence Pathogens and Pathology (DHCPP-PRB) | Gigante,C.M., Pavlick,J., Zhao,H., Batra,D., Hetrick,E.E., Howard,D.T., Kovar,L., Seabolt,M.H., Knipe,K., Burroughs,S., Lee,J., Wilkins,K., McCollum,A., Hutson,C., Davidson,W., Rao,A., Parrott,T. and Li,Y. |
| EPI_ISL_15332332, EPI_ISL_15332333 | Centers for Disease Control & Prevention (CDC), Division of High Consequence Pathogens and Pathology (DHCPP-PRB) | Centers for Disease Control & Prevention (CDC), Division of High Consequence Pathogens and Pathology (DHCPP-PRB) | Gigante,C.M., Lee,P., Zhao,H., Batra,D., Hetrick,E.E., Howard,D.T., Kovar,L., Seabolt,M.H., Knipe,K., Burroughs,S., Lee,J., Wilkins,K., McCollum,A., Hutson,C., Davidson,W., Rao,A., Stanek,D. and Li,Y. |
| EPI_ISL_15332334 | Centers for Disease Control & Prevention (CDC), Division of High Consequence Pathogens and Pathology (DHCPP-PRB) | Centers for Disease Control & Prevention (CDC), Division of High Consequence Pathogens and Pathology (DHCPP-PRB) | Gigante,C.M., Hauser,J.R., Zhao,H., Batra,D., Hetrick,E.E., Howard,D.T., Kovar,L., Seabolt,M.H., Knipe,K., Burroughs,S., Lee,J., Wilkins,K., McCollum,A., Hutson,C., Davidson,W., Rao,A., Mangla,A. and Li,Y. |
| EPI_ISL_15332335 | Centers for Disease Control & Prevention (CDC), Division of High Consequence Pathogens and Pathology (DHCPP-PRB) | Centers for Disease Control & Prevention (CDC), Division of High Consequence Pathogens and Pathology (DHCPP-PRB) | Gigante,C.M., Ventura,J., Zhao,H., Batra,D., Hetrick,E.E., Howard,D.T., Kovar,L., Seabolt,M.H., Knipe,K., Burroughs,S., Lee,J., Wilkins,K., McCollum,A., Hutson,C., Davidson,W., Rao,A., Nash,J. and Li,Y. |
| EPI_ISL_15332336, EPI_ISL_15332337 | Environmental, Agricultural, and Occupational Health, University of Nebraska Medical Center | Environmental, Agricultural, and Occupational Health, University of Nebraska Medical Center | Tegomoh,B., Cross,S.T., Chapman,R.C., Bernhard,K., McCutchen,E.L., Fauver,J.R., Pratt,C.B., Warden,D.E., Iwen,P.C., Donahue,M. and Wiley,M.R. |
| EPI_ISL_15332339, EPI_ISL_15332340 | Environmental, Agricultural, and Occupational Health, University of Nebraska Medical Center | Environmental, Agricultural, and Occupational Health, University of Nebraska Medical Center | Chapman,R.C., Bernhard,K., McCutchen,E.L., Fauver,J.R., O'Dell,J.X., Mannell,M., Wiley,M.R. and Cross,S.T. |
| EPI_ISL_15352249, EPI_ISL_15352251, EPI_ISL_15352252, EPI_ISL_15352254, EPI_ISL_15352256, EPI_ISL_15352258, EPI_ISL_15352259, EPI_ISL_15352262, EPI_ISL_15352263, EPI_ISL_15352265, EPI_ISL_15352267, EPI_ISL_15352268, EPI_ISL_15352271, EPI_ISL_15352276, EPI_ISL_15352278, EPI_ISL_15352279, EPI_ISL_15352280, EPI_ISL_15352282, EPI_ISL_15352283, EPI_ISL_15352284, EPI_ISL_15352285, EPI_ISL_15352287, EPI_ISL_15352289, EPI_ISL_15352291, EPI_ISL_15352297, EPI_ISL_15352299, EPI_ISL_15352300, EPI_ISL_15352302, EPI_ISL_15352305, EPI_ISL_15352307, EPI_ISL_15352308, EPI_ISL_15352310, EPI_ISL_15352311, EPI_ISL_15352312, EPI_ISL_15352314 | | | |
| see above | Centre for Biological Threats, Highly Pathogenic Viruses, Robert Koch Institute | Centre for Biological Threats, Highly Pathogenic Viruses, Robert Koch Institute | Brinkmann,A., Kohl,C., Pape,K., Schrick,L., Michel,J., Schaade,L. and Nitsche,A |
| EPI_ISL_15367957, EPI_ISL_15367958, EPI_ISL_15367959, EPI_ISL_15367960, EPI_ISL_15367961, EPI_ISL_15367962, EPI_ISL_15367963, EPI_ISL_15367964, EPI_ISL_15367965, EPI_ISL_15367966, EPI_ISL_15367967, EPI_ISL_15367969, EPI_ISL_15367970, EPI_ISL_15367971, EPI_ISL_15367972, EPI_ISL_15367973, EPI_ISL_15367974, EPI_ISL_15367975, EPI_ISL_15367976, EPI_ISL_15367977, EPI_ISL_15367978, EPI_ISL_15367979, EPI_ISL_15367980, EPI_ISL_15367981, EPI_ISL_15367983, EPI_ISL_15367984, EPI_ISL_15367985, EPI_ISL_15367986, EPI_ISL_15367987, EPI_ISL_15367988, EPI_ISL_15367989, EPI_ISL_15367990, EPI_ISL_15367991, EPI_ISL_15367992, EPI_ISL_15367993, EPI_ISL_15367994, EPI_ISL_15367995, EPI_ISL_15367996, EPI_ISL_15367997, EPI_ISL_15367998, EPI_ISL_15367999, EPI_ISL_15368000, EPI_ISL_15368001, EPI_ISL_15368002, EPI_ISL_15368003, EPI_ISL_15368004, EPI_ISL_15368005, EPI_ISL_15368006, EPI_ISL_15368007, EPI_ISL_15368008, EPI_ISL_15368009, EPI_ISL_15368010, EPI_ISL_15368011 | | | |
| see above | Laboratory Medicine, UW Virology | Laboratory Medicine, UW Virology | Sereewit,J., Xie,H., Roychoudhury,P. and Greninger,A.L. |
| EPI_ISL_15370026, EPI_ISL_15370027, EPI_ISL_15370058, EPI_ISL_15370059, EPI_ISL_15370060 | Laboratory for Diagnostics of Zoonoses and WHO Centre, Institute of Microbiology and Immunology, Faculty of Medicine, University of Ljubljana | Laboratory for Diagnostics of Zoonoses and WHO Centre, Institute of Microbiology and Immunology, Faculty of Medicine, University of Ljubljana | Zakotnik,S., Vlaj,D., Suljic,A., Zorec,T.M., Korva,M., Poljak,M. and Avsic Zupanc,T. |
| EPI_ISL_15380492 | Eastwood Medical City | Molecular Biology Laboratory, Research Institute for Tropical Medicine | Samantha Louise P. Bado, Niquitta B. Galap, Bea C. Mateo, Chelsea Mae M. Reyes, Amalea Dulcene Nicolasora, Miguel Francisco B. Abulencia, Francisco Gerardo M. Polotan on behalf of the Research Institute for Tropical Medicine |
| EPI_ISL_15384398, EPI_ISL_15384399, EPI_ISL_15384400, EPI_ISL_15384402, EPI_ISL_15384403, EPI_ISL_15384404, EPI_ISL_15384405, EPI_ISL_15384406, EPI_ISL_15384407, EPI_ISL_15384408 | Laboratory Medicine, UW Virology | Laboratory Medicine, UW Virology | Sereewit,J., Xie,H., Roychoudhury,P. and Greninger,A.L. |
| EPI_ISL_15390495, EPI_ISL_15390496, EPI_ISL_15390497, EPI_ISL_15390498, EPI_ISL_15390499, EPI_ISL_15390500, EPI_ISL_15390501 | Centre for Biological Threats, Highly Pathogenic Viruses, Robert Koch Institute | Centre for Biological Threats, Highly Pathogenic Viruses, Robert Koch Institute | Brinkmann,A., Kohl,C., Pape,K., Uddin,S., Schrick,L., Michel,J., Schaade,L. and Nitsche,A. |
| EPI_ISL_15390502 | Division of High-Consequence Pathogens and Pathology (DHCPP-PRB), CDC | Division of High-Consequence Pathogens and Pathology (DHCPP-PRB), CDC | Gigante,C.M., Pavlick,J., Zhao,H., Batra,D., Hetrick,E.E., Howard,D.T., Kovar,L., Seabolt,M.H., Weigand,M.R., Burroughs,M., Lee,J., Wilkins,K., McCollum,A., Hutson,C., Davidson,W., Rao,A., Parrott,T. and Li,Y. |
| EPI_ISL_15419131 | Instituto de Infectologia Emilio Ribas | Instituto Adolfo Lutz Strategic Laboratory | Claudio Tavares Sacchi, Karoline Rodrigues Campos, Ariadne Ferreira Amarante, Marlon Benedito Nascimento Santos, Adriano Abbud, Adriana Bugno |
| EPI_ISL_15419133 | UBS II COHAB Presidente Prudente | Instituto Adolfo Lutz Strategic Laboratory | Claudio Tavares Sacchi, Karoline Rodrigues Campos, Ariadne Ferreira Amarante, Marlon Benedito Nascimento Santos, Adriano Abbud, Adriana Bugno |
| EPI_ISL_15419134 | CTA Centro de Testagem e Aconselhamento de Caieiras | Instituto Adolfo Lutz Strategic Laboratory | Claudio Tavares Sacchi, Karoline Rodrigues Campos, Ariadne Ferreira Amarante, Marlon Benedito Nascimento Santos, Adriano Abbud, Adriana Bugno |
| EPI_ISL_15419135 | SAE DST AIDS Cidade Dutra | Instituto Adolfo Lutz Strategic Laboratory | Claudio Tavares Sacchi, Karoline Rodrigues Campos, Ariadne Ferreira Amarante, Marlon Benedito Nascimento Santos, Adriano Abbud, Adriana Bugno |
| EPI_ISL_15419136 | UBS J Nordeste | Instituto Adolfo Lutz Strategic Laboratory | Claudio Tavares Sacchi, Karoline Rodrigues Campos, Ariadne Ferreira Amarante, Marlon Benedito Nascimento Santos, Adriano Abbud, Adriana Bugno |
| EPI_ISL_15419137 | CTA Centro de Testagem e Aconselhamento Favo de Mel | Instituto Adolfo Lutz Strategic Laboratory | Claudio Tavares Sacchi, Karoline Rodrigues Campos, Ariadne Ferreira Amarante, Marlon Benedito Nascimento Santos, Adriano Abbud, Adriana Bugno |
| EPI_ISL_15419138 | Vigilancia Epidemiologica e Controle de Vetores de Pirassununga | Instituto Adolfo Lutz Strategic Laboratory | Claudio Tavares Sacchi, Karoline Rodrigues Campos, Ariadne Ferreira Amarante, Marlon Benedito Nascimento Santos, Adriano Abbud, Adriana Bugno |
| EPI_ISL_15419140 | Secretaria Municipal de Saude de Batatais SP | Instituto Adolfo Lutz Strategic Laboratory | Claudio Tavares Sacchi, Karoline Rodrigues Campos, Ariadne Ferreira Amarante, Marlon Benedito Nascimento Santos, Adriano Abbud, Adriana Bugno |
| EPI_ISL_15419141 | UBS J Nordeste | Instituto Adolfo Lutz Strategic Laboratory | Claudio Tavares Sacchi, Karoline Rodrigues Campos, Ariadne Ferreira Amarante, Marlon Benedito Nascimento Santos, Adriano Abbud, Adriana Bugno |
| EPI_ISL_15419142 | Instituto de Infectologia Emilio Ribas | Instituto Adolfo Lutz Strategic Laboratory | Claudio Tavares Sacchi, Karoline Rodrigues Campos, Ariadne Ferreira Amarante, Marlon Benedito Nascimento Santos, Adriano Abbud, Adriana Bugno |
| EPI_ISL_15419143 | Santa Casa de Barretos | Instituto Adolfo Lutz Strategic Laboratory | Claudio Tavares Sacchi, Karoline Rodrigues Campos, Ariadne Ferreira Amarante, Marlon Benedito Nascimento Santos, Adriano Abbud, Adriana Bugno |
| EPI_ISL_15419144 | Hospital Vera Cruz | Instituto Adolfo Lutz Strategic Laboratory | Claudio Tavares Sacchi, Karoline Rodrigues Campos, Ariadne Ferreira Amarante, Marlon Benedito Nascimento Santos, Adriano Abbud, Adriana Bugno |
| EPI_ISL_15419145 | NotreDame Intermedica Saude | Instituto Adolfo Lutz Strategic Laboratory | Claudio Tavares Sacchi, Karoline Rodrigues Campos, Ariadne Ferreira Amarante, Marlon Benedito Nascimento Santos, Adriano Abbud, Adriana Bugno |
| EPI_ISL_15419146 | Hospital e Maternidade Santa Maria Cruz Azul | Instituto Adolfo Lutz Strategic Laboratory | Claudio Tavares Sacchi, Karoline Rodrigues Campos, Ariadne Ferreira Amarante, Marlon Benedito Nascimento Santos, Adriano Abbud, Adriana Bugno |
| EPI_ISL_15419147 | Pronto Socorro Central de Diadema | Instituto Adolfo Lutz Strategic Laboratory | Claudio Tavares Sacchi, Karoline Rodrigues Campos, Ariadne Ferreira Amarante, Marlon Benedito Nascimento Santos, Adriano Abbud, Adriana Bugno |
| EPI_ISL_15419148 | NotreDame Intermedica Saude Santo Andre | Instituto Adolfo Lutz Strategic Laboratory | Claudio Tavares Sacchi, Karoline Rodrigues Campos, Ariadne Ferreira Amarante, Marlon Benedito Nascimento Santos, Adriano Abbud, Adriana Bugno |
| EPI_ISL_15419151 | UPA 24H Brotas | Instituto Adolfo Lutz Strategic Laboratory | Claudio Tavares Sacchi, Karoline Rodrigues Campos, Ariadne Ferreira Amarante, Marlon Benedito Nascimento Santos, Adriano Abbud, Adriana Bugno |
| EPI_ISL_15419152 | Hospital Nossa Senhora de Lourdes | Instituto Adolfo Lutz Strategic Laboratory | Claudio Tavares Sacchi, Karoline Rodrigues Campos, Ariadne Ferreira Amarante, Marlon Benedito Nascimento Santos, Adriano Abbud, Adriana Bugno |
| EPI_ISL_15419153 | AMA Paraisopolis | Instituto Adolfo Lutz Strategic Laboratory | Claudio Tavares Sacchi, Karoline Rodrigues Campos, Ariadne Ferreira Amarante, Marlon Benedito Nascimento Santos, Adriano Abbud, Adriana Bugno |
| EPI_ISL_15419154 | Pronto Atendimento Infantil e Central de Quimioterapia de Sao Jose do Rio Preto | Instituto Adolfo Lutz Strategic Laboratory | Claudio Tavares Sacchi, Karoline Rodrigues Campos, Ariadne Ferreira Amarante, Marlon Benedito Nascimento Santos, Adriano Abbud, Adriana Bugno |
| EPI_ISL_15419155 | Santa Casa de Atibaia Pro Saude | Instituto Adolfo Lutz Strategic Laboratory | Claudio Tavares Sacchi, Karoline Rodrigues Campos, Ariadne Ferreira Amarante, Marlon Benedito Nascimento Santos, Adriano Abbud, Adriana Bugno |
| EPI_ISL_15419156 | UPA Vila Mariana | Instituto Adolfo Lutz Strategic Laboratory | Claudio Tavares Sacchi, Karoline Rodrigues Campos, Ariadne Ferreira Amarante, Marlon Benedito Nascimento Santos, Adriano Abbud, Adriana Bugno |
| EPI_ISL_15419157 | Secretaria de Saude de Mogi das Cruzes | Instituto Adolfo Lutz Strategic Laboratory | Claudio Tavares Sacchi, Karoline Rodrigues Campos, Ariadne Ferreira Amarante, Marlon Benedito Nascimento Santos, Adriano Abbud, Adriana Bugno |

| | | | |
|---|---|---|---|
| EPI_ISL_15419158 | Unidade Mista de saude Mariano Gayoso Castelo Branco | Instituto Adolfo Lutz Strategic Laboratory | Claudio Tavares Sacchi, Karoline Rodrigues Campos, Ariadne Ferreira Amarante, Marlon Benedito Nascimento Santos, Adriano Abbud, Adriana Bugno |
| EPI_ISL_15419161 | Centro de Saude I Albertino Affonso Jaboticabal | Instituto Adolfo Lutz Strategic Laboratory | Claudio Tavares Sacchi, Karoline Rodrigues Campos, Ariadne Ferreira Amarante, Marlon Benedito Nascimento Santos, Adriano Abbud, Adriana Bugno |
| EPI_ISL_15419162 | SAE DST AIDS M Boi Mirim Servico de Atencao Especializada | Instituto Adolfo Lutz Strategic Laboratory | Claudio Tavares Sacchi, Karoline Rodrigues Campos, Ariadne Ferreira Amarante, Marlon Benedito Nascimento Santos, Adriano Abbud, Adriana Bugno |
| EPI_ISL_15433343, EPI_ISL_15433348 | Laboratory Medicine, UW Virology | Laboratory Medicine, UW Virology | Sereewit,J., Xie,H., Roychoudhury,P. and Greninger,A.L. |
